# Genomic Dimensionality Bounds Mixed-Model Association Power, Fine-Mapping Resolution, and Genomic Prediction Reliability

**DOI:** 10.64898/2026.06.02.729628

**Authors:** Jicai Jiang

## Abstract

Mixed-model genome-wide association studies (GWAS) behave differently in livestock than in humans, yet a unified explanation is lacking. Analyses using the full genomic relationship matrix (full-GRM; from genome-wide SNPs) yield only a few significant peaks even with hundreds of thousands of animals, whereas leave-one-chromosome-out (LOCO), numerator-relationship-matrix, and sparse-GRM approaches report many broad associations over similar data. Here we develop a framework that traces these behaviors to the low effective genomic dimensionality, *M*_*e*_, of small-*N*_*e*_ populations. Starting from the mixed-model association statistic, we derive the per-SNP non-centrality parameter under full-GRM testing and show that its sample-size dependence is fully captured by a sigmoid sum *S*(*N*) over LD-matrix eigenmodes. *S*(*N*) grows concavely with *N* toward a practical ceiling *M*_*e*_, from which the framework predicts a full-GRM detection floor *q*_min_ ≈ 30*h*^2^*/M*_*e*_ on per-SNP proportion of phenotypic variance explained at 50% power (e.g., ~0.09% for cattle at *h*^2^ = 0.3), and a fine-mapping resolution limit through both *M*_*e*_ and 4*N*_*e*_-scaled LD decay. LOCO bypasses the full-GRM ceiling but detects LD-aggregated block-level signals rather than SNP-level excess effects, explaining its inflation in livestock and agreement with full-GRM in humans. The framework is supported by analyses of livestock chip panels, coalescent eigenvalue spectra, and phenotype simulations. The same *S*(*N*) sets the in-sample GBLUP reliability and bounds the out-of-sample reliability, 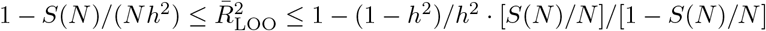, explaining why genomic prediction is comparatively easy while SNP-level mapping and fine-mapping remain difficult in livestock (vice versa in humans). For livestock GWAS aimed at SNP-level interpretation (e.g., candidate-gene prioritization, fine-mapping, or molecular-QTL colocalization), the framework supports full-GRM methods as the appropriate default.

**Article Summary:** Genome-wide association methods that largely agree in humans can give strikingly different results in livestock, complicating their use and interpretation for animal breeding. This study develops a framework, derived from the association test statistic, that traces this divergence to one cause. The small effective population size of livestock explains why some methods detect few signals even in huge datasets, why others report many broad associations, why livestock differ from humans, how precisely causal variants can be mapped, and why prediction stays comparatively easy. The framework predicts these outcomes quantitatively, with explicit formulas. It guides method choice and sets realistic expectations.

## Introduction

Mixed-model association methods are the standard approach for genome-wide association studies (GWAS) of complex traits. By modeling polygenic background as a random-effect term with covariance proportional to a genomic relationship matrix (GRM), these methods account for population structure and relatedness while testing individual SNP effects (Yu *et al*. 2006; Kang *et al*. 2008). Subsequent methodological developments have improved computational efficiency and statistical modeling. GEMMA (Zhou and Stephens 2012) made exact mixed-model GWAS computationally efficient by reusing a one-time eigendecomposition of the GRM across genome-wide SNPs. GCTA (Yang *et al*. 2014) applied variance components estimated once from the null model to all SNPs being tested, avoiding repeated optimization. GRAMMAR-Gamma (Svishcheva *et al*. 2012) approximates mixed-model association testing by applying a single genome-wide scalar correction to per-SNP score statistics computed on mixed-model residuals. BOLT-LMM (Loh *et al*. 2015) replaced the infinitesimal polygenic prior with Bayesian mixture-of-normals priors for SNP effects, increasing power for non-infinitesimal genetic architectures, and scaled to biobank-sized datasets. fastGWA (Jiang *et al*. 2019b) used a sparse GRM for computational efficiency. SAIGE (Zhou *et al*. 2018) extended mixed-model association to binary traits with case-control imbalance. Orthogonal to these developments, the leave-one-chromosome-out (LOCO) strategy, which excludes SNPs on the focal chromosome from the GRM to avoid proximal contamination, has been adopted in many GWAS software tools such as GCTA, BOLT-LMM, and SAIGE. Although originally developed for human genetics, these methods are now applied across species, including livestock, companion animals, and plants. GWAS summary statistics produced by these methods underpin a range of downstream analyses, including genetic fine-mapping (Schaid *et al*. 2018), colocalization (Giambartolomei *et al*. 2014), Mendelian randomization (Davey Smith and Hemani 2014), and partitioning heritability by functional annotations (Finucane *et al*. 2015), so the accuracy and interpretability of association results matter far beyond the GWAS itself.

A key observation in livestock genomics is that **moderate-density chip panels appear sufficient** both for genomic prediction and for GRM construction in association analysis. In dairy cattle, prediction accuracy using ~60,000 chip SNPs closely matches that from high-density chips or whole-genome sequence data (VanRaden *et al*. 2017), and association results are broadly concordant across GRM marker densities once the panel exceeds ~30,000 SNPs (Wang *et al*. 2025). This is consistent with the limited effective dimensionality of genomic information in livestock. Only a limited number of eigenvalues of the GRM are non-negligible (Pocrnic *et al*. 2016), corresponding to approximately *M*_*e*_ = 4*N*_*e*_*L* independent chromosome segments (Sved 1971; Stam 1980; Goddard 2009), where *N*_*e*_ is the effective population size and *L* is the genome length in Morgans. For livestock species with *N*_*e*_ ≈ 50–200, *M*_*e*_ is on the order of 10^3^–10^4^, smaller than the number of SNPs on a standard genotyping chip.

Against this background, two empirical patterns in livestock GWAS motivate the present study. First, **different mixed-model approaches produce strikingly different association results in large livestock GWAS**. Under LOCO, large dairy cattle GWAS can show severe genomic inflation and significant associations spanning entire chromosomes (Jiang *et al*. 2019a); GWAS using a pedigree-based relationship matrix shows a similar though less severe inflation pattern (Jiang *et al*. 2019a). In contrast, full-GRM mixed-model analysis consistently identifies only a small number of association peaks, even at very large sample sizes. A full-GRM mixed-model GWAS of 1.16 million genotyped Holstein cattle detected a limited number of significant loci (Jiang *et al*. 2022). A similar full-GRM analysis of sequence variants in ~50,000 Holstein bulls, each with highly reliable deregressed breeding values as pseudo-phenotypes, identified a limited number of peaks for all 30 dairy traits analyzed (Wang *et al*. 2025). Single-step GWAS of hundreds of thousands of genotyped pigs from purebred maternal lines, which is analogous to full-GRM mixed-model association despite its APY approximation (Leite *et al*. 2024), showed a similar pattern: only a few significant association clusters for backfat thickness (Kayondo *et al*. 2026). A full-GRM mixed-model GWAS of ~27,000 Duroc pigs with 11.7 million quality-controlled sequence variants reported just 10 association peaks (Wang *et al*. 2025b). By contrast, fastGWA analysis of pig samples of similar or smaller size reported substantially more associations (Zeng *et al*. 2024). The choice of method appears to be the primary driver of the discrepancy. Such large differences between full-GRM, LOCO, and sparse-GRM methods have not, to our knowledge, been reported in human GWAS. This raises several questions: what explains the large discrepancy between methods in livestock? What are the inherent limitations of each approach? Is there a ceiling on GWAS power under the full-GRM mixed model, and if so, what determines it? And which method is most appropriate for livestock GWAS?

Second, **genetic fine-mapping is difficult in livestock**. The livestock genome is characterized by long-range, strong LD, a direct consequence of small *N*_*e*_, and this makes it challenging to resolve significant associations to individual causal variants (Wang *et al*. 2025). Candidate regions often span hundreds of kbp or multiple Mbp. How difficult is fine-mapping in livestock quantitatively, and how large must a causal effect be to be resolvable given the genotypes and phenotypes of a reasonably large sample?

Answers to these questions have direct practical implications. GWAS is a critical tool for identifying genes and variants underlying economically important traits in breeding programs and for downstream integration analyses, including colocalization and Mendelian randomization, that probe biological mechanisms. Without understanding the theoretical limits of GWAS methods in livestock, we risk misusing statistical approaches and reporting misleading findings. This concern is particularly acute for methods originally developed and validated in human genetics, where *N*_*e*_ is orders of magnitude larger: a method that performs well in humans may behave very differently in livestock, and these differences need to be understood from first principles rather than assumed away.

In this study, we develop a unified theoretical framework with explicit closed-form formulas that quantify and predict how *N*_*e*_ (through *M*_*e*_) governs the mixed-model association power ceiling, the difficulty of fine-mapping, and the ease of genomic prediction. Starting from the score test under the mixed model, we derive the non-centrality parameter (NCP) for a focal SNP and show that full-GRM mixed-model association power is bounded by a practical ceiling at *M*_*e*_ and that fine-mapping resolution is governed by 4*N*_*e*_-scaled LD decay. Variants whose effects are too small to be detected at this full-GRM power ceiling are generally not fine-mappable to a reasonably high resolution either (e.g., tens of kbp), establishing a unified effect-size floor for genotype-phenotype associations. Conversely, the same low effective dimensionality makes genomic prediction reliable, explaining the chip-sufficiency for prediction noted above and revealing a trade-off in which prediction is easiest where mapping is hardest. We further show that LOCO power grows without this ceiling, but that LOCO detects **block-level** signals, aggregate LD-weighted effects that include the polygenic background, rather than the **SNP-level** excess effects identified by the full-GRM approach. This distinction compromises the utility of LOCO for any analysis where SNP-level resolution is essential, such as declaring significantly associated genes or colocalization with other omics features. Because *N*_*e*_ and *M*_*e*_ differ by orders of magnitude between livestock and humans (*M*_*e*_ ~ 10^3^–10^4^ vs ~10^6^), the same *M*_*e*_-sensitive method can exhibit fundamentally different behavior across species. Since SNP-level resolution is required for the downstream analyses that motivate most livestock GWAS (e.g., candidate-gene identification, fine-mapping, and colocalization with molecular QTL), we recommend full-GRM approaches for livestock GWAS. We validate the theory with coalescent simulations and real livestock genotype data, and translate it into practical detection and fine-mapping thresholds for cattle, pig, and chicken GWAS.

Although motivated by livestock GWAS, the framework developed here is not livestock-specific: it describes how effective genomic dimensionality determines the relative difficulty of prediction and mapping across populations. Small-*N*_*e*_ populations such as livestock provide the clearest low-*M*_*e*_ regime, in which the sample size *N* can exceed the effective genomic dimensionality *M*_*e*_, making genomic prediction comparatively easy, while long-range LD and the full-GRM power ceiling constrain SNP-level association and fine-mapping. Humans are the contrasting high-*M*_*e*_ regime, in which association and fine-mapping continue to benefit from growing sample sizes and shorter LD, whereas polygenic score prediction remains comparatively data-hungry because far more effective genomic dimensions must be learned.

## Theory

This section presents the central theoretical results. Full derivations and technical assumptions are given in the Supplementary Methods. Equation numbering is shared between the main text and the Supplementary Methods. Sections with an “S” prefix (e.g., Section S5.2) reside in the Supplementary Methods.

### Notation and Setup

#### Generative model

Let **y** be the *N* × 1 vector of standardized phenotypes and **Z** = [**z**_1_, …, **z**_*M*_] the *N* × *M* matrix of standardized genotypes (with zero mean and unit variance) for the *M* model SNPs (these need not be true causal variants; their effects are reflected through LD with the causal variants they tag). The phenotype is generated by

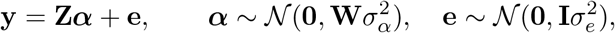

where the diagonal matrix **W** = diag(*W*_11_, …, *W*_*MM*_) encodes the effect-variance architecture: most entries are close to 1, while a few corresponding to major-effect loci are substantially larger. The true **W** is unknown. We define 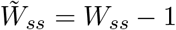 as the excess effect of SNP *s* above the polygenic expectation; under a purely polygenic architecture 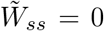, while a major QTL has 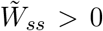. Total phenotypic variance is normalized to one 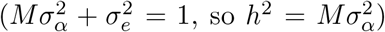. In practice, a moderate-density chip panel (*M* ~ 50,000) adequately captures the genomic signal when *M* ≫ *M*_*e*_, the effective number of independent genomic segments. Further details on this generative model are given in Section S1.1.

Intuitively, **W** is the shape of the trait’s genetic architecture: most SNPs contribute the background per-SNP effect variance (*W*_*ss*_ ≈ 1), while a small number of QTL contribute disproportionately (*W*_*ss*_ ≫ 1).

#### Working model

Association testing assumes equal effect variances (**W** = **I**) and tests a focal SNP **x** as a fixed effect:

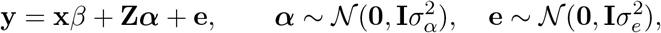

with phenotypic covariance **V** = **G***h*^2^ + **I**(1 − *h*^2^) where **G** = **ZZ***T /M* is the GRM. Under **full-GRM** analysis, **Z** includes all *M* model SNPs. Throughout, “full-GRM” refers to a GRM that adequately tags the genomic segment containing the tested variant, not necessarily a GRM that literally contains the tested marker itself. Including the tested SNP in the GRM, excluding only the tested marker (leave-one-marker-out), or testing imputed or sequence variants against a chip-based GRM all fall under the same full-GRM framework when nearby markers in LD with the tested variant remain in the GRM. Under **leave-one-chromosome-out (LOCO)** analysis, **Z** excludes SNPs on the focal chromosome, so the focal chromosome’s genetic variance is absorbed into the residual. REML estimation of variance components is robust to moderate deviations from the equal-variance assumption.

Intuitively, the test detects QTL by asking whether the focal SNP carries trait variance that the random effect has not already explained. Full-GRM and LOCO differ in what the random effect absorbs: under full-GRM, **Z** includes the focal-region SNPs, so their polygenic baseline is absorbed and the test sees only the focal excess; under LOCO, **Z** excludes them, so the focal-region variance remains in the residual and the test sees the full effect.

#### Spectral quantities

The GRM has eigenvalues *d*_1_ ≥ · · · ≥ *d*_*r*_ ≥ 0 with *r* = rank(**Z**), and the corresponding LD matrix **R** = **Z**^*T*^**Z***/N* has eigenvalues *ℓ*_*k*_ = *Md*_*k*_*/N* satisfying _*k*_ *ℓ*_*k*_ = *M*. We write *λ* = (1 − *h*^2^)*/h*^2^ for the noise-to-signal ratio. Full notation is in Section S1.6.

Geometrically, each eigenvalue *d*_*k*_ indexes one independent direction of variation in the chip-genotype data. Large *d*_*k*_ corresponds to a major axis along which many SNPs co-vary because of common ancestry, pedigree relatedness, or large haplotype blocks; small *d*_*k*_ corresponds to fine-grained, near-independent variation. The dominant eigenvalues therefore capture most of the patterns of genetic relatedness that mixed-model methods depend on.

### 2.1 The per-SNP non-centrality parameter

Under the null hypothesis, single-SNP association test statistics follow a central chi-squared distribution with one degree of freedom. Under the alternative hypothesis (with non-zero effect), the distribution shifts right to a non-central chi-squared; the non-centrality parameter (NCP) is the value of this shift, and a larger NCP corresponds to higher statistical power.

Under the full-GRM analysis with REML-estimated variance components, the score chi-squared statistic for focal SNP *l* has the non-centrality parameter (Sections S2–S4)

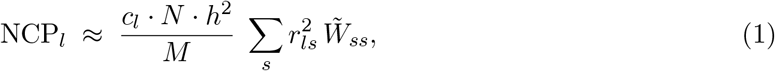

where *r*_*ls*_ is the sample LD correlation between the focal SNP and GRM SNP *s*, and *c*_*l*_ is the **per-SNP GRAMMAR-Gamma coefficient**

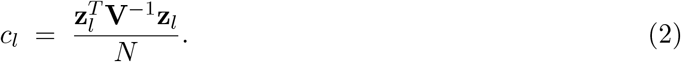

A focal SNP under full-GRM analysis effectively lies in col(**Z**) regardless of whether it is literally a GRM SNP (Section S2.4). We therefore use **z**_*l*_ to represent it in Equation 2.

The original GRAMMAR-Gamma method (Svishcheva *et al*. 2012) applies a single genome-wide scalar (the “gamma” factor) to correct score statistics computed on mixed-model residuals. The coefficient *c*_*l*_ above is the **per-SNP generalization** of that scalar; it varies across SNPs, with the mechanism made explicit by the eigenvalue expansion below.

The eigenvalue expansion of *c*_*l*_ reveals the mechanism of proximal contamination:

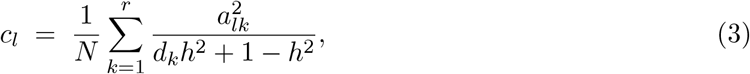

where 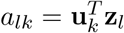 is the projection of the focal SNP onto the *k*-th GRM eigenvector (**u**_*k*_). Because the denominator grows with *d*_*k*_, projections onto large-eigenvalue modes are heavily shrunk. For instance, consider a SNP that aligns with a dominant axis of GRM variation (e.g., an ancestry-informative SNP, or one tagging a large LD block). Mathematically, the SNP’s projection concentrates on large-eigenvalue modes of the GRM, exactly those most heavily shrunk in Equation 3, so *c*_*l*_ is correspondingly small (Section S4.2). Intuitively, the polygenic random effect strongly competes for the focal-region variance and absorbs much of it as background, leaving less for the fixed-effect test to detect.

The architectural factor 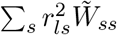 aggregates the *r*^2^-weighted excess effect-variance contributions of all GRM SNPs in LD with the focal SNP. For a single focal signal whose effect is fully captured by the generative model and local LD tagging (the focal SNP need not be the literal causal variant; the signal’s effect can be carried by the focal SNP directly or distributed across nearby GRM SNPs in tight LD), this sum approximately reduces to *q/*(*h*^2^*/M*) − 1 in the ideal tagging case, where *q* is the focal SNP’s proportion of phenotypic variance explained (PVE). Equivalently, *q/*(*h*^2^*/M*) is a **fold-enrichment factor** (the focal SNP’s PVE relative to the per-SNP average), and subtracting 1 strips off the polygenic background. The architectural factor therefore quantifies how much trait variance is concentrated at the focal locus in excess of the polygenic background; this is what the test detects under the full-GRM GWAS.

### 2.2 The 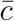 approximation and the sigmoid sum

Averaging *c*_*l*_ over the *M* GRM SNPs and using 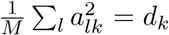 (Section S4.3) gives the **average GRAMMAR-Gamma coefficient**

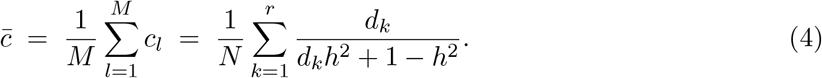

Although 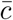 is an average over the *M* GRM SNPs (Equation 4), it can approximate the per-SNP GRAMMAR-Gamma coefficient for any tested variant (e.g., a sequence variant outside the GRM). When the full-GRM SNPs adequately tag the genomic segments containing the test variants, those variants share similar projection patterns onto the GRM eigenspace as the GRM SNPs themselves. We verify the modest variation of *c*_*l*_ around 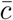 in real livestock chip data in Section 4.1. Replacing the per-SNP *c*_*l*_ with the genome-wide average 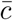 is the GRAMMAR-Gamma approximation; substituting it into (1) and rewriting in terms of LD-matrix eigenvalues gives

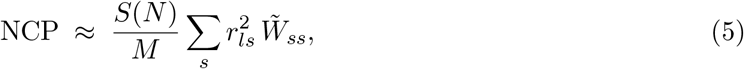

where the **sigmoid sum**

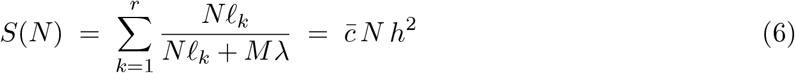

is the sum of *r* sigmoid functions of *N* (strictly speaking, each sigmoid in log *N*), one per LD eigenmode. Each mode contributes a sigmoid that rises from 0 to 1 with *N*, with half-saturation at the mode-specific scale 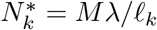. The sample-size dependence of full-GRM GWAS power is therefore entirely encoded in *S*(*N*): intuitively, *S*(*N*) is the effective number of LD eigenmodes the data have captured at sample size *N*. The NCP is exactly invariant to SNP duplication in the GRM (Section S4.5), confirming that power is governed by the spectral structure of the GRM, not the raw marker count.

#### NCP in the ideal tagging case

When a focal signal is fully captured by the generative model and local LD tagging, 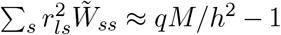 (Section 2.1). Substituting into Equation 5 gives the compact form

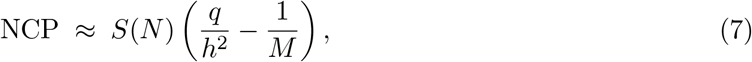

where *q* is the focal SNP’s PVE. At chip scale (*M* ~ 50,000) the 1*/M* term is negligible compared to *q/h*^2^ in practice, so NCP ≈ *S*(*N*) *q/h*^2^.

### 2.3 Two ceilings on *S*(*N*) and the practical bound *M*_*e*_ = 4*N*_*e*_*L*

*S*(*N*) is strictly increasing and strictly concave in *N* (Section S6) and admits two distinct ceilings. The **hard ceiling** is *S*(*N*) → *M* as *N* → ∞ (every eigenmode eventually saturates). The **practical ceiling** is *M*_*e*_ = 4*N*_*e*_*L*, the number of independent genomic segments predicted by drift–recombination theory (Sved 1971; Stam 1980) and observed empirically as the number of GRM eigenvalues carrying ~98% of the total spectral mass (Pocrnic *et al*. 2016). The “practical” qualifier reflects that currently achievable livestock sample sizes (e.g., ~10^6^) can bring *S*(*N*) near *M*_*e*_, while remaining far below the hard ceiling *M*.

To make this precise, partition the *M* LD eigenvalues into Group A (the top *M*_*e*_, carrying mass fraction *ρ* ≈ 0.98–0.99) and Group B (the remaining *M*_*B*_ = *M* − *M*_*e*_ near-zero eigenvalues). Jensen’s inequality applied to each group (Section S7.3) yields two transition scales (sample-size checkpoints at which *S*(*N*)’s growth regime changes):

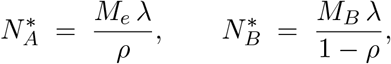

with 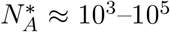 and 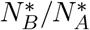 on the order of 10^2^–10^3^ for livestock chip panels (Section S7.4). In the **practical regime** 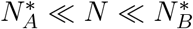, Group A is nearly saturated while Group B remains negligible, giving

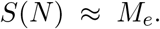

The NCP under the 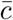 approximation therefore takes two limiting forms (Sections S7.5 and S8):

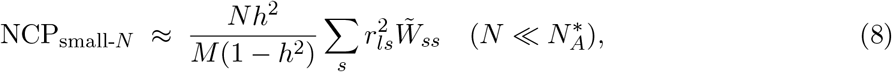

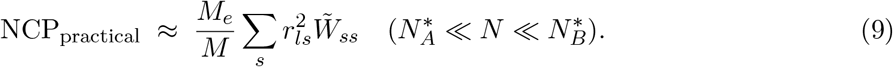

The practical-ceiling NCP (9) is approximately independent of *N* : once sample size is much larger than 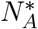, additional samples produce negligible further increase in full-GRM association power. When *N* is well below 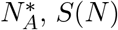 largely grows linearly with *N* (Equation 8) and additional samples translate directly into added detection power; above 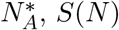 approaches *M*_*e*_ concavely (Figure 3), with diminishing returns. This is the theoretical explanation for the first empirical pattern noted in the Introduction. In modern livestock GWAS, sample sizes can be well above 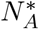, so the practical ceiling can be reached at achievable sample sizes. In human GWAS, *M*_*e*_ ≈ 1.4 × 10^6^ and 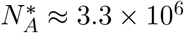 (for *h*^2^ = 0.3), so most studies operate in the small-*N* regime (8) where all three approaches (linear regression, LOCO, and full-GRM GWAS) scale similarly with *N* (Section S8).

Substituting the practical-regime saturation *S*(*N*) ≈ *M*_*e*_ into Equation 7 and requiring NCP ≈ 30 at genome-wide significance (*p* < 5 × 10^−8^, equivalently *χ*^2^ > 29.7; Section S8 clarifies why *M*_*e*_ is not the multiple-testing divisor) for 50% power yields the minimum detectable PVE (Section S8):

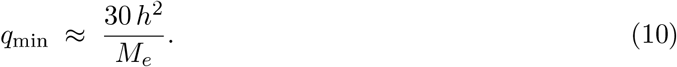

**Table 1.**
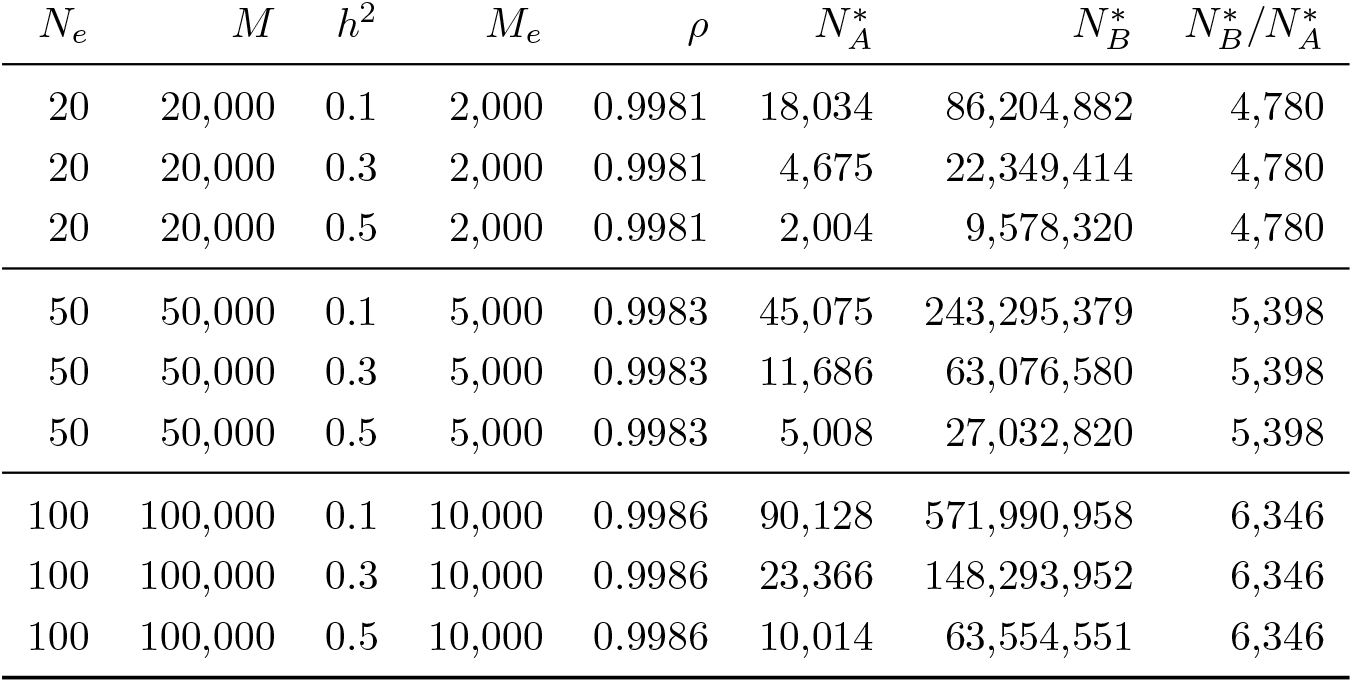
Eigenspectrum parameters at the theoretical *M*_*e*_ = 4*N*_*e*_*L*. For each combination of effective population size and heritability, the table reports the total marker count *M*, the theoretical *M*_*e*_, the proximal-mass fraction *ρ* (sum of eigenvalues held by the top *M*_*e*_ modes, normalized by the total spectral mass), and the two transition scales 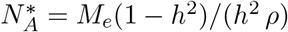 and 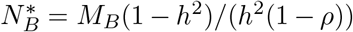 that delimit the practical regime 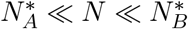. The ratio 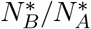 in the final column quantifies the width of the practical regime. Eigenvalues are computed at the largest available sample size for each *N*_*e*_ (*N* = 100,000 for *N*_*e*_ = 20 and 50; *N* = 200,000 for *N*_*e*_ = 100). The two alternative *M*_*e*_ definitions, based on capturing 98% and 99% of total eigenvalue mass, are reported in Supplementary Tables 6 and 7, respectively.

| $N_e$ | $M$ | $h^2$ | $M_e$ | $\rho$ | $N_A^*$ | $N_B^*$ | $N_B^*/N_A^*$ |
| --- | --- | --- | --- | --- | --- | --- | --- |
| 20 | 20,000 | 0.1 | 2,000 | 0.9981 | 18,034 | 86,204,882 | 4,780 |
| 20 | 20,000 | 0.3 | 2,000 | 0.9981 | 4,675 | 22,349,414 | 4,780 |
| 20 | 20,000 | 0.5 | 2,000 | 0.9981 | 2,004 | 9,578,320 | 4,780 |
| 50 | 50,000 | 0.1 | 5,000 | 0.9983 | 45,075 | 243,295,379 | 5,398 |
| 50 | 50,000 | 0.3 | 5,000 | 0.9983 | 11,686 | 63,076,580 | 5,398 |
| 50 | 50,000 | 0.5 | 5,000 | 0.9983 | 5,008 | 27,032,820 | 5,398 |
| 100 | 100,000 | 0.1 | 10,000 | 0.9986 | 90,128 | 571,990,958 | 6,346 |
| 100 | 100,000 | 0.3 | 10,000 | 0.9986 | 23,366 | 148,293,952 | 6,346 |
| 100 | 100,000 | 0.5 | 10,000 | 0.9986 | 10,014 | 63,554,551 | 6,346 |

For cattle (*N*_*e*_ ≈ 100, *L* ≈ 25 Morgans, and *h*^2^ = 0.3), *M*_*e*_ ≈ 10,000 and *q*_min_ ≈ 0.09%. This is the practical detection floor for full-GRM GWAS for any practically achievable *N*. The detection floor *q*_min_ is robust to reasonable choices of GWAS power and significance threshold, remaining on the order of tens of *h*^2^*/M*_*e*_ (e.g., *q*_min_ ≈ 10*h*^2^*/M*_*e*_ for 1% power at *p <* 5 × 10^−8^, or *q*_min_ ≈ 24*h*^2^*/M*_*e*_ for 50% power at *p <* 10^−6^).

### 2.4 *N* –*h*^2^ exchange under fixed genetic architecture

The sigmoid sum *S*(*N*) depends on sample size *N* and heritability *h*^2^ only through one combination. Each term of *S*(*N*) is *Nℓ*_*k*_*/*(*Nℓ*_*k*_ + *Mλ*), so two scenarios 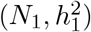 and 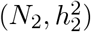 yield identical *S*(*N*) term-by-term (and hence identical sums) whenever the ratio *Nℓ*_*k*_*/Mλ* is preserved. Treating the *ℓ*_*k*_ as population-level eigenvalues (sample LD matrices converge to the population spectrum), each *ℓ*_*k*_ drops out, and the conserved quantity is

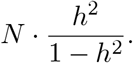

Solving for *N*_1_ in terms of *N*_2_ gives the *N* **-equivalence formula** (Section S9.1)

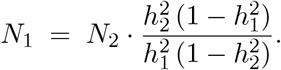

#### Pseudo-phenotype leverage

This exchange has direct practical relevance for livestock GWAS that use de-regressed breeding values (or de-regressed proofs; DRPs) as pseudo-phenotypes (e.g., dairy bull studies with highly reliable proofs). DRP rescales the phenotype so that its effective heritability approximately equals the reliability, 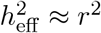. Crucially, DRP transformation preserves the *genetic architecture*: each causal variant’s relative contribution to total genetic variance is unchanged (Section S9.2). The aggregate weight 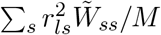 in the *S*(*N*) form of the NCP (Equation 5) is therefore invariant under DRP, and the NCP equivalence between two scenarios reduces to the *S*(*N*) equivalence above. Setting scenario 1 to “original phenotype with single-record heritability 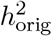” and scenario 2 to “DRP-based analysis with cohort-average reliability *r*^2^”,

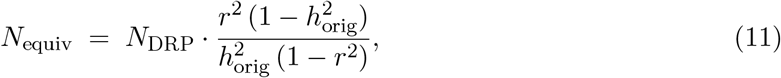

where *N*_equiv_ is the sample size of original-phenotype records that would yield the same per-SNP detection power as *N*_DRP_ records of DRPs. The cohort-average *r*^2^ substitution is a simplification of the actual per-individual reliability heterogeneity in DRP cohorts.

Intuitively, a moderate-size sample of individuals with reliable DRPs can match the GWAS power of a much larger sample with single-record phenotypes; in dairy bull GWAS the equivalence multiplier *N*_equiv_*/N*_DRP_ can reach about 16 (Section S9.2). This multiplier underlies the practical leverage of breeding-value GWAS in livestock. Because the leverage acts through *S*(*N*), it raises genomic prediction reliability as well as detection power (Section 2.7).

### 2.5 LOCO: block-level signals and the LOCO coefficient

Under leave-one-chromosome-out (LOCO), the GRM is constructed from SNPs *excluding* the focal chromosome. Because the focal SNP **x** has no physical linkage with any GRM SNP, focal-region genetic variance is no longer absorbed by the random effect, and the association statistic becomes a regional, LD-aggregated signal. The LOCO coefficient (Section S5.2)

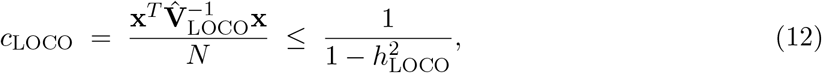

where 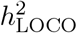 is the PVE by SNPs outside the focal chromosome. The upper bound is approached *N* ≫ *M*_LOCO_ (the number of LOCO SNPs) and pedigree structure is weak. In livestock, pedigree relationships create correlations between the focal SNP and the LOCO-GRM SNPs, which reduces *c*_LOCO_ below the ideal bound (Section S5.2). The LOCO NCP is

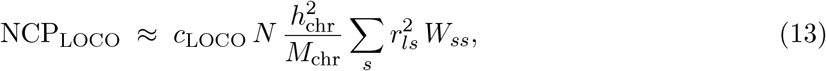

where 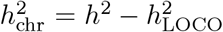 and *M*_chr_ are the PVE and number of SNPs on the focal chromosome, respectively.

Two structural features distinguish (13) from the full-GRM NCP (1):

1. **Linear, unbounded growth in** *N*. Because *c*_LOCO_ is not limited by *N* (in the practical regime), the LOCO NCP grows linearly with sample size and has no analogue of the practical ceiling (9). LOCO power keeps increasing as *N* grows.
2. *W*_*ss*_ **rather than** 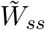. The focal chromosome’s genetic variance 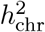 is entirely unmodeled under LOCO, so the test statistic captures the **full** effect of each focal-region SNP, not just its excess above the polygenic expectation. Under a weak independence assumption (that 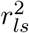 and *W*_*ss*_ are approximately uncorrelated), the LOCO NCP reduces to 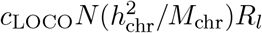, where 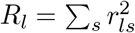 is the focal SNP’s LD score (Section S14.1). In a region of strong long-range LD (large *R*_*l*_), the LOCO NCP can be substantial *without any large- or moderate-effect QTL being present*, because *R*_*l*_ alone is enough to accumulate significance once *N* is large.

In contrast, the full-GRM NCP at the practical ceiling (9) is negligible for diffuse polygenic effects, because 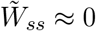. **The full-GRM mixed model filters block-level polygenic signals; LOCO does not** (Section S14). This is the theoretical basis for the first empirical pattern noted in the Introduction: at large *N* in livestock populations with long-range LD, LOCO produces inflated test statistics across broad genomic regions while the full-GRM approach identifies a small number of association peaks.

### 2.6 Fine-mapping resolution

Fine-mapping asks which SNP among a set of LD-linked candidates carries the causal effect. Under simple regression (equivalently, the idealized case in which polygenic confounding is set aside) with a true causal SNP *i* of effect *β*_*i*_ (for standardized phenotypes), the expected log-likelihood difference between the true model (for the causal SNP *i*) and an alternative model (for SNP *j*) is (Section S11.2)

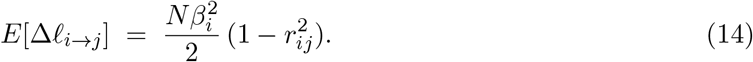

Intuitively, Δ*ℓ* measures the data’s confidence in distinguishing the true causal SNP from an LD-correlated alternative: it scales with both the effect size 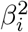 (per-SNP PVE) and the LD imperfection 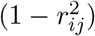 between the two SNPs.

The same form holds for the LOCO and full-GRM mixed models, with an effective coefficient multiplying the simple-regression expression (Sections S12.2 and S12.3):

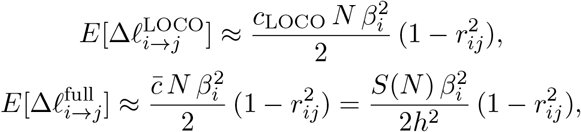

where 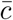 is the genome-wide average GRAMMAR-Gamma coefficient (Equation 4), *c*_LOCO_ its LOCO counterpart (Section 2.5), and 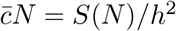 (Equation 6). All three frameworks (simple regression, LOCO, and full-GRM) share the same 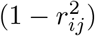 factor governing within-block resolution and differ only in the effective coefficient. Because *S*(*N*) saturates at *M*_*e*_ in the practical regime (Sections 2.3 and S12.3), the full-GRM fine-mapping evidence is practically bounded, paralleling the detection NCP ceiling (9). In contrast, the LOCO and simple-regression evidence grow without bound in *N*.

The LD decay under drift–recombination equilibrium (Sved 1971) gives

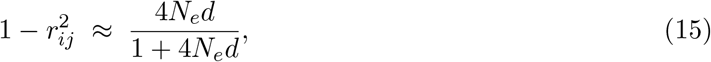

where *d* is the genetic distance between SNPs *i* and *j* in Morgans. Fine-mapping resolution is therefore governed by the same 4*N*_*e*_-scaling that governs the practical full-GRM NCP ceiling (Equation 9): as *M*_*e*_ = 4*N*_*e*_*L* for effective genomic dimensionality and as 4*N*_*e*_*d* for LD decay (Equation 15). Intuitively, smaller *N*_*e*_ allows LD to persist over much larger distances: in livestock (*N*_*e*_ ≈ 50–200), substantial LD extends across multi-Mbp regions; in humans (*N*_*e*_ ≈ 10,000), LD decays much more rapidly with distance. The same small *N*_*e*_ that limits full-GRM detection power therefore also enlarges the fine-mapping uncertainty in livestock.

Setting *E*[Δ*ℓ*_*i*→*j*_] > *θ* for a decision threshold *θ* (e.g., *θ* = 3 for a Bayes factor ≈ 20) and substituting Equation (15) in Equation (14) gives the minimum effect size resolvable at distance *d* in a sample of size *N* (Section S13.3):

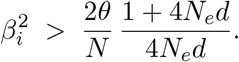

Under the full-GRM mixed model, where *S*(*N*) → *M*_*e*_ (its practical ceiling; Section 2.3), the analogous threshold is

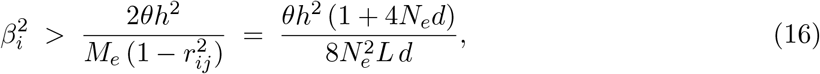

which depends only on *N*_*e*_, *L, h*^2^, and the inter-marker distance *d*; no further increase in *N* can practically lower this threshold (Section S13.4). The floor further implies that polygenic effects within an LD block cannot be successfully fine-mapped with genotype-phenotype associations alone: when each causal variant explains only ~*h*^2^*/M* of phenotypic variance, the per-SNP PVE 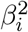 falls below both fine-mapping thresholds at a 10 kb resolution for any practical *N* (Section S13.2).

For cattle (*N*_*e*_ ≈ 100 and *d* = 10 kb ≈ 10^−4^ Morgans), 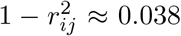; at *θ* = 3, the minimum resolvable PVE is 0.16% under simple regression at *N* = 100,000 and 0.47% under the full-GRM method at *h*^2^ = 0.3 and any practically achievable *N*. Comparing with the full-GRM GWAS detection floor (10) at the same *h*^2^ (*q*_min_ ≈ 0.09%): both fine-mapping thresholds exceed the detection threshold. Variants whose effects are too small to be detected at the practical full-GRM NCP ceiling are generally not fine-mappable either to a reasonably high resolution (e.g., tens of kbp). This unified effect-size floor, established quantitatively across species in Section S15, is the theoretical basis for the second empirical pattern noted in the Introduction.

### 2.7 Genomic prediction reliability

Genomic prediction reliability is governed by the same sigmoid sum *S*(*N*) as mixed-model association power. The per-SNP association NCP scales with *S*(*N*) (Equations 5 and 7), and the in-sample reliability of SNP-BLUP (equivalently, GBLUP), namely the fraction of genetic variance the model recovers within the training set, is

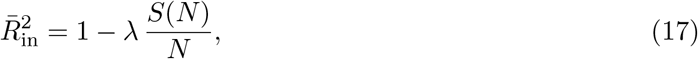

a quantity in (0, 1) that rises toward 1 as *S*(*N*)*/N* falls (Section S10). In the regime 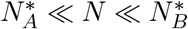 (practical for livestock) it is approximated by

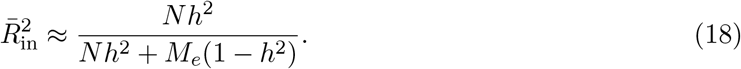

Reliability for individuals outside the training set is lower. The leave-one-out reliability, a proxy for out-of-sample prediction, satisfies

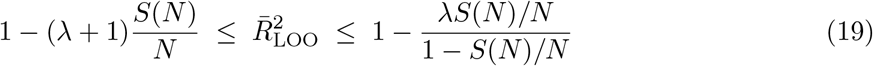

and lies below 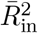 (Section S10.6). Both in-sample and leave-one-out reliabilities are thus controlled by the single ratio *S*(*N*)*/N*. We have quantified *S*(*N*)’s growth patterns and practical ceiling *M*_*e*_ in Section 2.3.

Equations 17 and 19, coupled with the characteristics of *S*(*N*), show contrasting outcomes for GBLUP-like methods in livestock and humans. Because *M*_*e*_ is small in livestock (~10^3^–10^4^), genomic evaluation programs can have an *N* (i.e., reference population size) large enough to drive *S*(*N*)*/N* ≈ *M*_*e*_*/N* ≪ *h*^2^, so 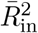 and 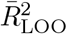 are both close to 1: prediction is reliable and transfers out of sample. Because *M*_*e*_ is large in humans (~10^6^), even biobank-scale *N* (e.g., 5 × 10^5^) is lower than *M*_*e*_ and leaves *S*(*N*)*/N* of order *h*^2^, so in-sample reliability is moderate and out-of-sample reliability small. In the *N* ≪ *M*_*e*_ regime, 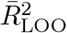 and the classical Daetwyler reliability 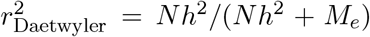 (Daetwyler *et al*. 2008, 2010) both reduce to *Nh*^2^*/M*_*e*_, so our framework recovers the classical Daetwyler result as a special case (Section S10.7).

The sigmoid sum *S*(*N*), with practical ceiling *M*_*e*_, is common to the full-GRM NCP and the reliability formulas above, so the single quantity *M*_*e*_ bounds both association power and prediction reliability. Its two effects run in opposite directions: a small *M*_*e*_ caps the number of distinguishable above-background association signals yet drives the reliabilities toward 1 with practically achievable *N*. Genomic prediction is therefore comparatively easy precisely where SNP-level mapping is comparatively hard.

#### Summary

Equations 1–19 collectively show that a single population parameter (*N*_*e*_) governs, through two related quantities (*M*_*e*_ = 4*N*_*e*_*L* and 4*N*_*e*_*d*-scaled LD decay), the limiting behaviors of three otherwise-disparate types of mixed-model genomic analysis: GWAS, fine-mapping, and GBLUP. The per-SNP association NCP is captured by a sigmoid sum *S*(*N*) over LD eigenmodes (Equations 1–7) that grows toward a practical ceiling *M*_*e*_ (Equations 8 and 9), imposing a detection floor *q*_min_ ≈ 30*h*^2^*/M*_*e*_ (Equation 10). The same *S*(*N*) sets GBLUP reliability, both in-sample (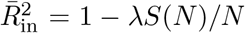; Equations 17 and 18) and out-of-sample (Equation 19), so the ceiling *M*_*e*_ bounds detection power and prediction reliability. Because *S*(*N*) depends on *N* and *h*^2^ only through *Nh*^2^*/*(1 − *h*^2^), deregressed breeding values as pseudo-phenotypes amplify the effective sample size in both association and prediction (Equation 11). LOCO grows without this ceiling but detects block-level rather than SNP-level signals (Equations 12 and 13), while fine-mapping resolution is governed by 4*N*_*e*_*d*-scaled LD decay (Equations 14–16). In livestock, the same small *M*_*e*_ that imposes a unified effect-size floor on GWAS detection and fine-mapping resolution makes GBLUP reliable. Genomic prediction is comparatively easy precisely where SNP-level mapping is hard. The remainder of the paper validates the per-SNP NCP formula with phenotype simulations, verifies the predicted concave approach of *S*(*N*) toward the *M*_*e*_ ceiling with msprime eigenvalue spectra, empirically checks the 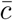 approximation and the LOCO bound in real livestock chip data, and translates the theory into practical thresholds for cattle, pig, and chicken GWAS.

## Materials and Methods

### 3.1 Real livestock chip-panel datasets and GRAMMAR-Gamma coefficients

We computed the per-SNP GRAMMAR-Gamma coefficient *c*_*l*_ (Equation 2) in three publicly available livestock single-nucleotide-polymorphism (SNP) chip datasets. The Chinese Holstein cattle dataset (Huang *et al*. 2019) comprises *N* = 2,510 cows genotyped on the Illumina BovineSNP50 BeadChip; after standard quality control (QC; call rate ≥ 0.95, minor allele frequency (MAF) ≥ 0.01, and Hardy–Weinberg equilibrium *p >* 10^−6^), *M* = 42,775 autosomal SNPs were retained. The Karacabey Merino sheep dataset (Yaman *et al*. 2025) comprises *N* = 734 animals genotyped on the Illumina OvineSNP50 BeadChip, with *M* = 35,774 post-QC SNPs. The German Holstein cattle dataset (Zhang *et al*. 2015) comprises *N* = 5,024 bulls genotyped on the Illumina BovineSNP50 BeadChip, with *M* = 42,216 post-QC SNPs.

For each dataset, the genomic relationship matrix (GRM) **G** = **ZZ**^⊤^*/M* was constructed from the column-standardized genotype matrix **Z** (each SNP centered to zero mean and scaled to unit variance within the dataset; VanRaden 2008). The working-model variance matrix **V** = *h*^2^**G** + (1 − *h*^2^)**I** was formed at three working-model heritabilities, *h*^2^ ∈ {0.1, 0.3, 0.5}. The per-SNP coefficient 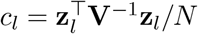 was then computed for every autosomal SNP in each dataset and at each *h*^2^.

For the two datasets with chromosome maps (Chinese Holstein and Karacabey Merino), the leave-one-chromosome-out (LOCO) coefficient *c*_LOCO_ was computed under the working-model variance 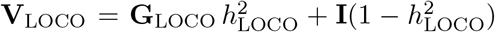, where **G**_LOCO_ is the GRM constructed from all autosomal SNPs except those on the chromosome harboring the focal SNP *l*. We specified 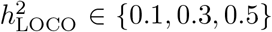, the same three values as the full-GRM working-model heritability. The German Holstein dataset has no chromosome map and was used for full-GRM analysis only.

### 3.2 Coalescent simulations and phenotype simulations

Our coalescent simulations are intended to test the spectral mechanism implied by the theory, not to reproduce the complete demographic history of any livestock species. Real breeding populations experience bottlenecks, selection, admixture, family structure, and recent genomic selection, all of which may alter the detailed shape of the leading eigenvalue spectrum. The key assumption required by our theoretical framework is weaker: in small-*N*_*e*_ livestock populations, a relatively small number of leading eigenvalues carries most of the LD-matrix spectral mass, with effective dimension on the order of *M*_*e*_ = 4*N*_*e*_*L*. Prior effective-dimensionality studies (Pocrnic *et al*. 2016) and analyses based on the Algorithm for Proven and Young (APY; Misztal *et al*. 2014; Pocrnic *et al*. 2016b) support this order of magnitude by showing that the number of leading eigenvalues explaining approximately 98% of total spectral mass, or the APY core size required for genomic prediction stability, is close to 4*N*_*e*_*L*.

Coalescent genotype matrices were generated with msprime v1.3.4 (Baumdicker *et al*. 2022) under the Hudson coalescent. Each population was simulated under a two-epoch demographic model so that variants would be old enough to be embedded in chip-scale LD: a recent epoch with effective size *N*_*e*_ ∈ {20, 50, 100} for 10 *N*_*e*_ generations, followed by an ancestral epoch with 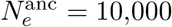. Genome architecture matched the cattle reference (ARS-UCD1.2): 29 autosomes ranging from 158.5 Mb (BTA1) to 42.9 Mb (BTA25), totalling ~2.5 Gb. Recombination rate was uniform at 10^−8^ per base pair per generation, giving total genetic length *L* ≈ 25 Morgans. Mutations were generated under the binary-mutation model at rate 5 × 10^−8^ per base pair per generation. As a stylized approximation to chip ascertainment, which in practice involves more elaborate selection criteria (e.g., common-variant prioritization, genotype-calling robustness, and even spacing across the genome) than we model here, we retained only biallelic segregating SNPs with MAF ≥ 0.05, then randomly thinned the retained SNPs on each chromosome to a target count proportional to chromosome length, yielding exact totals *M* = 20,000 for *N*_*e*_ = 20, *M* = 50,000 for *N*_*e*_ = 50, and *M* = 100,000 for *N*_*e*_ = 100. Diploid sample sizes were *N*_full_ = 100,000 for *N*_*e*_ = 20 and *N*_*e*_ = 50, and *N*_full_ = 200,000 for *N*_*e*_ = 100 to permit eigenvalue analysis at the largest sample sizes for the largest effective size.

To validate the per-SNP non-centrality parameter (NCP) formula (Equation 1), we simulated phenotypes under the linear mixed model. From each population, five focal SNPs were preselected once and reused across all subsequent scenarios. One SNP was selected per chromosome 1–5, with chromosome-specific target MAFs of 0.075, 0.175, 0.275, 0.375, and 0.475, respectively; the SNP whose MAF (computed from all *N*_full_ individuals) was closest to the chromosome’s target was selected. This design covers the allele-frequency spectrum and ensures that the five SNPs sample independent linkage-disequilibrium (LD) environments.

Phenotypes were simulated on a 3 × 3 × 2 × 4 grid: three effective sizes (*N*_*e*_ = 20, 50, 100), three trait heritabilities (*h*^2^ = 0.1, 0.3, 0.5), two focal-SNP variance contributions (*β*^2^ = 0.001 and 0.01, where *β*^2^ is the proportion of phenotypic variance explained by each focal SNP), and four sample sizes (*N* = 5,000, 10,000, 20,000, and 50,000); the same four *N* values were used for all *N*_*e*_. For each scenario, 100 replicate phenotypes were generated under the multi-SNP effect-variance architecture of Section 2 (Notation and Setup) with 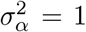. Background-SNP effects on the standardized-genotype scale were drawn as 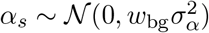 with *w*_bg_ = (*M* − 5*W*_*ss*_)*/*(*M* − 5) chosen so that the average effect-variance weight equals unity. Focal-SNP effects had fixed magnitudes 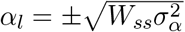 with *W*_*ss*_ = *β*^2^*M/h*^2^ on the standardized-genotype scale; the ± signs were drawn once at the start of the analysis with a fixed random seed and held constant across all 100 replicates of every scenario, so that only the background effects varied between replicates. The genomic breeding value (GEBV) was computed for each individual in each replicate, and an independent residual drawn from N (0, *M* (1 − *h*^2^)*/h*^2^) was added to give the simulated phenotype. The total phenotypic variance scales with *M/h*^2^, which is irrelevant for the score *χ*^2^ statistic but reflects the underlying linear-mixed-model variance accounting.

For each replicate, variance components were estimated by restricted maximum likelihood (REML) using SLEMM (Cheng *et al*. 2023), with the five focal SNPs included as fixed-effect covariates and the GRM constructed from all *M* SNPs of a subsample of *N* individuals. SLEMM was chosen for its fast REML implementation, which is essential for the heavy simulation workload (100 replicates across 72 scenarios). With all five focal SNPs jointly fitted as fixed effects, the chi-squared statistic for focal SNP *l* in replicate *r* was computed as 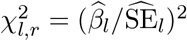, the squared ratio of SLEMM’s best linear unbiased estimate (BLUE) and standard error (SE) for that focal SNP. The empirical NCP for each focal SNP was 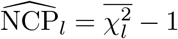, the mean across the 100 replicates.

The theoretical NCP per focal SNP was computed by way of an *N* × *N* GRM eigendecomposition. For each (*N*_*e*_, *N*) pair, we performed this eigendecomposition once on the standardized subsample, projected each focal SNP onto the resulting eigenvectors as 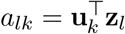, and stored the eigenvalues {*d*_*k*_} and projections {*a*_*lk*_} for reuse. The per-SNP coefficient was then 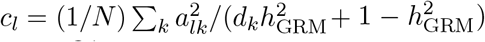 (Equation 3) and the theoretical NCP was 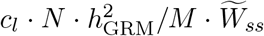 (Equation 1) with 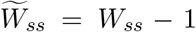. Because the five focal SNPs are fitted as fixed-effect covariates, the GRM captures only the polygenic background, so the working-model heritability of the GRM was set 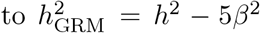 rather than the full *h*^2^. PLINK v1.9 (Chang *et al*. 2015) was used for genotype-format conversion and basic QC of the simulated genotypes.

### 3.3 LD-matrix eigenvalue spectra and average GRAMMAR-Gamma coefficients

For each population and each target sample size *N*, an LD-matrix eigenvalue spectrum was obtained as follows. A subset of *N* individuals was drawn from the simulated population without replacement; the resulting *N* × *M* genotype matrix was re-standardized within the subsample (each SNP centered to zero mean and scaled to unit variance); and the eigenvalues of the LD matrix **R** = **Z**^⊤^**Z***/N* were computed.

Two algorithms were used depending on the matrix size. For all (*N*_*e*_, *N*) scenarios with *N*_*e*_ ∈ {20, 50} at any *N*, and for *N*_*e*_ = 100 at *N* ≤ 50,000, exact symmetric eigendecomposition was applied via the dual trick: diagonalizing the smaller of the *N* × *N* GRM or the *M* × *M* LD matrix and recovering the spectrum of the full *M* × *M* LD matrix by rescaling and zero-padding (min(*N, M*) ≤ 50,000 in all of these cases). For *N*_*e*_ = 100 at *N* ∈ {100,000, 200,000} the LD matrix is *M* × *M* = 100,000 × 100,000, for which exact eigendecomposition is memory-prohibitive on our compute server (512 GB RAM). For these two scenarios we used randomized truncated singular value decomposition (SVD) as implemented in scikit-learn, retaining 20,000 singular values (with one power iteration, QR normalization, and fixed random seed). The remaining 80,000 eigenvalue slots were set to zero, and downstream summary statistics (participation ratio and the eigenvalue-fraction thresholds EIG_98%_ and EIG_99%_) were normalized by the known trace *M*_total_ = *M* rather than the truncated sum, so that truncation does not bias the reported quantities. The 20,000-eigenvalue cutoff is well above all thresholds used in the analysis (the largest is EIG_99.9%_ ≈ 10,648 for the largest case considered), so no quantity reported in Section 4.3 depends on eigenvalues beyond the truncation point. Eigenvalue spectra were computed at *N* ∈ {5,000, 10,000, 20,000, 50,000, 100,000} for *N*_*e*_ = 20 and *N*_*e*_ = 50, and at the same five values plus *N* = 200,000 for *N*_*e*_ = 100.

For each (*N*_*e*_, *N*) pair, the sigmoid sum *S*(*N*) and its Group A / Group B decomposition (Equation 6; Section S7) were computed directly from the *N*-specific eigenvalue array described above. Two definitions of the practical ceiling *M*_*e*_ were considered. The main text uses the theoretical value *M*_*e*_ = 4*N*_*e*_*L* (Sved 1971; Stam 1980), which is the central prediction of the framework. Two supplementary analyses use data-driven definitions in which *M*_*e*_ equals the smallest number of top eigenvalues whose cumulative sum captures, respectively, 98% and 99% of the total spectral mass (the “effective dimensionality” definition of Pocrnic *et al*. 2016); the corresponding analyses are presented in the Results. The Jensen upper bound *U*_*B*_(*N*) on the Group B contribution (used in the closed-form expressions of Section 2.3) was computed from the same eigenvalue arrays via the formula given in Section S7.3. When *N* ≤ *M*_*e*_ the sample LD matrix has rank min(*N, M*) ≤ *M*_*e*_, so the zero-padded “Group B” eigenvalue slots contain no observable mass and both *S*_*B*_(*N*) and *U*_*B*_(*N*) evaluate numerically to zero; the population-level quantities are positive but cannot be estimated from samples in this regime.

To examine how the average GRAMMAR-Gamma coefficients change with sample size, we computed the full-GRM coefficient 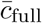 (the genome-wide average 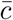 of Equation 4) and its LOCO analogue 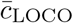 (the chromosome-wide average of the per-SNP *c*_LOCO_ of Equation 12) across a range of *N* by two different procedures. 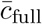 was computed directly from the LD-matrix eigenvalues via Equation 4: 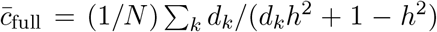, with the GRM eigenvalues *d*_*k*_ = *Nℓ*_*k*_*/M*. 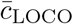 was computed by averaging *c*_LOCO_ over the SNPs of the focal chromosome (chromosome 1 was used throughout): explicitly, 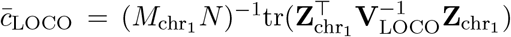), with 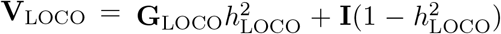 and the LOCO heritability set in proportion to the SNP count, 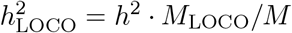, where *M*_LOCO_ is the number of SNPs outside the focal chromosome. The trace was evaluated by Cholesky decomposition with a dual trick: when *N* ≤ *M*_LOCO_ the *N* × *N* matrix **V**_LOCO_ was factorized directly; when *N* > *M*_LOCO_ the Woodbury identity reduced the problem to a Cholesky factorization of an *M*_LOCO_ × *M*_LOCO_ matrix. The LOCO upper bound 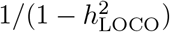 (Equation 12) was evaluated analytically at the corresponding 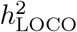. This analysis was carried out at *N* ∈ {5,000, 10,000, 20,000, 50,000, 100,000} for *N*_*e*_ = 20 and *N*_*e*_ = 50, and at *N* ∈ {5,000, 10,000, 20,000, 50,000} for *N*_*e*_ = 100; the upper *N* bound for *N*_*e*_ = 100 was set by the memory limits of our compute server (512 GB RAM), which became binding for the LOCO computation at *M* = 100,000 when *N* exceeded 50,000.

### 3.4 In-sample GBLUP reliability

We reused the simulated genotypes and phenotypes as described in Section 3.2 to validate the theoretical in-sample GBLUP reliability 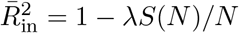 (Equation 17). For the scenarios with *β*^2^ = 0.001 (each of the five focal SNPs explaining 0.1% of phenotypic variance) across the 3 × 3 × 4 grid of *N*_*e*_, *h*^2^, and *N* (*N* = 5,000, 10,000, 20,000, and 50,000; 100 replicates each), the GREML model in SLEMM fitted the five focal SNPs as fixed effects and all *M* genome-wide SNPs as random effects. For each replicate, the prediction of an individual’s genetic value was the genotype-weighted sum of both the focal-SNP fixed-effect estimates and the genome-wide random SNP-effect estimates. The empirical in-sample reliability was the squared Pearson correlation between this prediction and the simulated true genetic value (averaged over the 100 replicates) and was compared with the theoretical 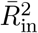, evaluated from 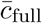 (calculated in Section 3.3) using 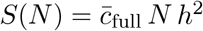 (Equation 6) at each simulated *h*^2^.

### 3.5 Closed-form practical detection and fine-mapping thresholds

#### Detection thresholds

We computed the practical detection threshold *q*_min_ ≈ 30*h*^2^*/M*_*e*_ (Equation 10; minimum per-SNP PVE for 50% power at the genome-wide significance threshold *χ*^2^ > 29.7, equivalently *p* < 5 × 10^−8^) at three working-model heritabilities *h*^2^ ∈ {0.1, 0.3, 0.5}. The transition scale 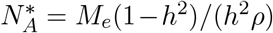 was evaluated at the same three *h*^2^ values, using *ρ* = 1 as a conservative simplification (the empirical spectra give *ρ* = 0.98 or 0.99). Species parameters follow Section S15 of the Supplementary Methods: cattle (*N*_*e*_ ≈ 100, *L* ≈ 25 Morgans, and *M*_*e*_ ≈ 10,000); pig (*N*_*e*_ ≈ 50, *L* ≈ 20 Morgans, and *M*_*e*_ ≈ 4,000); chicken (*N*_*e*_ ≈ 50, *L* ≈ 30 Morgans, and *M*_*e*_ ≈ 6,000); and human (*N*_*e*_ ≈ 10,000, *L* ≈ 35 Morgans, and *M*_*e*_ ≈ 1,400,000).

#### Fine-mapping thresholds

We computed the minimum PVE for fine-mapping resolution at inter-marker distances *d* ∈ {10, 100} kb (converted to Morgans via the standard 1 cM ≡ 1 Mb scaling) under two frameworks. Under the idealized simple-regression formula setting aside polygenic confounding (Section S13.3), 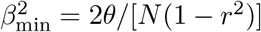 with 1 − *r*^2^ = 4*N*_*e*_*d/*(1 + 4*N*_*e*_*d*), evaluated at *N* = 100,000 and *θ* = 3 (Bayes factor ≈ 20). Under the full-GRM mixed model (Section S13.4), the asymptotic effect-size floor (per-SNP PVE) 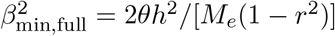 was evaluated at the same *θ* and *d*, and at the same three working-model heritabilities *h*^2^ ∈ {0.1, 0.3, 0.5}.

### 3.6 Software

Coalescent simulations used msprime v1.3.4 (Baumdicker *et al*. 2022). REML variance-component estimation and *χ*^2^ association testing in the phenotype simulations were performed with SLEMM (Cheng *et al*. 2023). PLINK v1.9 (Chang *et al*. 2015) was used for genotype-format conversion and basic QC. All other numerical analyses were performed in Python 3 with NumPy (linear algebra and exact eigendecomposition), scikit-learn (randomized SVD), pandas (tabular processing), SciPy (kernel density estimation), and Matplotlib (figure generation). Code and intermediate data are deposited as described in the Data Availability section.

## Results

### 4.1 Empirical support for the GRAMMAR-Gamma approximation

The per-SNP NCP formula (Equation 1) depends on the GRAMMAR-Gamma coefficient *c*_*l*_ (Equation 2), which varies across SNPs according to their projections onto the GRM eigenspace (Equation 3). The GRAMMAR-Gamma approximation replaces *c*_*l*_ with the genome-wide average 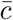 (Equation 4), converting the per-SNP power formula into the sigmoid sum *S*(*N*) (Equations 5–6). To assess whether this approximation is empirically justified, we computed *c*_*l*_ for every autosomal SNP in three publicly available livestock genotype datasets: Chinese Holstein cattle (*N* = 2,510 and *M* = 42,775 SNPs), Karacabey Merino sheep (*N* = 734 and *M* = 35,774 SNPs), and German Holstein cattle (*N* = 5,024 and *M* = 42,216 SNPs). For Chinese Holstein and Karacabey Merino, where chromosome maps were available, we additionally computed the LOCO coefficient *c*_LOCO_ by excluding the focal chromosome from the GRM for each SNP.

Figure 1 shows kernel density estimates of *c*_*l*_ at *h*^2^ = 0.1, 0.3, and 0.5 for each dataset, with the full-GRM coefficient on panels A–C and the LOCO coefficient on panels D–E. The distributions are concentrated around their means: the in-panel coefficient-of-variation (CV) annotations report values in the range 0.05–0.18 across all datasets and modes at *h*^2^ = 0.1–0.5, supporting the use of 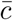 as a representative summary of *c*_*l*_ for standard livestock chip panels. The CV is consistently smaller for the LOCO mode than for the full-GRM mode at the same (dataset, *h*^2^) (for example, 0.09 versus 0.16 at *h*^2^ = 0.5 for Chinese Holstein), because the dominant component of *c*_LOCO_ comes from the subspace orthogonal to the LOCO GRM and is approximately constant across SNPs (Section 2.5, Equation 12). The CV also rises modestly with *h*^2^ in the full-GRM panels, reflecting the eigenvalue expansion (Equation 3): at higher *h*^2^, the shrinkage contrast between top and bottom eigenvectors is larger, so SNPs with different eigenspace projections experience more heterogeneous attenuation. Detailed summary statistics (mean, SD, CV, and 10th, 50th, 90th percentiles) for all panels are reported in Supplementary Table 1. Overall, these results show that SNP-to-SNP variation in *c*_*l*_ is modest for standard livestock chip panels.

**Figure 1.**
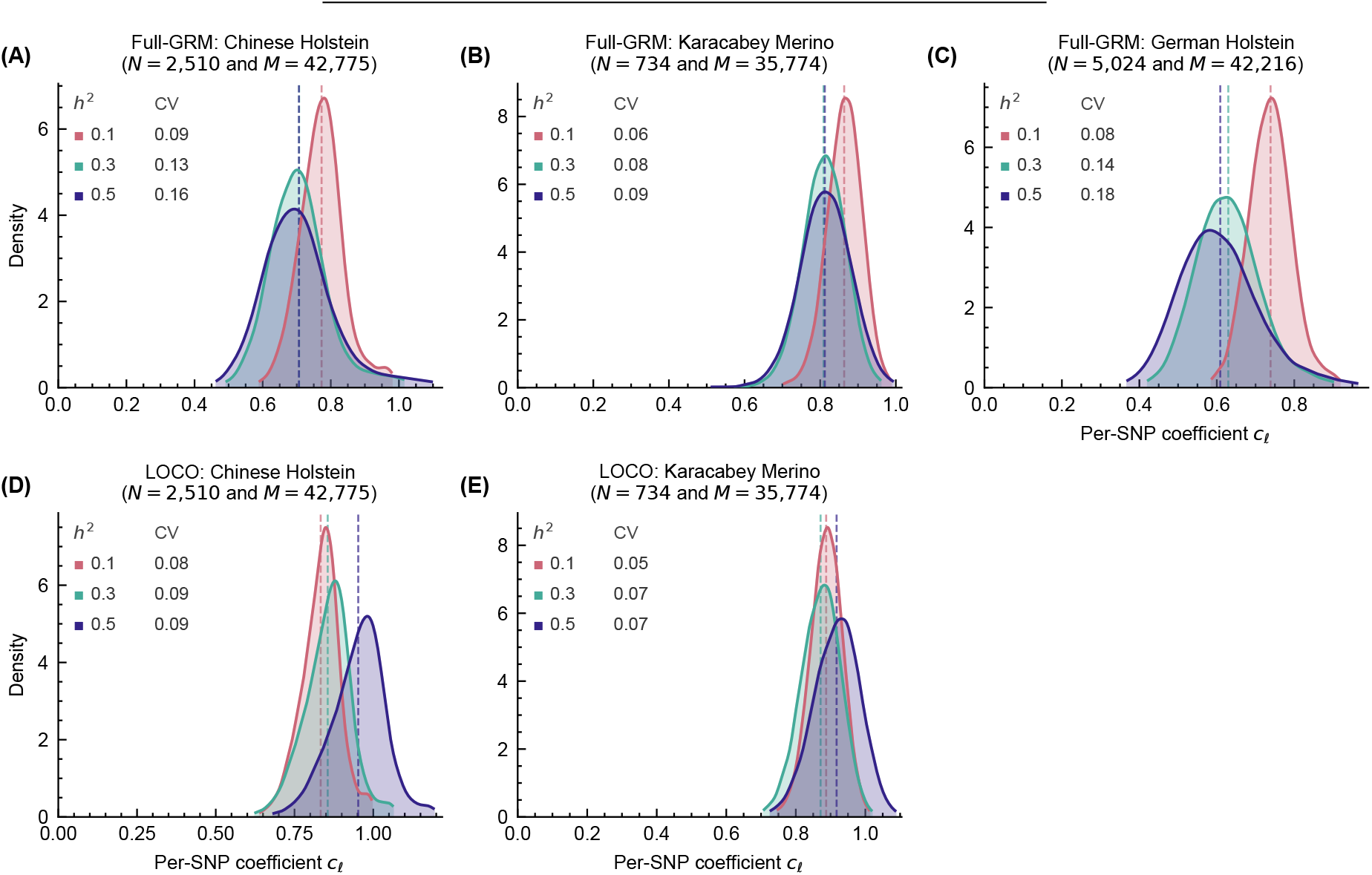
Distribution of the per-SNP GRAMMAR-Gamma coefficient *c*_*ℓ*_ in three livestock chip datasets. Kernel density estimates of *c*_*ℓ*_ across all autosomal SNPs, computed at heritability *h*^2^ = 0.1 (rose), 0.3 (teal), and 0.5 (indigo). (A–C) Full-GRM analyses in Chinese Holstein cattle (*N* = 2,510 and *M* = 42,775), Karacabey Merino sheep (*N* = 734 and *M* = 35,774), and German Holstein cattle (*N* = 5,024 and *M* = 42,216). (D, E) LOCO analyses in Chinese Holstein and Karacabey Merino; LOCO was not run on German Holstein because the dataset lacks per-SNP chromosome assignments. Vertical dashed lines mark the per-*h*^2^ mean (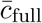 or 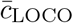); the inset annotation reports the coefficient of variation CV = SD*/*mean for each *h*^2^. Distributions are narrow (CV in the range 0.05–0.18 across all panels and heritabilities), supporting the GRAMMAR-Gamma approximation 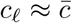 used in Equations 4–8. Summary statistics are reported in Supplementary Table 1.

### 4.2 Validation of the per-SNP NCP formula with phenotype simulations

To validate Equation 1 directly, we simulated phenotypes under the mixed model using coalescent-derived genotypes from msprime (v1.3.4; Baumdicker *et al*. 2022). We generated populations at *N*_*e*_ = 20, 50, and 100, each with a 25-Morgan genome (29 autosomes with cattle chromosome lengths), yielding *M* = 20,000 (*N*_*e*_ = 20), 50,000 (*N*_*e*_ = 50), and 100,000 (*N*_*e*_ = 100) segregating markers. For each population, we selected five focal SNPs spanning a wide allele-frequency spectrum (one per chromosome 1–5 at target minor allele frequencies of 0.075, 0.175, 0.275, 0.375, and 0.475) and simulated 100 replicate phenotypes at each combination of sample size (*N* = 5,000, 10,000, 20,000, and 50,000), heritability (*h*^2^ = 0.1, 0.3, 0.5), and focal-SNP proportion of phenotypic variance explained (*β*^2^ = 0.001 and 0.01). Each replicate used the full-GRM mixed model (GRM from all *M* SNPs, with the five focal SNPs included as fixed-effect covariates) and recorded the chi-squared statistic at each focal SNP.

Figure 2 compares the empirical NCP (mean *χ*^2^ − 1 across the 100 replicates per scenario) with the theoretical prediction from Equation 1 using the per-SNP coefficient *c*_*l*_. Across all 360 focal-SNP scenarios (3 *N*_*e*_ × 3 *h*^2^ × 2 *β*^2^ × 4 *N* × 5 SNPs), the per-SNP formula tracks the empirical NCP closely: the empirical-to-theoretical ratio has mean 1.03 and SD 0.08, and the 1:1 line falls within the point cloud throughout the NCP range from ~2 to ~250. Points are coded by color (*N*_*e*_) and marker shape (*h*^2^), showing that the agreement holds uniformly across all population structures, heritability levels, effect sizes, sample sizes, and minor allele frequencies. The full per-SNP table (Supplementary Table 2) lists each row’s *c*_*l*_, theoretical NCP, empirical NCP, and ratio. Combined with the empirical support in Section 4.1 that *c*_*l*_ varies only modestly around 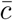, this result establishes the per-SNP NCP formula as the basis for the saturation argument in Section 4.3 (which uses 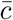).

**Table 2.** Detection and fine-mapping thresholds for livestock versus human GWAS. Closed-form predictions from the unified framework, with one column per species. Species parameters are *N*_*e*_ ≈ 100, *L* ≈ 25 M, and *M*_*e*_ ≈ 10,000 for cattle; *N*_*e*_ ≈ 50, *L* ≈ 20 M, and *M*_*e*_ ≈ 4,000 for pig; *N*_*e*_ ≈ 50, *L* ≈ 30 M, and *M*_*e*_ ≈ 6,000 for chicken; and *N*_*e*_ ≈ 10,000, *L* ≈ 35 M, and *M*_*e*_ ≈ 1,400,000 for human. For each *h*^2^ ∈ {0.1, 0.3, 0.5} the table reports the per-SNP practical detection floor *q*_min_ ≈ 30*h*^2^*/M*_*e*_ (Equation 10) and the transition sample size 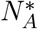. The simple-regression fine-mapping threshold 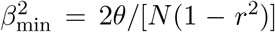 is reported at inter-marker distances *d* = 10 kb and *d* = 100 kb, evaluated at *N* = 100,000 and significance parameter *θ* = 3, with 1 − *r*^2^ = 4*N*_*e*_*d/*(1 + 4*N*_*e*_*d*) from the Sved (1971) 4*N*_*e*_*d* relation (with *d* as genetic distance in Morgans; the 10 and 100 kb values are converted via the standard 1 cM ≡ 1 Mb scaling). The full-GRM fine-mapping effect-size floor 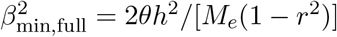 (Equation 16) is reported alongside for direct comparison.

| Parameter | Cattle | Pig | Chicken | Human |
| --- | --- | --- | --- | --- |
| $N_e$ | 100 | 50 | 50 | 10,000 |
| $L$ (Morgans) | 25 | 20 | 30 | 35 |
| $M_e$ | 10,000 | 4,000 | 6,000 | 1,400,000 |
| $q_{\min}$ (%), $h^2=0.1$ | 0.030 | 0.075 | 0.050 | 0.0002 |
| $q_{\min}$ (%), $h^2=0.3$ | 0.090 | 0.225 | 0.150 | 0.0006 |
| $q_{\min}$ (%), $h^2=0.5$ | 0.150 | 0.375 | 0.250 | 0.0011 |
| $N_A^*$, $h^2=0.1$ | 90,000 | 36,000 | 54,000 | 12,600,000 |
| $N_A^*$, $h^2=0.3$ | 23,333 | 9,333 | 14,000 | 3,266,667 |
| $N_A^*$, $h^2=0.5$ | 10,000 | 4,000 | 6,000 | 1,400,000 |
| Min PVE (%), simple regression, $d=10$ kb | 0.156 | 0.306 | 0.306 | 0.0075 |
| Min PVE (%), simple regression, $d=100$ kb | 0.021 | 0.036 | 0.036 | 0.0062 |
| Min PVE (%), full-GRM, $d=10$ kb ($h^2=0.1$) | 0.156 | 0.765 | 0.510 | $5.4 \times 10^{-5}$ |
| Min PVE (%), full-GRM, $d=100$ kb ($h^2=0.1$) | 0.021 | 0.090 | 0.060 | $4.4 \times 10^{-5}$ |
| Min PVE (%), full-GRM, $d=10$ kb ($h^2=0.3$) | 0.468 | 2.295 | 1.530 | 0.0002 |
| Min PVE (%), full-GRM, $d=100$ kb ($h^2=0.3$) | 0.063 | 0.270 | 0.180 | 0.0001 |
| Min PVE (%), full-GRM, $d=10$ kb ($h^2=0.5$) | 0.780 | 3.825 | 2.550 | 0.0003 |
| Min PVE (%), full-GRM, $d=100$ kb ($h^2=0.5$) | 0.105 | 0.450 | 0.300 | 0.0002 |

**Figure 2.**
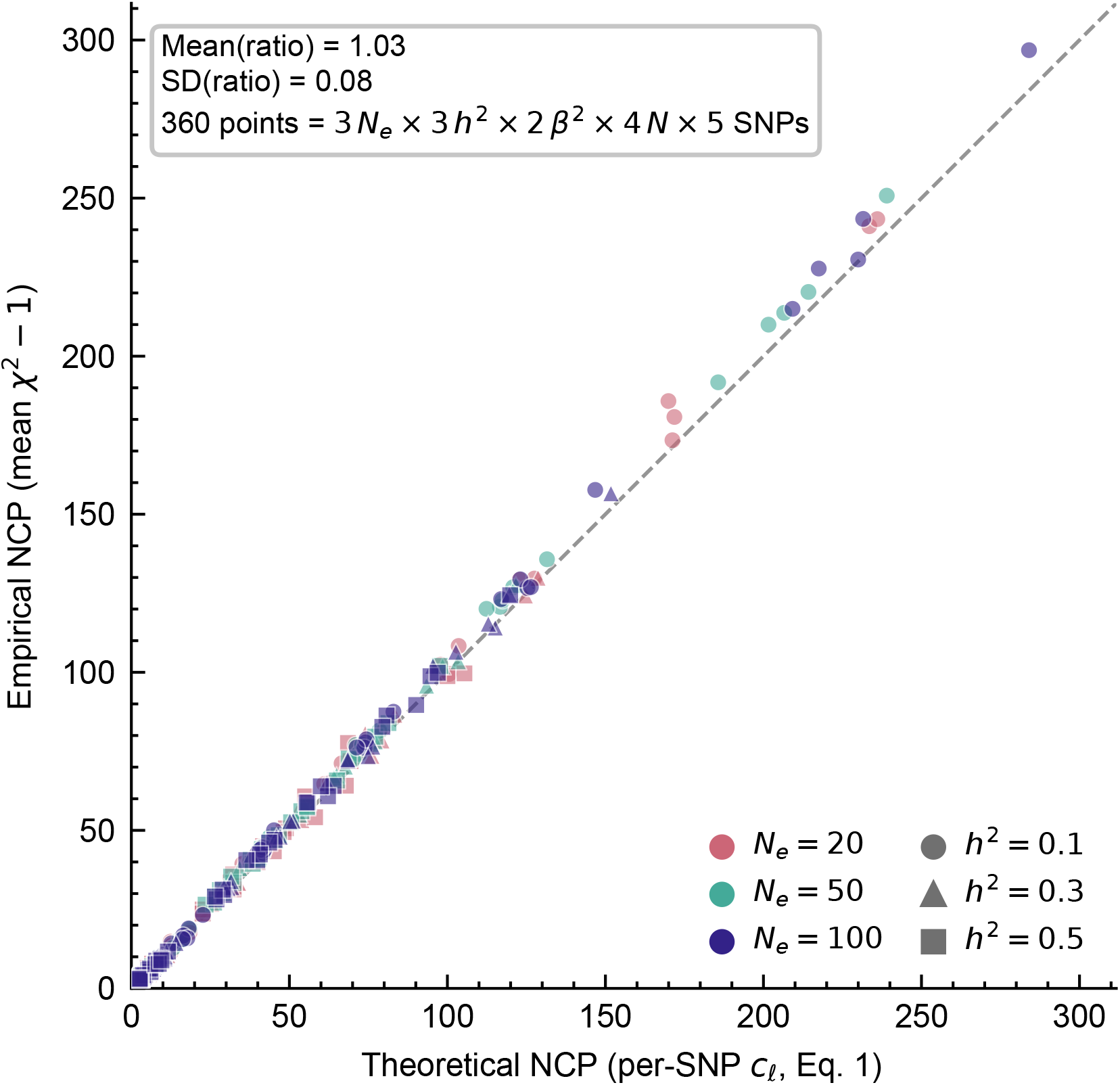
Theoretical versus empirical non-centrality parameter across 360 focal-SNP scenarios. Empirical NCP (mean score *χ*^2^ minus 1 across 100 replicates per scenario) plotted against the theoretical NCP computed from Equation 1 using the eigenvalue-specific coefficient *c*_*ℓ*_. Each point is one focal-SNP scenario: 3 *N*_*e*_ values × 3 *h*^2^ values × 2 *β*^2^ levels (0.001 and 0.01) × 4 sample sizes (*N* = 5,000, 10,000, 20,000, and 50,000) × 5 focal SNPs (one per chromosome 1–5, with target minor allele frequencies of 0.075, 0.175, 0.275, 0.375, and 0.475), for 360 points. Color encodes *N*_*e*_ (20, 50, 100); marker shape encodes *h*^2^ (circle 0.1, triangle 0.3, square 0.5). The dashed gray 1:1 line is shown for reference. The empirical-to-theoretical NCP ratio has mean 1.03 and SD 0.08 across all 360 scenarios, with the 1:1 line falling within the point cloud throughout the NCP range from ~2 to ~250. The close agreement across population structures, heritabilities, effect sizes, sample sizes, and minor allele frequencies validates the per-SNP NCP decomposition (Equation 1). Per-SNP values, including focal-SNP MAFs and *c*_*ℓ*_ values, are tabulated in Supplementary Table 2.

**Figure 3.**
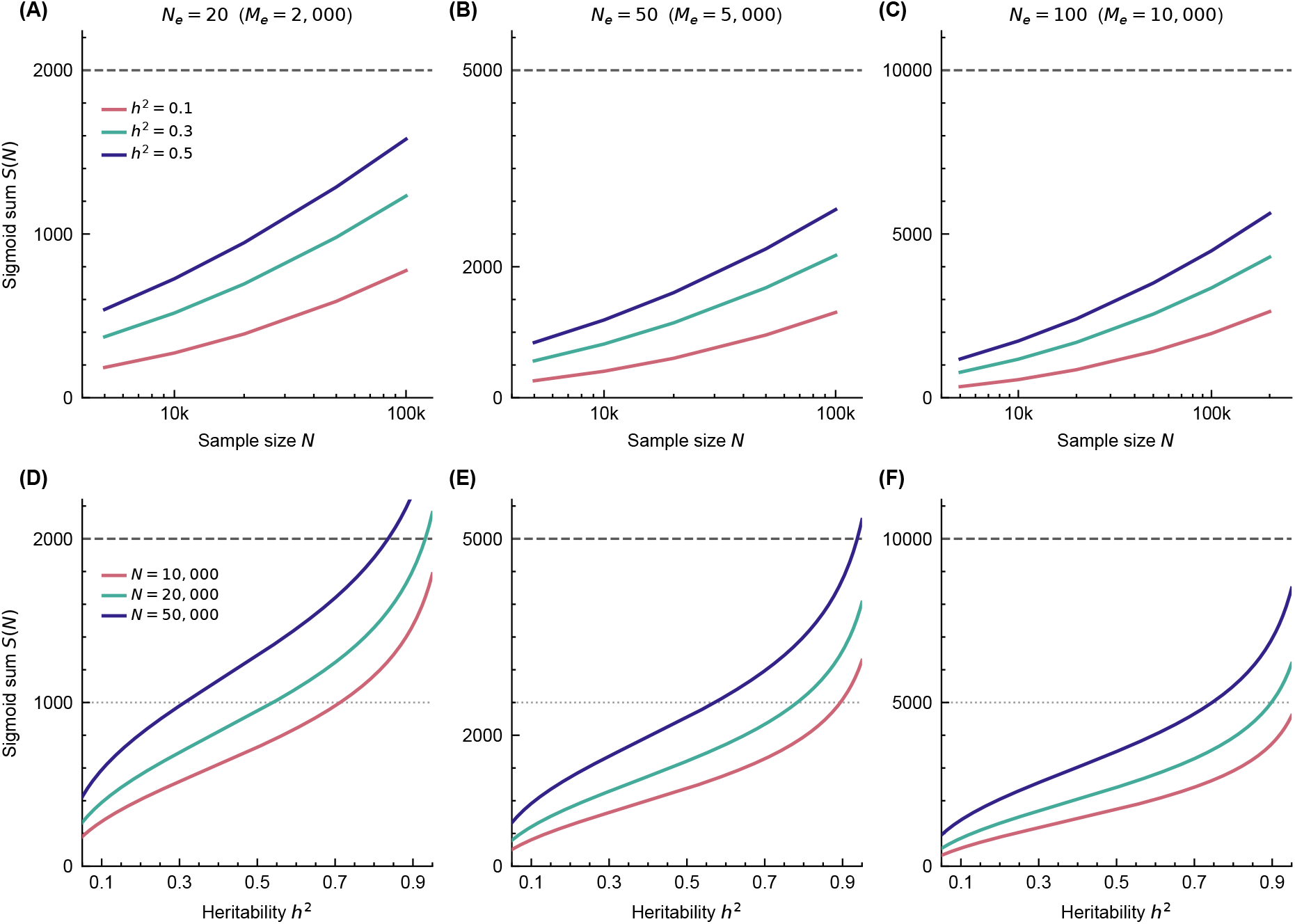
The sigmoid sum *S*(*N*) rises toward *M*_*e*_ = 4*N*_*e*_*L*, with (*N, h*^2^) exchangeable in *S*(*N*). Sigmoid sum *S*(*N*) = ∑_*k*_ *Nℓ*_*k*_*/*(*Nℓ*_*k*_ + *Mλ*) with *λ* = (1 − *h*^2^)*/h*^2^, evaluated using the *N*-specific LD-matrix eigenvalue spectrum computed from the msprime simulations at each sample size. Columns correspond to (A, D) *N*_*e*_ = 20, (B, E) *N*_*e*_ = 50, and (C, F) *N*_*e*_ = 100. The horizontal dashed line in every panel marks the practical ceiling *M*_*e*_ = 4*N*_*e*_*L* (*L* = 25 Morgans, the cattle genetic-map length used in all simulations). **Top row (A–C):** *S*(*N*) as a function of sample size *N* (log scale) for three heritability values, *h*^2^ = 0.1 (rose), 0.3 (teal), and 0.5 (indigo). Saturation more easily occurs at smaller *N*_*e*_ and larger *h*^2^, consistent with the transition-scale prediction 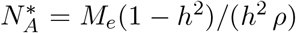. **Bottom row (D–F):** *S*(*N*) as a function of *h*^2^ over the broad range [0.05, 0.95] (covering both single-record heritabilities and possible DRP reliabilities), at three fixed sample sizes on the scale of livestock DRP cohorts: *N* = 10,000 (rose), *N* = 20,000 (teal), and *N* = 50,000 (indigo). A light gray dotted reference line at *M*_*e*_*/*2 aids reading the *N* –*h*^2^ trade-off: any fixed value of *S*(*N*) corresponds to a one-parameter family of (*N, h*^2^) pairs related by the equivalence *N h*^2^*/*(1 − *h*^2^) = constant (Section 2.4). For very small *N*_*e*_ at high *h*^2^ (visible in panels D and E), *S*(*N*) rises slightly above the practical ceiling *M*_*e*_: this reflects the non-negligible Group B contribution in the small-*N*_*e*_ regime (the hard ceiling is *M*, not *M*_*e*_; Section 2.3 and Supplementary Figure 1) and is a feature of the two-ceiling structure rather than a numerical artifact. Numerical *S*(*N*), *S*_*A*_(*N*), and *S*_*B*_(*N*) values are reported in Supplementary Table 3; eigenspectral parameters *ρ*, 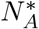, and 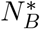 are reported in Table 1.

### 4.3 Concave approach of the sigmoid sum toward *M*_*e*_

The central prediction of the framework is that the sigmoid sum *S*(*N*) has a practical ceiling at *M*_*e*_ = 4*N*_*e*_*L*, approached at sample sizes well above the transition scale 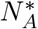 (Equations 5–6; Section 2.3). We tested this prediction using LD-matrix eigenvalue spectra computed from the coalescent genotypes at multiple sample sizes per population (each spectrum being a separate computation on *N* subsampled individuals), and we evaluated *S*(*N*) at each *N* using the corresponding *N*-specific spectrum.

Table 1 reports the theoretical *M*_*e*_ = 4*N*_*e*_*L*, the eigenvalue mass fraction *ρ* in the top *M*_*e*_ eigenvalues, and the transition scales 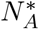 and 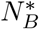 for each combination of *N*_*e*_ and three heritability values. The eigenvalue spectra used for Table 1 are at the largest available sample sizes for each *N*_*e*_ (*N* = 100,000 for *N*_*e*_ = 20 and 50; *N* = 200,000 for *N*_*e*_ = 100), so that the spectra are well-resolved. For all three populations, *ρ* ∈ [0.9981, 0.9986]: the top *M*_*e*_ eigenvalues account for 99.81–99.86% of the total spectral mass, leaving 0.14–0.19% in the remaining *M*_*B*_ = *M* − *M*_*e*_ near-zero modes. The ratio 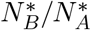 ranges from 4,780 to 6,346, confirming that the two transition scales are separated by several orders of magnitude. Supplementary Tables 6 and 7 report the analogous parameters when *M*_*e*_ is defined data-drivenly as the number of top eigenvalues capturing 98% or 99% of the spectral mass; in both alternatives, the qualitative two-ceiling structure is preserved.

The top row of Figure 3 shows *S*(*N*) computed from per-*N* eigenvalue spectra as a function of *N* (log scale) for each combination of *N*_*e*_ and *h*^2^ (values in Supplementary Table 3). In every panel, *S*(*N*) rises with *N* toward the *M*_*e*_ ceiling (dashed horizontal line) but has not reached it: at the largest simulated *N, S*(*N*)*/M*_*e*_ ranges from ~0.26 (*N*_*e*_ = 100 and *h*^2^ = 0.1) to ~0.79 (*N*_*e*_ = 20 and *h*^2^ = 0.5). The curves rise more slowly with each additional decade of *N* at the higher end of the simulated range, indicating the diminishing-returns phase characteristic of approach to the ceiling. The Group B component *S*_*B*_(*N*) remains a small fraction of *S*_*A*_(*N*) throughout the practical regime (Supplementary Figure 1): *S*_*B*_*/S*_*A*_ grows with both *h*^2^ and *N*, reaching a peak of 0.11 for *N*_*e*_ = 20 at *h*^2^ = 0.5 and *N* = 100,000, and staying at or below 0.06 for *N*_*e*_ = 50 and *N*_*e*_ = 100 across the simulated grids. In all cases *S*_*B*_*/S*_*A*_ is small enough that the practical ceiling is set by the top *M*_*e*_ eigenmodes. The Jensen upper bound *U*_*B*_(*N*) used in the derivation tracks *S*_*B*_(*N*) closely (Supplementary Figure 2), validating the use of *U*_*B*_(*N*) as a proxy for *S*_*B*_(*N*) in the closed-form expressions.

The bottom row of Figure 3 shows the complementary cross-section: *S*(*N*) plotted against *h*^2^ over the broad range [0.05, 0.95] at three fixed sample sizes (*N* = 10,000, 20,000, and 50,000). Across all simulated (*N*_*e*_, *N*) combinations, *S*(*N*) rises substantially with *h*^2^: increasing *h*^2^ from 0.05 to 0.9 multiplies *S*(*N*) by approximately five-to eleven-fold (Supplementary Table 3). The leverage is largest at smaller *N*, where *S*(*N*) remains far from *M*_*e*_ at moderate *h*^2^. For very small *N*_*e*_ at high *h*^2^ (e.g., panel D at *N* = 50,000 and *h*^2^ = 0.9: *S*(*N*)*/M*_*e*_ ≈ 1.14), *S*(*N*) rises above *M*_*e*_ via the non-negligible Group B contribution, a manifestation of the two-ceiling structure rather than a numerical artifact (Section 2.3 and Supplementary Figure 1). The *h*^2^ leverage provides the theoretical basis for DRP-pseudo-phenotype GWAS (an example discussed in Section 5.5): raising the effective *h*^2^ from a single-record value to a DRP reliability substantially amplifies per-SNP detection power at fixed *N*.

The saturation of *S*(*N*) as *N* grows at fixed *h*^2^ ties to the declining full-GRM average coefficient 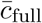 (the 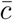 of Equation 4), since 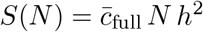 (Equation 6). Figure 4 shows 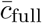 and 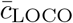 (its LOCO analogue) computed from coalescent-simulated genotypes (using the same per-*N* eigenvalue spectra as in Figure 3) across sample sizes *N* = 5,000 to 100,000 for *N*_*e*_ = 20 and *N*_*e*_ = 50, and *N* = 5,000 to 50,000 for *N*_*e*_ = 100 (the LOCO Cholesky becomes memory-prohibitive at larger *N* when *M* = 100,000; Methods §3.3). In every panel, 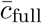 decreases monotonically with *N*, reflecting the progressive absorption of focal-SNP signal by the GRM random effects. For example, at *N*_*e*_ = 50 and *h*^2^ = 0.3, 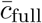 drops from 0.38 (*N* = 5,000) to 0.07 (*N* = 100,000), a five-fold decrease. In contrast, 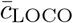 remains nearly constant or increases slightly, approaching the theoretical upper bound 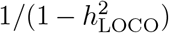 (dotted lines) as *N* grows: at *N*_*e*_ = 50 and *h*^2^ = 0.3, 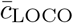 rises from 1.25 to 1.36, approaching its bound of 1.39. This divergence between 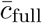 and 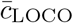 explains the empirical observation that GWAS power saturates under full-GRM analysis at large *N* while LOCO power continues to grow linearly with sample size (Section 2.5). The complete tabulated values of 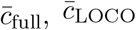, and the bound at every simulated *N* are in Supplementary Table 4.

**Figure 4.**
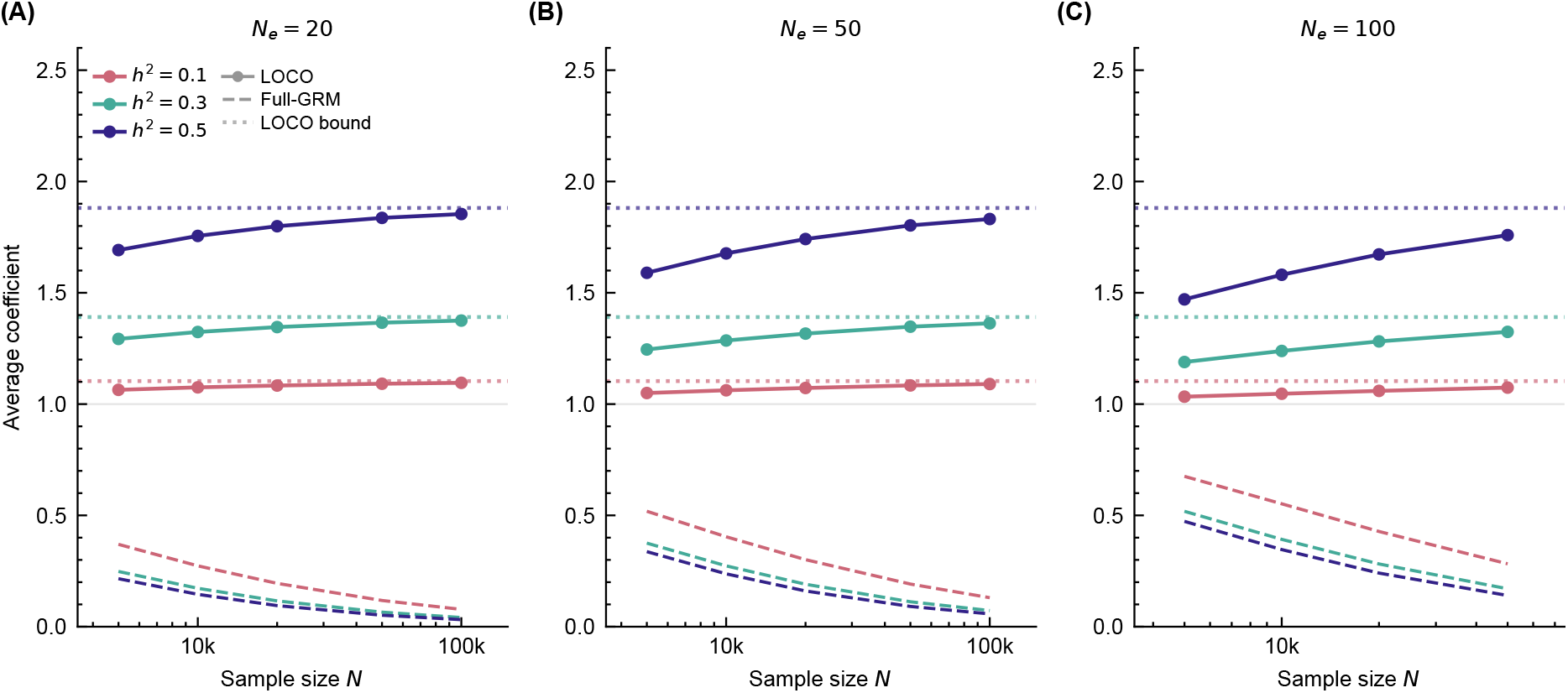
Full-GRM and LOCO average GRAMMAR-Gamma coefficients diverge with sample size. Average coefficients 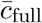 (dashed lines) and 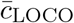 (solid lines with markers) plotted against sample size *N* (log scale), with one panel per population: (A) *N*_*e*_ = 20, (B) *N*_*e*_ = 50, and (C) *N*_*e*_ = 100. Plotted range is *N* = 5,000 to 100,000 for *N*_*e*_ = 20 and *N*_*e*_ = 50, and *N* = 5,000 to 50,000 for *N*_*e*_ = 100 (the LOCO Cholesky factorization becomes memory-prohibitive at larger *N* when *M* = 100,000; Section 3.3). Colors encode heritability (*h*^2^ = 0.1 rose, 0.3 teal, and 0.5 indigo). The thin gray horizontal line at *y* = 1 marks the full-GRM / LOCO boundary; the per-*h*^2^ asymptotic upper bound for 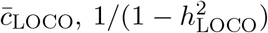 (Equation 12), is shown as a thin horizontal dotted line in the corresponding *h*^2^ color. 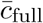 decreases monotonically with *N*, consistent with *S*(*N*) → *M*_*e*_ and 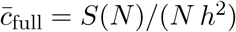(Equation 6); in contrast, 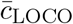 remains nearly constant or rises slightly toward its upper bound, reflecting the fact that LOCO does not saturate at the same ceiling because the focal chromosome is excluded from the GRM (Section 2.5). The divergence widens with *h*^2^ and is larger at smaller *N*_*e*_. Underlying values are reported in Supplementary Table 4.

The practical consequence of these results is that, in livestock populations with small *N*_*e*_ and hence small *M*_*e*_, increasing sample size beyond 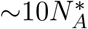 yields rapidly diminishing returns in full-GRM association power, a saturation pattern that does not appear in mixed-model GWAS of large-*N*_*e*_ species such as humans because *M*_*e*_ is orders of magnitude larger there.

### 4.4 Genomic prediction reliability

The sigmoid sum *S*(*N*) that governs full-GRM association power also sets the in-sample GBLUP reliability, 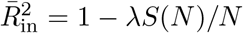 (Equation 17; Sections 2.7 and S10). We validated this equation directly with the phenotype simulations of Section 4.2. For the scenarios in which each focal SNP explained 0.1% of phenotypic variance (*β*^2^ = 0.001), we computed the empirical in-sample reliability as the squared correlation between the prediction and the true genetic value, averaged over 100 replicates, and compared it with the predicted 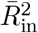 across all 36 (*N*_*e*_, *h*^2^, *N*) combinations, with *N* = 5,000, 10,000, 20,000, and 50,000 (Supplementary Table 8). The empirical and theoretical reliabilities, which span 0.40 to 0.97, agreed closely, with a mean ratio of 1.00 (Figure 5). This agreement held even though the five focal SNPs were fitted as fixed effects, a departure from the infinitesimal genetic architecture assumed by Equation 17. The theoretical reliability prediction is thus robust to this degree of genetic-architecture misspecification.

**Figure 5.**
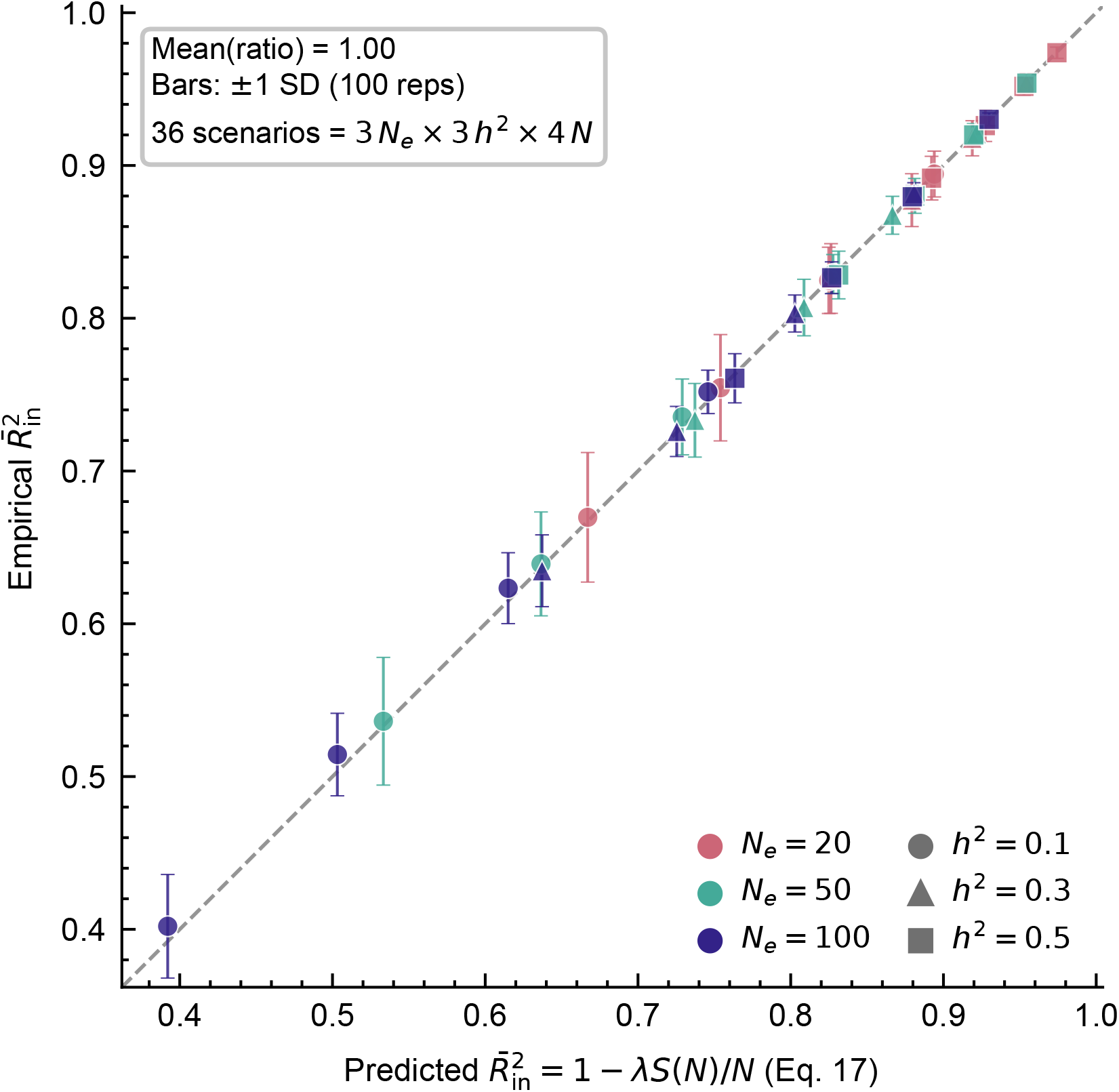
The predicted in-sample GBLUP reliability matches the empirical reliability across simulated scenarios. Each point is one (*N*_*e*_, *h*^2^, *N*) scenario from the phenotype simulations with *β*^2^ = 0.001 (each of the five focal SNPs explaining 0.1% of phenotypic variance). The empirical in-sample reliability (the *y* axis) is the squared Pearson correlation between the prediction and the true genetic value, averaged over 100 replicates; vertical bars give ±1 standard deviation across these replicates. The *x* axis shows the predicted 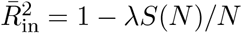 (Equation 17). Color encodes *N*_*e*_ (20 rose, 50 teal, and 100 indigo) and marker shape encodes heritability (*h*^2^ = 0.1 circle, 0.3 triangle, and 0.5 square); the four points sharing a color and marker are the sample sizes *N* = 5,000, 10,000, 20,000, and 50,000. The dashed gray line is the 1:1 identity. Across all 36 scenarios the mean empirical/predicted ratio is 1.00. Underlying values are reported in Supplementary Table 8.

### 4.5 Practical detection and fine-mapping thresholds

The theory yields closed-form expressions for the minimum detectable proportion of variance explained (PVE) at the practical full-GRM NCP ceiling (Equation 10) and the minimum resolvable effect size for fine-mapping at a given genomic distance (Section 2.6). We computed these thresholds for three livestock species representing a range of *N*_*e*_ and genome lengths, and compared them with human GWAS parameters. Species parameters follow Section S15 of the Supplementary Methods: cattle (*N*_*e*_ ≈ 100, *L* ≈ 25 Morgans, and *M*_*e*_ ≈ 10,000), pig (*N*_*e*_ ≈ 50, *L* ≈ 20 Morgans, and *M*_*e*_ ≈ 4,000), chicken (*N*_*e*_ ≈ 50, *L* ≈ 30 Morgans, and *M*_*e*_ ≈ 6,000), and human (*N*_*e*_ ≈ 10,000, *L* ≈ 35 Morgans, and *M*_*e*_ ≈ 1,400,000). All results are collected in Table 2.

#### Detection thresholds

For cattle, *q*_min_ ≈ 30 *h*^2^*/M*_*e*_ (Equation 10) gives *q*_min_ ≈ 0.090% at *h*^2^ = 0.3: under the practical full-GRM NCP ceiling, variants below this PVE are below the approximate 50%-power detection floor for genome-wide significance (*p* < 5 × 10^−8^). For pig, *q*_min_ ≈ 0.225% at *h*^2^ = 0.3; for chicken, *q*_min_ ≈ 0.150%. These thresholds are readily interpretable: large-effect QTL such as *DGAT1* in dairy cattle (PVE ~ 5%) are easily detectable, but moderate-effect loci explaining <0.1% of phenotypic variance are below the practical detection floor. The corresponding transition scale 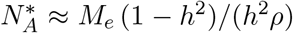, above which power growth slows toward the ceiling, is 23,333 (cattle), 9,333 (pig), and 14,000 (chicken) at *h*^2^ = 0.3 (well within current large-scale livestock GWAS sample sizes), and rises to 90,000, 36,000, and 54,000, respectively, at *h*^2^ = 0.1. The human threshold (*q*_min_ ≈ 6 × 10^−4^% at *h*^2^ = 0.3) is orders of magnitude lower than the livestock values, and 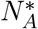 exceeds current GWAS sample sizes (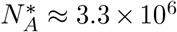 at *h*^2^ = 0.3), so human mixed-model power continues to grow approximately linearly with *N* at all currently feasible sample sizes. This explains why the full-GRM power saturation phenomenon is specific to livestock: the same mixed-model method behaves very differently in species with small *N*_*e*_ (livestock) versus large *N*_*e*_ (humans).

#### Fine-mapping resolution under the idealized simple-regression formula

Setting aside polygenic confounding, the minimum effect size (per-SNP PVE) resolvable at genetic distance *d* (in Morgans) from a focal QTL in a sample of *N* individuals is 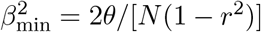 (Equations 14–15), with 1 − *r*^2^ = 4*N*_*e*_*d/*(1 + 4*N*_*e*_*d*) (Sved 1971). The resolution improves indefinitely with *N*. Evaluated at *N* = 100,000 and a Bayes-factor threshold *θ* = 3 (with physical distance converted to Morgans via the standard 1 cM ≡ 1 Mb scaling), the minimum resolvable effect size at 10 kb is 0.156% in cattle (1 − *r*^2^ ≈ 0.038), 0.306% in pig and chicken (1 − *r*^2^ ≈ 0.020), and 0.0075% in humans (1 − *r*^2^ ≈ 0.80). At 100 kb, the livestock thresholds drop to 0.021–0.036%, suggesting that, under this idealized analysis, 100-kb-level localization is achievable for moderate-to-large-effect QTL in livestock. Fine-mapping resolution depends on *N*_*e*_ through the LD decay function but not on genome length *L*, so pig and chicken share identical fine-mapping thresholds. We note that this idealized analysis has its own intrinsic limit: polygenic block signals (in which all SNPs in a region carry similar small effects) cannot be successfully fine-mapped at any *N* with genotype-phenotype associations alone, because the expected log-likelihood difference between within-block SNPs is proportional to the per-SNP effect size *β*^2^, which is negligible under diffuse polygenic effects (Section 2.6; Section S13.2).

#### Fine-mapping under the full-GRM mixed model

Once polygenic confounding is explicitly modeled through a GRM (typically constructed from chip SNPs), fine-mapping evidence saturates as *N* grows: the expected log-likelihood difference 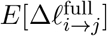 plateaus at *M*_*e*_*β*^2^(1 − *r*^2^)*/*(2*h*^2^) (Section S12.3), giving a floor on resolvable PVE, 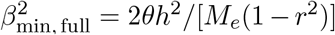, that no further increase in *N* can practically lower. At *h*^2^ = 0.3 and 10 kb, the floor is 0.47% for cattle, 2.30% for pig, and 1.53% for chicken, three to seven times the corresponding simple-regression threshold at *N* = 100,000. The floor scales linearly in *h*^2^, so the corresponding values at *h*^2^ = 0.1 are one-third and at *h*^2^ = 0.5 are five-thirds of these. For humans the floor is 0.0002% at 10 kb (at *h*^2^ = 0.3), comfortably below the simple-regression value because *M*_*e*_*/h*^2^ is far larger than 10^5^.

#### Unified effect-size floor

The simple-regression and full-GRM fine-mapping analyses converge on the same conclusion. Whether we evaluate the simple-regression threshold at currently feasible *N* or the full-GRM effect-size floor at arbitrarily large *N*, livestock fine-mapping requires effect sizes of similar or higher order of magnitude than the practical GWAS detection threshold *q*_min_ (typically around tenths of a percent of phenotypic variance) because all three quantities scale through *N*_*e*_: the detection threshold *q*_min_ and the full-GRM fine-mapping effect-size floor through *M*_*e*_ = 4*N*_*e*_*L*, and both fine-mapping thresholds also through the 4*N*_*e*_*d*-scaled LD-decay term 1 − *r*^2^ at the inter-marker distance *d*. A variant that is marginally detectable at the practical full-GRM NCP ceiling is generally not fine-mappable to a high resolution of 10 kb either. In humans, the much larger *M*_*e*_ (1.4 × 10^6^) together with the much weaker LD (1 − *r*^2^ approaching 1 at typical fine-mapping distances) makes both the detection and fine-mapping thresholds orders of magnitude lower, so neither poses a practical barrier at currently achievable sample sizes. This unified effect-size floor in livestock thus arises directly from the small *N*_*e*_.

## Discussion

### 5.1 A unified framework anchored on *N*_*e*_

We have shown that a single population parameter, the effective population size *N*_*e*_, governs three classical applications in livestock quantitative genetics through two related quantities: the effective number of independent chromosome segments *M*_*e*_ = 4*N*_*e*_*L* (Sved 1971; Stam 1980) and the 4*N*_*e*_*d*-scaled LD decay (Sved 1971) at inter-marker distance *d*. For genomic prediction, *M*_*e*_ has long been recognized as the dimensionality limit beyond which marker density yields diminishing returns (Goddard 2009; Pocrnic *et al*. 2016). We show that the same sigmoid sum *S*(*N*) that scales GWAS detection power (Equations 5 and 7) also sets prediction reliability: the in-sample GBLUP reliability 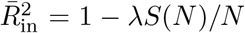 (Equation 17) and the leave-one-out reliability bounds (Equation 19) both depend on *S*(*N*) (Section 2.7). For mixed-model GWAS, this work establishes *M*_*e*_ as the practical ceiling of *S*(*N*) in the full-GRM NCP, approached at sample sizes much larger than the transition scale 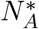, and the per-SNP detection floor is *q*_min_ ≈ 30*h*^2^*/M*_*e*_ at 50% power (Equation 10). Because *S*(*N*) is common to detection power, in-sample reliability, and the leave-one-out bounds, its practical ceiling *M*_*e*_ bounds all three. For genetic fine-mapping, *N*_*e*_ appears in two places: through 4*N*_*e*_*d*-scaled LD decay, which governs the within-block resolution under any framework (linear regression, LOCO, or full-GRM mixed model), and additionally through *M*_*e*_ in the full-GRM mixed-model fine-mapping effect-size floor (Section 4.5). The framework also makes a clean prediction about LOCO: by excluding the focal chromosome from the GRM, LOCO bypasses the practical full-GRM NCP ceiling but at the cost of detecting block-level rather than SNP-level signals (Section 2.5). The numerical implications for cattle, pig, and chicken are quantified in Table 2 and contrasted with human GWAS, where *M*_*e*_ is two to three orders of magnitude larger and the full-GRM NCP ceiling does not bind at currently feasible sample sizes.

#### Quantitative consistency check from external data

The simulation study by Jang *et al*. (2023), which examined GWA performance across *N*_*e*_ (20 and 200) and trait heritability in a cattle-genome simulation, provides an external quantitative test. We focus on the more polygenic Q2000 scenarios (2,000 quantitative trait nucleotides; QTN) at *N* = 30,000 and high heritabilities *h*^2^ ∈ {0.9, 0.99}, where the practical-saturation condition 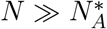 holds (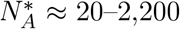 across the four (*N*_*e*_, *h*^2^) combinations). Under *S*(*N*) ≈ *M*_*e*_, Equation 7 reduces to NCP ≈ *M*_*e*_ · *q/h*^2^. Because the per-QTN PVE in Jang *et al*.’s simulation scales as *q* ∝ *h*^2^ (gamma-distributed effects rescaled so total genetic variance matches the simulated *h*^2^), NCP and hence the expected count of detected QTN at genome-wide significance (*p <* 10^−7^, equivalently *χ*^2^ *>* 28.374) are independent of *h*^2^ under this saturated formula. Summing 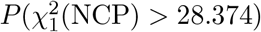 across the 2,000 QTN, with *M*_*e*_ approximated by Jang *et al*.’s EIG99 estimates (6,774 at *N*_*e*_ = 20; 21,177 at *N*_*e*_ = 200), predicts approximately 61 and 186 detected QTN at *N*_*e*_ = 20 and 200, respectively. These predictions broadly match the empirical counts (Jang *et al*. 2023, Tables S1b and S1d): 46 and 117 at *h*^2^ = 0.9, and 138 and 218 at *h*^2^ = 0.99. Different choices of *M*_*e*_ (EIG98, EIG99, or theoretical 4*N*_*e*_*L*) shift the predicted counts, but the broad consistency with empirical counts is preserved. The underestimation in some scenarios (notably *N*_*e*_ = 20 and *h*^2^ = 0.99) reflects *S*(*N*) passing beyond the practical ceiling *M*_*e*_ via Group B eigenmode contributions. This impractical regime (*N* = 30,000 combined with *h*^2^ = 0.99) is anticipated by the two-ceiling structure (Sections 2.3 and S8).

### 5.2 What full-GRM mixed-model GWAS actually detects

A direct consequence of the *M*_*e*_ ceiling is that full-GRM mixed-model GWAS in livestock can identify only a limited number of association peaks on a Manhattan plot, regardless of how large the sample becomes. This limit is a consequence of how the test is constructed: it detects significant *deviations from the polygenic background* that the GRM has already absorbed. Under a small-*N*_*e*_ population the polygenic background has its LD-patterned variance (variance aligned with the GRM eigenspace) concentrated within the top *M*_*e*_ eigenmodes of the GRM (Sections 2.3 and 4.3), so what remains after GRM adjustment are precisely the variants whose SNP-level effects exceed what diffuse polygenicity alone would predict. In livestock, such variants are typically few in number. The same finding has been reported across multiple large livestock GWAS: a full-GRM mixed-model GWAS of 1.16 million Holstein cattle (Jiang *et al*. 2022); a sequence-variant analysis of ~50,000 Holstein bulls covering 30 dairy traits, using highly reliable de-regressed breeding values as pseudo-phenotypes (Wang *et al*. 2025); and a single-step GWAS of hundreds of thousands of pigs from purebred maternal lines (Kayondo *et al*. 2026), an APY-based ssGWAS analogous to full-GRM mixed-model association (Leite *et al*. 2024). All return a small set of peaks.

Pocrnic *et al*. (2024) reached a complementary conclusion from a different starting point. Examining the SNP-effect “profile” around true causal variants in small-*N*_*e*_ populations, they showed that the profile is wide rather than narrow because the LD-driven signal spreads across many in-block variants, that the number of QTL with truly large per-SNP effects is small in a small-*N*_*e*_ population, and that identification of the actual causal variant is therefore hard. Their SNP-profile perspective and the present test-statistic perspective triangulate the same *M*_*e*_ ceiling from different directions. The contribution of the present work is to derive the *mechanism* that generates that ceiling at the level of the statistical model: the per-SNP NCP decomposition (Equation 1), the sigmoid-sum saturation *S*(*N*) → *M*_*e*_ (Section 2.3), and the GRAMMAR-Gamma approximation that links the two. The mechanism then yields the closed-form detection threshold *q*_min_ ≈ 30*h*^2^*/M*_*e*_ and the full-GRM fine-mapping effect-size floor that Pocrnic *et al*. (2024)’s SNP-profile view does not directly produce.

Empirically, alternative methods to full-GRM produce many more significant associations under the same data. Even under fully diffuse genetic architectures, in which each of 10,000 causal variants contributes only an individually tiny effect, methods such as LOCO and fastGWA can flag many seemingly strong associations spanning long LD blocks (Wang *et al*., in preparation). These additional “associations” do not reflect newly discovered large-effect variants; they reflect aggregate LD-weighted effects of diffuse polygenic background that the full-GRM model deliberately absorbs.

### 5.3 Comparison with alternative mixed-model methods

The distinct behavior of full-GRM, LOCO, fastGWA, and pedigree-based mixed models in livestock is explained naturally within our framework. LOCO removes the focal chromosome from the GRM, which allows focal-chromosome association statistics to continue gaining power as sample size grows, but the resulting signals are block-level aggregates rather than SNP-level excess effects (Section 2.5 and Section S5). Pedigree mixed-model GWAS uses the pedigree-based numerator relationship matrix as the random-effect covariance and therefore cannot capture the Mendelian-sampling component of genetic variance that distinguishes individuals from their pedigree relatives, approximately half of the additive genetic variance. A substantial fraction of LD-patterned genetic signal therefore escapes the random-effect term and is partially picked up by the SNP being tested; the qualitative behavior resembles LOCO but with somewhat less inflation (Jiang *et al*. 2019a). fastGWA adopts a sparse GRM by zeroing entries below a small threshold (e.g., 0.05), which approximates the pedigree relationship matrix and is therefore conceptually analogous to a pedigree mixed model in its coverage of relatedness. APY (Misztal *et al*. 2014) is a separate case discussed below.

A central observation is that these methods behave very differently across species. In humans, where *M*_*e*_ ≈ 1.4 × 10^6^ generally exceeds currently feasible sample sizes, the full-GRM mixed model never approaches its *M*_*e*_ ceiling; LOCO and fastGWA results are largely the same as the full-GRM result (Loh *et al*. 2015; Zhou *et al*. 2018; Jiang *et al*. 2019b). In livestock, where *M*_*e*_ is two to three orders of magnitude smaller, full-GRM can hit the NCP ceiling within currently achievable *N*, and the three methods can return strikingly different Manhattan plots. The driving factor is *N*_*e*_. This species-by-method divergence is not unique to GWAS: methods originally calibrated for humans, such as LD-score regression and Haseman–Elston regression for heritability estimation and partitioning, also perform poorly in livestock for the same reason and require alternative approaches calibrated to livestock populations (Jiang 2024).

The Algorithm for Proven and Young (APY; Misztal *et al*. 2014), originally introduced for single-step GBLUP and now also used in single-step GWAS (Leite *et al*. 2024), can be viewed within the same framework. APY approximates the inverse of the GRM by partitioning genotyped animals into a “core” of size *n*_*c*_ and a “non-core” set whose breeding values, conditional on the core breeding values, are modeled as mutually independent with a diagonal residual covariance. Pocrnic *et al*. (2016b) showed empirically that prediction accuracy plateaus when *n*_*c*_ ≈ *M*_*e*_, a result that has guided practical core-size selection. Our framework provides a rationale: APY’s rank-*n*_*c*_ factor structure with diagonal residual absorbs LD-patterned signal up to dimension *n*_*c*_, largely matching the *M*_*e*_ ceiling. For ssGWAS, this means APY inherits the practical full-GRM detection threshold *q*_min_ when *n*_*c*_ ≈ *M*_*e*_; the ceiling is converted from a soft plateau into a hard ceiling at *n*_*c*_ because the diagonal-residual block cannot recover residual LD-patterned mass at any *N*, but in livestock applications where currently feasible *N* is far below 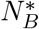 this distinction is operationally invisible (Leite *et al*. 2024).

### 5.4 The implicit goal of GWAS and why full-GRM is the right tool for livestock

Given the mechanistic contrast above, whether the desired outcome is a few peaks or many depends on what GWAS is for. Common downstream practices reveal an implicit assumption: candidate-gene lookup near peaks, sparse-prior fine-mapping, and colocalization with molecular QTL all assume that the underlying biological signal is sparse and individually identifiable. Block-level polygenic associations do not facilitate any of these analyses, and can actively mislead them: diffuse polygenic effects across a long LD block are by construction not localizable to a candidate gene within that block, and fine-mapping a block whose true architecture is uniformly small effects yields spurious “credible sets” that mix hundreds of variants of essentially equal evidence.

A second, more fundamental point is that the additional block-level “associations” reported by alternative methods (LOCO, fastGWA, and related approaches) do not represent new biological knowledge. Widespread polygenicity of complex traits in livestock has long been a working assumption: formalized in the Fisher infinitesimal model (Fisher 1918), refined into the omnigenic perspective (Boyle *et al*. 2017), and corroborated in practice by the successful implementation of genomic selection using moderate-density SNP chips (García-Ruiz *et al*. 2016). The existence of polygenic effects is settled science; what livestock GWAS contributes that *cannot* be obtained from the polygenic view is the identification of variants whose individual effects are strong enough for follow-up variant- or gene-targeted studies. One might counter that block-level associations could still feed downstream summary-statistic methods such as stratified LD-score regression for partitioned heritability or functional-category enrichment. In livestock, however, these methods themselves perform poorly for the same *M*_*e*_-related reasons (Jiang 2024), so block-level associations in livestock support neither variant-level follow-up nor reliable partitioned-heritability analysis. The full-GRM mixed model is specifically calibrated for this task: it filters out block-level polygenic signal so that the surviving peaks correspond to variants of genuinely above-background effect.

Furthermore, even if one accepts that block-level polygenic associations are an interesting endpoint in themselves, the unified effect-size floor we derive (Section 4.5) implies that diffuse polygenic effects within a block are impractical to fine-map with genotype-to-phenotype association frameworks in livestock at any feasible sample size, because the per-variant 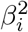 is too small to clear either the simple-regression or the full-GRM resolution threshold. The block-level associations are therefore not as actionable as variant-level peaks for downstream biology. We conclude that full-GRM mixed-model GWAS, with the limited number of peaks it produces, is the appropriate default for livestock studies whose goal is SNP-level biological interpretation (e.g., candidate-gene prioritization, fine-mapping, or molecular-QTL colocalization).

### 5.5 Practical recommendations for livestock GWAS

The limited number of significant peaks identified by full-GRM mixed-model GWAS is well suited to focused functional validation: these are by construction the variants with the largest above-background effects, and by the unified-effect-size-floor argument they are also the variants most likely to be fine-mappable to a small number of candidate genes. In dairy cattle, a full-GRM sequence-variant GWAS of 30 traits in ~50,000 Holstein bulls identified a tractable number of peaks per trait, all of which could be examined for candidate genes through sparse-prior fine-mapping (Wang *et al*. 2025); this style of analysis exemplifies the recommended workflow. In our view, functional and validation resources in livestock genetics should preferentially target this narrower set of full-GRM peaks rather than the longer lists produced by alternative methods (such as LOCO and fastGWA), because the latter are dominated by block-level polygenic signals that cannot be narrowed to a small number of candidate genes.

The use of de-regressed breeding values (or de-regressed proofs; DRPs) as pseudo-phenotypes substantially amplifies the per-SNP detection power of livestock GWAS at fixed *N*, by raising the effective heritability to the cohort-average reliability and hence *S*(*N*) (Section 4.3 and Figure 3, bottom row). The *N*-equivalence formula 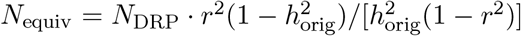 (Equation 11) converts this leverage into concrete numbers. For a recent dairy GWAS by Wang *et al*. (2025) — *N*_DRP_ = 50,309 Holstein bulls with cohort-average reliability *r*^2^ ≈ 0.8 on a milk-yield trait whose single-record heritability is 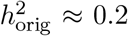 — the multiplier is 0.8 · 0.8*/*(0.2 · 0.2) = 16, so per-SNP detection power matches a single-record-phenotype GWAS of approximately 800,000 genotyped cows. This *N*-equivalence explains why moderate-cohort DRP-based GWAS in dairy populations recover meaningful sequence-variant peaks comparable to single-record studies of more than a million cows (e.g., Jiang *et al*. 2022), and it provides a quantitative basis for choosing between original-phenotype and DRP-based designs when both are feasible. Because de-regressed proofs can also be used as pseudo-phenotypes for genomic evaluation, this leverage benefits prediction as well as detection.

For practical analysis, we recommend full-GRM mixed-model implementations. GCTA --mlma (Yang *et al*. 2014) handles studies of tens of thousands of individuals and remains a convenient default at that scale. SLEMM (Cheng *et al*. 2023) scales to millions of genotyped individuals and sequence variants, and additionally accommodates breeding values as pseudo-phenotypes by modeling the per-individual reliability in the residual term, a common requirement in livestock applications. Where ssGBLUP has already been deployed for genomic evaluation, ssGWAS using APY (Leite *et al*. 2024) provides a convenient route to full-GRM-equivalent association testing without re-engineering the existing pipeline. All three approaches produce results consistent with the framework’s predictions in livestock settings.

### 5.6 The prediction–mapping trade-off under small *N*_*e*_

The small *N*_*e*_ that limits livestock GWAS makes livestock an easy setting for genomic prediction. Sections 2.2 and 2.7 traced both behaviors to the same sigmoid sum *S*(*N*): the per-SNP association NCP scales with *S*(*N*), while the in-sample GBLUP reliability is 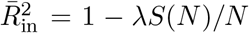 and the leave-one-out reliability lies below it. In livestock, where *M*_*e*_ is small and reference populations are large, *S*(*N*)*/N* ≈ *M*_*e*_*/N* is tiny, so both reliabilities are close to 1. Our framework quantifies both (the in-sample reliability exactly and the out-of-sample reliability by tight bounds) from realistic eigenvalue spectra rather than a heuristic, and recovers the classical Daetwyler reliability as a special case for *N* ≪ *M*_*e*_ (Section S10.7). The small *M*_*e*_ that makes prediction comparatively easy, however, caps the number of distinguishable above-background association signals in livestock. With genomic information of low effective dimensionality, a moderate-density chip panel of ~50,000 SNPs already contains largely all the LD information available. This is the theoretical basis for the empirical observation that including carefully selected variants from whole-genome sequences yields only marginal gains in prediction accuracy in dairy cattle (VanRaden *et al*. 2017) and related observations in other livestock species (Ros-Freixedes *et al*. 2022; Kamprasert *et al*. 2025). It is also why genomic selection has been a remarkable operational success across livestock breeding programs using only moderate-density chip data (e.g., García-Ruiz *et al*. 2016).

When a livestock reference population is already so large that the GBLUP reliability is approaching 1, external sources of biological prior information (e.g., molecular QTL mapping, functional annotation enrichment, and gene-set memberships) and additional sources of genetic variants (e.g., structural variants) are expected to deliver limited incremental gains over chip-based GBLUP/ssGBLUP, because the chip-based GRM can capture essentially all the LD-patterned genetic variance accessible within the *M*_*e*_-dimensional prediction space. This is the double-edged sword of small *N*_*e*_ in livestock genomics: it makes prediction comparatively easy, and mapping comparatively hard.

We have analyzed only GBLUP-type prediction, not variable-selection methods, whose advantage over GBLUP depends on the genetic architecture (Section 5.7). In livestock, GBLUP is nonetheless representative: it performs comparably even for traits with known large-effect QTL, the two types converge as the reference population grows, and weighting markers by minor allele frequency yields little gain (Cheng *et al*. 2023), so these conclusions apply to genomic prediction broadly in these populations.

### 5.7 Genetic architecture and the dimensionality of human polygenic prediction

Our reliability results assume an effectively infinitesimal genetic architecture (Section S10.1): heritability is spread across all *M* markers, so the dimensionality governing *S*(*N*) is the full *M*_*e*_ = 4*N*_*e*_*L*. For humans this is the pessimistic limit of prediction reliability, because *M*_*e*_ ≈ 1.4 × 10^6^ exceeds current biobank sample sizes, leaving *S*(*N*)*/N* of order *h*^2^ and the leave-one-out reliability far below 1 (Section S10.7). Yet many complex human traits are highly but not boundlessly polygenic: the variants carrying appreciable effect, though numerous, are far fewer than *M*_*e*_ (Yengo *et al*. 2022). Prioritizing or enriching for those variants reduces the effective prediction dimensionality entering *S*(*N*) from the trait-independent *M*_*e*_ toward a smaller trait-specific dimension, raising the attainable reliability at a given *N*; mapping is thus not only a parallel objective but a route to feasible prediction for the most polygenic human traits. Variable selection (Lloyd-Jones *et al*. 2019) or, more softly, the weighting of markers by functional priors (e.g., Márquez-Luna *et al*. 2021) accomplishes the same end, concentrating prior variance on likely-causal variants rather than spreading it uniformly across all markers. To sum up, human polygenic score prediction is less constrained than a literal substitution of *M*_*e*_ = 4*N*_*e*_*L* into our reliability formulas implies, because the dimensionality that matters can be the trait-specific number of relevant effects, not the trait-independent *M*_*e*_.

### 5.8 Scope and outlook

Three modeling choices in the present work simplify the genomic eigenvalue spectrum that underlies *M*_*e*_: we used the analytic formula *M*_*e*_ = 4*N*_*e*_*L* (which assumes a constant effective size rather than the bottlenecks, selection sweeps, and admixture of real livestock histories); our coalescent simulations used a simplified two-epoch demographic model (Section 3.2); and those simulations approximated chip ascertainment by a MAF ≥ 0.05 filter with uniform random thinning. Two facts make the conclusions robust to these choices. First, the value of *M*_*e*_ is insensitive to how it is computed: 4*N*_*e*_*L*, EIG98, and EIG99 agree to within about a factor of two (Supplementary Tables 6 and 7), well within an order of magnitude. Second, although the simulated spectrum is less precise than a real population’s, its essential feature is preserved: a small fraction of leading eigenvalues explains most of the spectral mass (low effective dimensionality), which is what the two-ceiling structure rests on. Prior effective-dimensionality and APY studies (Pocrnic *et al*. 2016; Pocrnic *et al*. 2016b) show that both the number of leading genomic eigenvalues explaining approximately 98% of total spectral mass and the APY core size at which genomic prediction performance stabilizes are close to 4*N*_*e*_*L*.

The framework’s predictions extend naturally to plant breeding. Programs based on outbred populations with large effective sizes should show human-like concordance among full-GRM, LOCO, and fastGWA results, whereas elite inbred-line breeding programs with strongly reduced *N*_*e*_ should show livestock-like divergence. A quantitative cross-species treatment of plant GWAS through this framework is a natural next step.

GWAS-related methods originally developed in human genetics (both mixed-model association methods and downstream methods that operate on GWAS summary statistics, such as LD-score regression, sparse-prior fine-mapping, and colocalization) are likely to continue being applied across species, but the species-by-method interaction we describe means their behavior in livestock may not be inferred from their behavior in humans (Jiang 2024; Wang *et al*. 2025b). Choosing the right method for each species and interpreting the results in light of the *M*_*e*_ ceiling are essential for accurate biological inference.

## Back Matter

### Data Availability

The three real-data genotype panels analyzed in this study are publicly available. The German Holstein 50K dataset accompanies Zhang *et al*. (2015). The Chinese Holstein 50K dataset accompanies Huang *et al*. (2019) and is deposited on Figshare (https://doi.org/10.6084/m9.figshare.5353498.v1). The Karacabey Merino 50K dataset accompanies Yaman *et al*. (2025) and is deposited on Figshare (https://doi.org/10.6084/m9.figshare.29184098.v1).

The complete analysis pipeline, including all scripts for coalescent genotype simulation, LD-matrix eigenvalue spectrum computation, per-SNP GRAMMAR-Gamma coefficient estimation, phenotype simulation and association testing, average-coefficient computation, and figure and table generation, is provided as a public code repository at https://github.com/jiang18/genomic-bounds. The repository also archives the LD-matrix eigenvalue arrays, the per-replicate score *χ*^2^ statistics, the per-population focal-SNP information, the average-coefficient tables for the full-GRM and LOCO analyses, and the per-SNP GRAMMAR-Gamma coefficient tables for the three real-data chip panels. The raw simulated genotype matrices, which run to several gigabytes per population, are not deposited; they can be regenerated from the documented msprime random seeds.

## Acknowledgments

The author thanks Christian Maltecca and Julong Wei for helpful comments on an earlier version of this manuscript. The author also thanks Peter M. Visscher for sharing a note on prediction accuracy that prompted the development of the genomic prediction sections.

## Funding

This work is supported by the Agriculture and Food Research Initiative (AFRI) Foundational and Applied Science Program, project award no. 2023-67015-39260, and the Research Capacity Fund (HATCH), project award no. 7008128, from the U.S. Department of Agriculture’s National Institute of Food and Agriculture.

## Conflicts of Interest

The author declares no conflicts of interest.

## Supplementary Figures

**Supplementary Figure 1.**
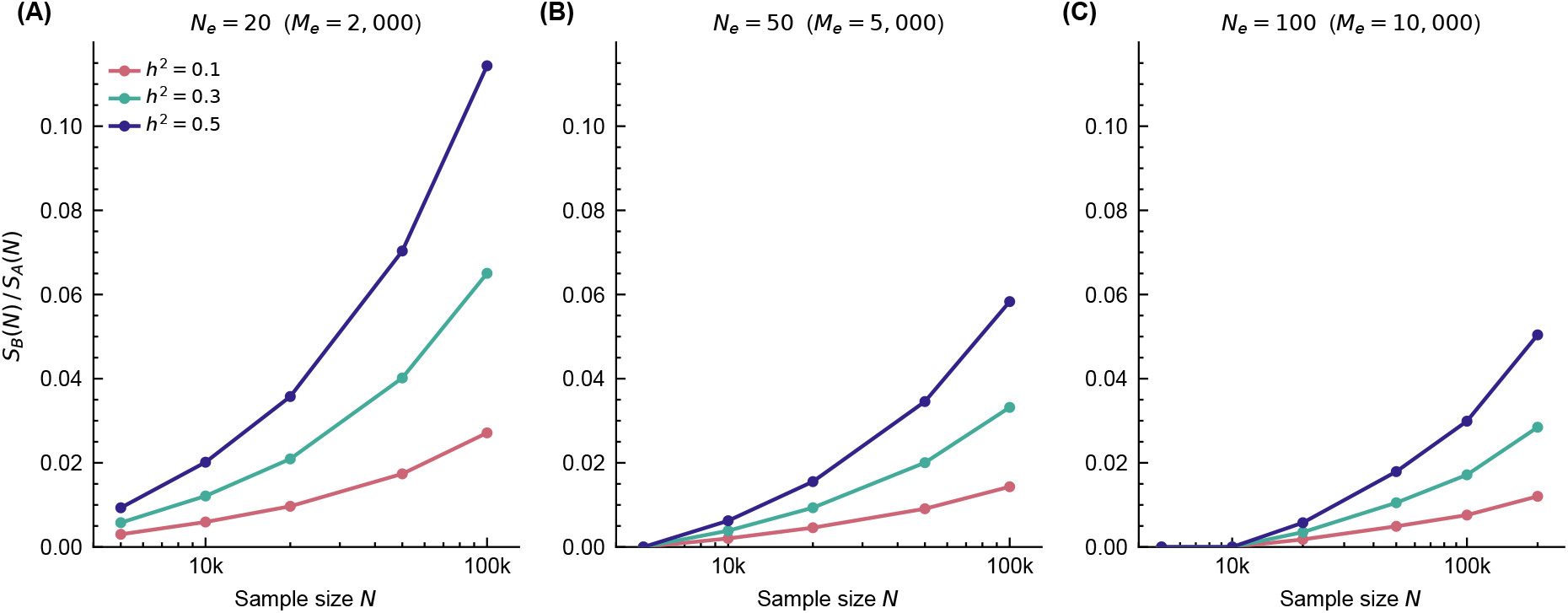
The Group B contribution to *S*(*N*) remains a small fraction of *S*_*A*_(*N*) across sample sizes. Ratio *S*_*B*_(*N*)*/S*_*A*_(*N*) as a function of sample size *N* (log scale), where *S*_*A*_(*N*) is the sigmoid sum (Equation 6) over the top *M*_*e*_ eigenmodes and *S*_*B*_(*N*) the sigmoid sum over the remaining *M* − *M*_*e*_ eigenmodes. Panels: (A) *N*_*e*_ = 20, (B) *N*_*e*_ = 50, and (C) *N*_*e*_ = 100, with *h*^2^ = 0.1 (rose), 0.3 (teal), and 0.5 (indigo) within each panel. The *y*-axis is shared across panels to make the between-*N*_*e*_ contrast direct. The ratio remains small (at or below 0.06 for *N*_*e*_ = 50 and *N*_*e*_ = 100 across the simulated grid; rising with both *h*^2^ and *N* to a peak of 0.11 for *N*_*e*_ = 20 at *h*^2^ = 0.5 and *N* = 100,000), confirming that the practical ceiling on *S*(*N*) is set by the top *M*_*e*_ eigenmodes (Section 4.3). *S*_*B*_*/S*_*A*_ is identically zero when *N* ≤ *M*_*e*_ because the sample LD matrix has rank min(*N, M*) and contains no Group-B eigenvalues in that regime, which affects the leftmost points of the *N*_*e*_ = 50 and *N*_*e*_ = 100 panels. Numerical values are reported in Supplementary Table 5.

**Supplementary Figure 2.**
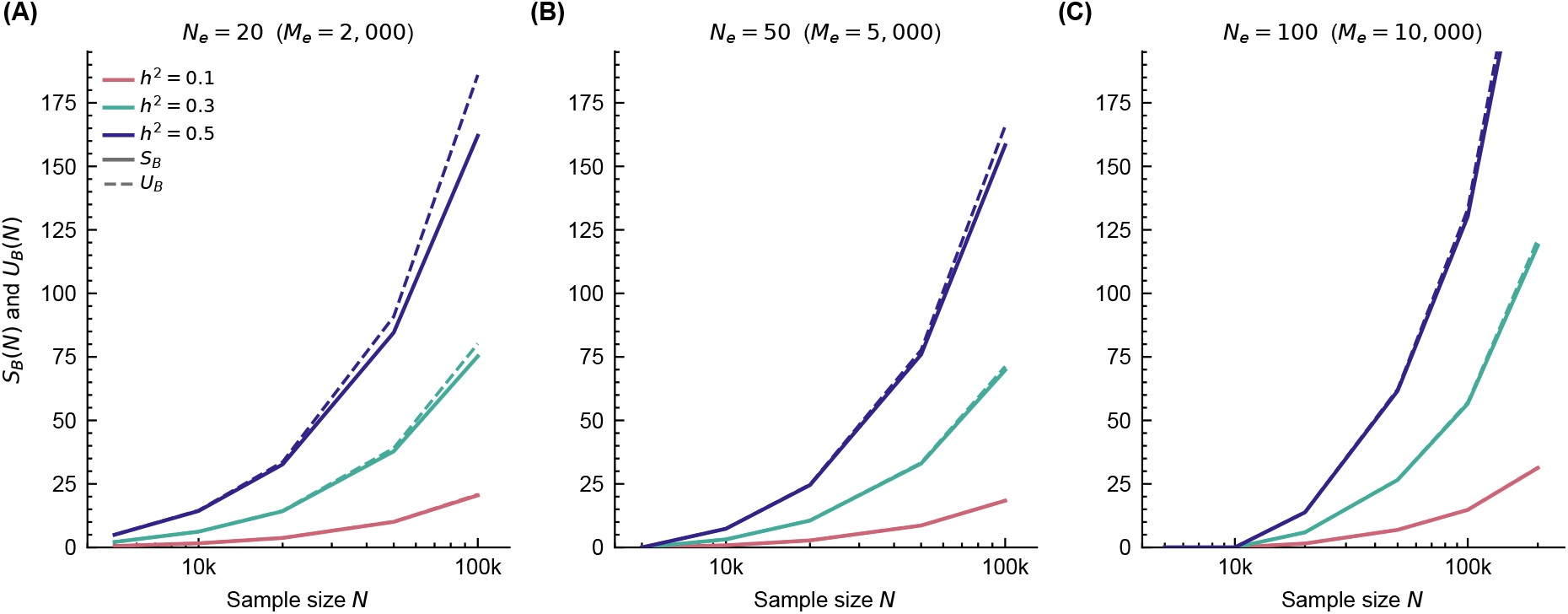
Jensen’s inequality is tight for Group B: *U*_*B*_(*N*) closely approximates *S*_*B*_(*N*). Exact Group B sigmoid sum *S*_*B*_(*N*) (solid lines) and its Jensen upper bound *U*_*B*_(*N*) = *N* (1−*ρ*)*M*_*B*_*/*[*N* (1−*ρ*)+*M*_*B*_*λ*] (dashed lines), where *λ* = (1−*h*^2^)*/h*^2^, *M*_*B*_ = *M* −*M*_*e*_, and *ρ* is the fraction of total eigenvalue mass held by the top *M*_*e*_ eigenvalues. Panels: (A) *N*_*e*_ = 20, (B) *N*_*e*_ = 50, and (C) *N*_*e*_ = 100, with *h*^2^ = 0.1 (rose), 0.3 (teal), and 0.5 (indigo). *S*_*B*_(*N*) and *U*_*B*_(*N*) track closely across all conditions, with a maximum relative gap (*U*_*B*_ − *S*_*B*_)*/U*_*B*_ of 13% (for *N*_*e*_ = 20 at *h*^2^ = 0.5 and *N* = 100,000), because the Group B eigenvalues are small, so the sigmoid function operates in a near-linear regime where Jensen’s inequality is essentially tight. This validates the use of *U*_*B*_(*N*) as a proxy for *S*_*B*_(*N*) in the analytic bound on the ratio *S*_*B*_*/S*_*A*_ derived in Supplementary Methods Section S7.5. As in Supplementary Figure 1, *S*_*B*_(*N*) and *U*_*B*_(*N*) are both identically zero in the regime *N* ≤ *M*_*e*_. Numerical values are reported in Supplementary Table 5.

## Supplementary Tables

**Supplementary Table 1. Per-SNP** *c*_*ℓ*_ **summary statistics in three livestock chip datasets (full-GRM and LOCO)**. Detailed statistics underlying Figure 1: for each (dataset, mode, *h*^2^) combination, the table reports *N* (sample size), *M* (SNP count), and the mean, SD, CV, minimum, 10th percentile, median, 90th percentile, and maximum of *c*_*ℓ*_. Datasets: Chinese Holstein (full-GRM + LOCO), Karacabey Merino (full-GRM + LOCO), and German Holstein (full-GRM only). Heritabilities: *h*^2^ = 0.1, 0.3, 0.5.

**Supplementary Table 2. Per-SNP non-centrality parameters underlying Figure 2**. Theoretical and empirical NCP for each of 360 focal-SNP scenarios (3 *N*_*e*_ × 3 *h*^2^ × 2 *β*^2^ × 4 *N* × 5 focal SNPs). For each row the table reports *N*_*e*_, *h*^2^, *β*^2^, *N*, focal-SNP identifier, chromosome, MAF (computed from all simulated individuals; one MAF per focal SNP, independent of subsample *N*), the eigenvalue-specific coefficient *c*_*ℓ*_, the theoretical NCP (from Equation 1), the empirical NCP (mean score *χ*^2^ minus 1 across replicates), and their ratio.

**Supplementary Table 3. Sigmoid sum** *S*(*N*) **values underlying Figure 3**. The table is organized into two blocks indicated by the fig3_row column. The **top-row block** (fig3_row = ‘top’; panels A–C of Figure 3) reports *S*(*N*) at heritability values *h*^2^ ∈ {0.1, 0.3, 0.5} over the full simulated sample-size grid for each *N*_*e*_. The **bottom-row block** (fig3_row = ‘bottom’; panels D–F of Figure 3) reports *S*(*N*) at the three fixed sample sizes *N* ∈ {10,000, 20,000, 50,000} over a dense heritability grid *h*^2^ ∈ {0.05, 0.10, 0.20, 0.30, 0.40, 0.50, 0.60, 0.70, 0.80, 0.90, 0.95} spanning the plotted range. Each row reports *N*_*e*_, *M, M*_*e*_ = 4*N*_*e*_*L, h*^2^, *N*, the scale parameter *Mλ* (with *λ* = (1 − *h*^2^)*/h*^2^), the full sigmoid sum *S*(*N*), its Group A component *S*_*A*_(*N*) (top *M*_*e*_ eigenmodes), its Group B component *S*_*B*_(*N*) (remaining eigenmodes), and the ratio *S*(*N*)*/M*_*e*_.

**Supplementary Table 4. Average GRAMMAR-Gamma coefficients** 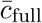 **and** 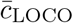 **underlying Figure 4**. For each (*N*_*e*_, *h*^2^, *N*) combination the table reports 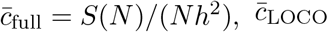, and the asymptotic LOCO upper bound 1*/*(1 − *h*^2^) (Equation 12).

**Supplementary Table 5. Group B properties underlying Supplementary Figures 1 and 2**. For each (*N*_*e*_, *h*^2^, *N*) combination the table reports *S*_*A*_(*N*), *S*_*B*_(*N*), and the ratio *S*_*B*_*/S*_*A*_ (plotted in Supplementary Figure 1), and *U*_*B*_(*N*), the gap *U*_*B*_ − *S*_*B*_, and the relative gap (*U*_*B*_ − *S*_*B*_)*/U*_*B*_ (the Jensen-tightness diagnostic plotted in Supplementary Figure 2). The near-zero relative gap validates the use of *U*_*B*_(*N*) as an analytic proxy for *S*_*B*_(*N*) (Supplementary Methods Section S7.5).

**Supplementary Table 6. Eigenspectrum parameters at the 98% spectral-mass** *M*_*e*_. Identical layout to Table 1 but with *M*_*e*_ defined operationally as the smallest number of top eigenvalues capturing 98% of total spectral mass. The added column `EIG98_over_4NeL` reports the ratio of this empirical *M*_*e*_ to the theoretical 4*N*_*e*_*L*.

**Supplementary Table 7. Eigenspectrum parameters at the 99% spectral-mass** *M*_*e*_. Identical to Supplementary Table 6 with a 99% mass cutoff. The added column <u>EIG99_over_4NeL</u> reports the ratio to the theoretical 4*N*_*e*_*L*.

**Supplementary Table 8. Empirical and predicted in-sample GBLUP reliability across the 36 simulated scenarios**. For each (*N*_*e*_, *h*^2^, *N*) combination of the phenotype simulations with *β*^2^ = 0.001 (100 replicates each), the table reports the empirical in-sample reliability (`R2_empirical_mean`, with its across-replicate standard deviation `R2_empirical_sd`), the predicted 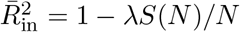 (`R2_theory`; Equation 17), and their ratio (`ratio_emp_over_theory`). These values underlie Figure 5.

## Notes

https://github.com/jiang18/genomic-bounds

### Competing Interest Statement

The authors have declared no competing interest.

### Summary of Updates

Genomic prediction reliability added as a third pillar (updated title and abstract): new Theory Section 2.7, Methods Section 3.4, Results Section 4.4, Supplementary Methods Section S10, Equations 17-19, Figure 5, and Supplementary Table 8 derive the in-sample and leave-one-out GBLUP reliability from the same quantity S(N); new Discussion Section 5.7 on human polygenic prediction. References added and minor clarifications throughout; association and fine-mapping results unchanged. Supplementary files updated.

## References

Franz Baumdicker, Gertjan Bisschop, Daniel Goldstein, Graham Gower, Aaron P Ragsdale, Georgia Tsambos, Sha Zhu, Bjarki Eldon, E Castedo Ellerman, Jared G Galloway, et al. Efficient ancestry and mutation simulation with msprime 1.0. Genetics, 220(3):iyab229, 2022.

Evan A Boyle, Yang I Li, and Jonathan K Pritchard. An expanded view of complex traits: from polygenic to omnigenic. Cell, 169:1177–1186, 2017. doi: 10.1016/j.cell.2017.05.038.

Christopher C Chang, Carson C Chow, Laurent C A M Tellier, Shashaank Vattikuti, Shaun M Purcell, and James J Lee. Second-generation PLINK: rising to the challenge of larger and richer datasets. GigaScience, 4:7, 2015. doi: 10.1186/s13742-015-0047-8.

Brian Charlesworth. Fundamental concepts in genetics: effective population size and patterns of molecular evolution and variation. Nature Reviews Genetics, 10:195–205, 2009. doi: 10.1038/nrg2526.

Jian Cheng, Christian Maltecca, Paul M VanRaden, Jeffrey R O’Connell, Li Ma, and Jicai Jiang. SLEMM: million-scale genomic predictions with window-based SNP weighting. Bioinformatics, 39(3):btad127, 2023. doi: 10.1093/bioinformatics/btad127.

Hans D Daetwyler, Beatriz Villanueva, and John A Woolliams. Accuracy of predicting the genetic risk of disease using a genome-wide approach. PLoS ONE, 3:e3395, 2008. doi: 10.1371/journal.pone.0003395.

Hans D Daetwyler, Ricardo Pong-Wong, Beatriz Villanueva, and John A Woolliams. The impact of genetic architecture on genome-wide evaluation methods. Genetics, 185(3):1021–1031, 2010. doi: 10.1534/genetics.110.116855.

George Davey Smith and Gibran Hemani. Mendelian randomization: genetic anchors for causal inference in epidemiological studies. Human Molecular Genetics, 23(R1):R89–R98, 2014. doi: 10.1093/hmg/ddu328.

Hilary K Finucane, Brendan Bulik-Sullivan, Alexander Gusev, Gosia Trynka, Yakir Reshef, Po-Ru Loh, Verneri Anttila, Han Xu, Chongzhi Zang, Kyle Farh, Stephan Ripke, Felix R Day, ReproGen Consortium, Schizophrenia Working Group of the Psychiatric Genomics Consortium, The RACI Consortium, Shaun Purcell, Eli Stahl, Sara Lindstrom, John R B Perry, Yukinori Okada, Soumya Raychaudhuri, Mark J Daly, Nick Patterson, Benjamin M Neale, and Alkes L Price. Partitioning heritability by functional annotation using genome-wide association summary statistics. Nature Genetics, 47:1228–1235, 2015. doi: 10.1038/ng.3404.

R A Fisher. The correlation between relatives on the supposition of Mendelian inheritance. Transactions of the Royal Society of Edinburgh, 52:399–433, 1918.

Adriana García-Ruiz, John B Cole, Paul M VanRaden, George R Wiggans, Felipe J Ruiz-López, and Curtis P Van Tassell. Changes in genetic selection differentials and generation intervals in U.S. Holstein dairy cattle as a result of genomic selection. Proceedings of the National Academy of Sciences, 113:E3995–E4004, 2016. doi: 10.1073/pnas.1519061113.

Dorian J Garrick, Jeremy F Taylor, and Rohan L Fernando. Deregressing estimated breeding values and weighting information for genomic regression analyses. Genetics Selection Evolution, 41:55, 2009. doi: 10.1186/1297-9686-41-55.

Claudia Giambartolomei, Damjan Vukcevic, Eric E Schadt, Lude Franke, Aroon D Hingorani, Chris Wallace, and Vincent Plagnol. Bayesian test for colocalisation between pairs of genetic association studies using summary statistics. PLoS Genetics, 10:e1004383, 2014. doi: 10.1371/journal.pgen.1004383.

Michael Goddard. Genomic selection: prediction of accuracy and maximisation of long term response. Genetica, 136:245–257, 2009. doi: 10.1007/s10709-008-9308-0.

Martien A M Groenen, Per Wahlberg, Mario Foglio, Hans H Cheng, Hendrik-Jan Megens, Richard P M A Crooijmans, Francois Besnier, Mark Lathrop, William M Muir, Gane Ka-Shu Wong, Ivo Gut, and Leif Andersson. A high-density SNP-based linkage map of the chicken genome reveals sequence features correlated with recombination rate. Genome Research, 19:510–519, 2009. doi: 10.1101/gr.086538.108.

Trevor Hastie, Robert Tibshirani, and Jerome Friedman. The Elements of Statistical Learning: Data Mining, Inference, and Prediction. Springer, New York, 2nd edition, 2009.

Hetian Huang, Jie Cao, Gang Guo, Xizhi Li, Yachun Wang, Ying Yu, Shengli Zhang, Qin Zhang, and Yi Zhang. Genome-wide association study identifies QTLs for displacement of abomasum in Chinese Holstein cattle. Journal of Animal Science, 97(3):1133–1142, 2019. doi: 10.1093/jas/skz031.

Sungbong Jang, Shogo Tsuruta, Natalia Galoro Leite, Ignacy Misztal, and Daniela Lourenco. Dimensionality of genomic information and its impact on genome-wide associations and variant selection for genomic prediction: a simulation study. Genetics Selection Evolution, 55:49, 2023. doi: 10.1186/s12711-023-00823-0.

Jicai Jiang. MPH: fast REML for large-scale genome partitioning of quantitative genetic variation. Bioinformatics, 40(5):btae298, 2024. doi: 10.1093/bioinformatics/btae298.

Jicai Jiang, Li Ma, Dzianis Prakapenka, Paul M VanRaden, John B Cole, and Yang Da. A large-scale genome-wide association study in U.S. Holstein cattle. Frontiers in Genetics, 10:412, 2019a. doi: 10.3389/fgene.2019.00412.

Jicai Jiang, Jian Cheng, Christian Maltecca, Li Ma, Paul M VanRaden, and Jeffrey R O’Connell. Mixed-model GWAS on milk production traits of 1.16M genotyped Holstein cattle. Journal of Dairy Science, 105(Suppl. 1):19 (abstr. 1047), 2022.

Longda Jiang, Zhili Zheng, Ting Qi, Kathryn E Kemper, Naomi R Wray, Peter M Visscher, and Jian Yang. A resource-efficient tool for mixed model association analysis of large-scale data. Nature Genetics, 51:1749–1755, 2019b. doi: 10.1038/s41588-019-0530-8.

Nantapong Kamprasert, Hassan Aliloo, Julius H J van der Werf, Christian J Duff, and Samuel A Clark. Effect of using preselected markers from imputed whole-genome sequence for genomic prediction in Angus cattle. Genetics Selection Evolution, 57:52, 2025. doi: 10.1186/s12711-025-00999-7.

Hyun Min Kang, Noah A Zaitlen, Claire M Wade, Andrew Kirby, David Heckerman, Mark J Daly, and Eleazar Eskin. Efficient control of population structure in model organism association mapping. Genetics, 178:1709–1723, 2008. doi: 10.1534/genetics.107.080101.

Fazhir Kayondo, Fernando Bussiman, Jorge Hidalgo, Ching-Yi Chen, Justin W Holl, Matias Bermann, and Daniela Lourenco. Single-step GWAS with APY and its application to a large pig population. In Plant and Animal Genome Conference (PAG 33), San Diego, California, USA, 2026.

Augustine Kong, Daniel F Gudbjartsson, Jesus Sainz, Gudrun M Jonsdottir, Sigurjon A Gudjonsson, Bjorgvin Richardsson, Sigrun Sigurdardottir, John Barnard, Bjorn Hallbeck, Gisli Masson, Adam Shlien, Stefan T Palsson, Michael L Frigge, Thorgeir E Thorgeirsson, Jeffrey R Gulcher, and Kari Stefansson. A high-resolution recombination map of the human genome. Nature Genetics, 31: 241–247, 2002. doi: 10.1038/ng917.

Natália Galoro Leite, Matias Bermann, Shogo Tsuruta, Ignacy Misztal, and Daniela Lourenco. Marker effect p-values for single-step GWAS with the algorithm for proven and young in large genotyped populations. Genetics Selection Evolution, 56:59, 2024. doi: 10.1186/s12711-024-00925-3.

Luke R Lloyd-Jones, Jian Zeng, Julia Sidorenko, Loïc Yengo, Gerhard Moser, Kathryn E Kemper, Huanwei Wang, Zhili Zheng, Reedik Mägi, Tõnu Esko, Andres Metspalu, Naomi R Wray, Michael E Goddard, Jian Yang, and Peter M Visscher. Improved polygenic prediction by Bayesian multiple regression on summary statistics. Nature Communications, 10(1):5086, 2019. doi: 10.1038/s41467-019-12653-0.

Po-Ru Loh, George Tucker, Brendan K Bulik-Sullivan, Bjarni J Vilhjálmsson, Hilary K Finucane, Rany M Salem, Daniel I Chasman, Paul M Ridker, Benjamin M Neale, Bonnie Berger, Nick Patterson, and Alkes L Price. Efficient Bayesian mixed-model analysis increases association power in large cohorts. Nature Genetics, 47:284–290, 2015. doi: 10.1038/ng.3190.

Emmanuel A Lozada-Soto, Francesco Tiezzi, Jicai Jiang, John B Cole, Paul M VanRaden, and Christian Maltecca. Genomic characterization of autozygosity and recent inbreeding trends in all major breeds of US dairy cattle. Journal of Dairy Science, 105:8956–8971, 2022. doi: 10.3168/jds.2022-22116.

Carla Márquez-Luna, Steven Gazal, Po-Ru Loh, Samuel S Kim, Nicholas Furlotte, Adam Auton, 23andMe Research Team, and Alkes L Price. Incorporating functional priors improves polygenic prediction accuracy in UK Biobank and 23andMe data sets. Nature Communications, 12(1):6052, 2021. doi: 10.1038/s41467-021-25171-9.

Ignacy Misztal, Andrés Legarra, and Ignacio Aguilar. Using recursion to compute the inverse of the genomic relationship matrix. Journal of Dairy Science, 97:3943–3952, 2014. doi: 10.3168/jds.2013-7752.

Ivan Pocrnic, Daniela A L Lourenco, Yutaka Masuda, Andrés Legarra, and Ignacy Misztal. The dimensionality of genomic information and its effect on genomic prediction. Genetics, 203:573–581, 2016a. doi: 10.1534/genetics.116.187013.

Ivan Pocrnic, Daniela A L Lourenco, Yutaka Masuda, and Ignacy Misztal. Dimensionality of genomic information and performance of the Algorithm for Proven and Young for different livestock species. Genetics Selection Evolution, 48(1):82, 2016b. doi: 10.1186/s12711-016-0261-6.

Ivan Pocrnic, Daniela Lourenco, and Ignacy Misztal. Single nucleotide polymorphism profile for quantitative trait nucleotide in populations with small effective size and its impact on mapping and genomic predictions. Genetics, 227(4):iyae103, 2024. doi: 10.1093/genetics/iyae103.

Saber Qanbari, Mathias Hansen, Steffen Weigend, Rudolf Preisinger, and Henner Simianer. Linkage disequilibrium reveals different demographic history in egg laying chickens. BMC Genetics, 11: 103, 2010. doi: 10.1186/1471-2156-11-103.

Roger Ros-Freixedes, Martin Johnsson, Andrew Whalen, Ching-Yi Chen, Bruno D Valente, William O Herring, Gregor Gorjanc, and John M Hickey. Genomic prediction with whole-genome sequence data in intensely selected pig lines. Genetics Selection Evolution, 54:65, 2022. doi: 10.1186/s12711-022-00756-0.

Daniel J Schaid, Wenan Chen, and Nicholas B Larson. From genome-wide associations to candidate causal variants by statistical fine-mapping. Nature Reviews Genetics, 19:491–504, 2018. doi: 10.1038/s41576-018-0016-z.

P Stam. The distribution of the fraction of the genome identical by descent in finite random mating populations. Genetics Research, 35:131–155, 1980.

J A Sved. Linkage disequilibrium and homozygosity of chromosome segments in finite populations. Theoretical Population Biology, 2:125–141, 1971. doi: 10.1016/0040-5809(71)90011-6.

Gulnara R Svishcheva, Tatiana I Axenovich, Nadezhda M Belonogova, Cornelia M van Duijn, and Yurii S Aulchenko. Rapid variance components–based method for whole-genome association analysis. Nature Genetics, 44:1166–1170, 2012. doi: 10.1038/ng.2410.

Flavie Tortereau, Bertrand Servin, Laurent Frantz, Hendrik-Jan Megens, Denis Milan, Gary Rohrer, Ralph Wiedmann, Jonathan Beever, Alan L Archibald, Lawrence B Schook, and Martien A M Groenen. A high density recombination map of the pig reveals a correlation between sex-specific recombination and GC content. BMC Genomics, 13:586, 2012. doi: 10.1186/1471-2164-13-586.

P M VanRaden. Efficient methods to compute genomic predictions. Journal of Dairy Science, 91: 4414–4423, 2008. doi: 10.3168/jds.2007-0980.

P M VanRaden, M E Tooker, J R O’Connell, J B Cole, and D M Bickhart. Selecting sequence variants to improve genomic predictions for dairy cattle. Genetics Selection Evolution, 49:32, 2017. doi: 10.1186/s12711-017-0307-4.

Junjian Wang, Yahui Gao, Sajjad Toghiani, John B Cole, Christian Maltecca, Li Ma, and Jicai Jiang. Genome-wide association study and fine-mapping using imputed sequences to prioritize candidate genes for 30 complex traits in 50,309 Holstein bulls. Journal of Dairy Science, 108(11): 12506–12518, 2025a. doi: 10.3168/jds.2025-27058.

Junjian Wang, Francesco Tiezzi, Yijian Huang, Garrett See, Clint Schwab, Julong Wei, Christian Maltecca, and Jicai Jiang. Fine-mapping methods for complex traits: essential adaptations for samples of related individuals. Briefings in Bioinformatics, 26(6):bbaf614, 2025b. doi: 10.1093/bib/bbaf614.

Zhiying Wang, Botong Shen, Jicai Jiang, Jinquan Li, and Li Ma. Effect of sex, age and genetics on crossover interference in cattle. Scientific Reports, 6:37698, 2016. doi: 10.1038/srep37698.

Yalçın Yaman, Ramazan Aymaz, Murat Keleş, Yiğit Emir Kişi, Ecem Hatipoğlu, Arzu Özdemir, and Elif Çetinkaya. Investigation of growth traits in Turkish Merino lambs using multi-locus GWAS approaches: Karacabey Merino. BMC Veterinary Research, 21(1):511, 2025. doi: 10.1186/s12917-025-04957-9.

Jian Yang, Noah A Zaitlen, Michael E Goddard, Peter M Visscher, and Alkes L Price. Advantages and pitfalls in the application of mixed-model association methods. Nature Genetics, 46:100–106, 2014. doi: 10.1038/ng.2876.

Loïc Yengo, Sailaja Vedantam, Eirini Marouli, et al. A saturated map of common genetic variants associated with human height. Nature, 610(7933):704–712, 2022. doi: 10.1038/s41586-022-05275-y.

Jianming Yu, Gael Pressoir, William H Briggs, Irie Vroh Bi, Masanori Yamasaki, John F Doebley, Michael D McMullen, Brandon S Gaut, Dahlia M Nielsen, James B Holland, Stephen Kresovich, and Edward S Buckler. A unified mixed-model method for association mapping that accounts for multiple levels of relatedness. Nature Genetics, 38:203–208, 2006. doi: 10.1038/ng1702.

Ricardo Zanella, Jane O Peixoto, Fernando F Cardoso, Leandro L Cardoso, Patrícia Biegelmeyer, Maurício E Cantão, Antonio Otaviano, Marcelo S Freitas, Alexandre R Caetano, and Mônica C Ledur. Genetic diversity analysis of two commercial breeds of pigs using genomic and pedigree data. Genetics Selection Evolution, 48:24, 2016. doi: 10.1186/s12711-016-0203-3.

Haonan Zeng, Wenjing Zhang, Qing Lin, Yahui Gao, Jinyan Teng, Zhiting Xu, Xiaodian Cai, Zhanming Zhong, Jun Wu, Yuqiang Liu, Shuqi Diao, Chen Wei, Wentao Gong, Xiangchun Pan, Zedong Li, Xiaoyu Huang, Xifan Chen, Jinshi Du, The PigGTEx Consortium, Fuping Zhao, Yunxiang Zhao, Maria Ballester, Daniel Crespo-Piazuelo, Marcel Amills, Alex Clop, Peter Karlskov-Mortensen, Merete Fredholm, Pinghua Li, Ruihua Huang, Guoqing Tang, Mingzhou Li, Xiaohong Liu, Yaosheng Chen, Qin Zhang, Jiaqi Li, Xiaolong Yuan, Xiangdong Ding, Lingzhao Fang, and Zhe Zhang. PigBiobank: a valuable resource for understanding genetic and biological mechanisms of diverse complex traits in pigs. Nucleic Acids Research, 52(D1):D980–D989, 2024. doi: 10.1093/nar/gkad1080.

Zhe Zhang, Malena Erbe, Jinlong He, Ulrike Ober, Ning Gao, Hao Zhang, Henner Simianer, and Jiaqi Li. Accuracy of whole-genome prediction using a genetic architecture-enhanced variance-covariance matrix. G3 Genes Genomes Genetics, 5:615–627, 2015. doi: 10.1534/g3.114.016261.

Wei Zhou, Jonas B Nielsen, Lars G Fritsche, Rounak Dey, Maiken E Gabrielsen, Brooke N Wolford, Jonathon LeFaive, Peter VandeHaar, Sarah A Gagliano, Aliya Gifford, Lisa A Bastarache, Wei-Qi Wei, Joshua C Denny, Maoxuan Lin, Kristian Hveem, Hyun Min Kang, Goncalo R Abecasis, Cristen J Willer, and Seunggeun Lee. Efficiently controlling for case-control imbalance and sample relatedness in large-scale genetic association studies. Nature Genetics, 50:1335–1341, 2018. doi: 10.1038/s41588-018-0184-y.

Xiang Zhou and Matthew Stephens. Genome-wide efficient mixed-model analysis for association studies. Nature Genetics, 44:821–824, 2012. doi: 10.1038/ng.2310.

