## Supplementary Methods for "Genomic Dimensionality Bounds Mixed-Model Association Power, Fine-Mapping Resolution, and Genomic Prediction Reliability"

Jicai Jiang*

June 25, 2026

Equations 1–19 are defined in the main text Theory section and are referenced here by number. Sections of this supplement carry an "S" prefix (e.g., Section S5.2).

---

### S1. Notation and Setup

#### S1.1. Generative Model

The generative model for the phenotype is:

$$\mathbf{y} = \mathbf{Z}\boldsymbol{\alpha} + \mathbf{e}$$

where

- $\mathbf{y}$ is the $N \times 1$ phenotype vector,
- $\mathbf{Z} = [\mathbf{z}_1, \ldots, \mathbf{z}_M]$ is the $N \times M$ matrix of standardized genotypes (with zero mean and unit variance) for all $M$ model SNPs (these need not be true causal variants),
- $\boldsymbol{\alpha} \sim \mathcal{N}(\mathbf{0}, \mathbf{W}\sigma_\alpha^2)$ is the vector of random SNP effects,
- $\mathbf{e} \sim \mathcal{N}(\mathbf{0}, \mathbf{I}\sigma_e^2)$ is the residual.

The generative model captures the phenotypic variation attributable to the model SNPs. The architecture matrix $\mathbf{W}$ is tied to the chosen SNP set: different SNP panels with their respective $\mathbf{W}$ can produce similar generative models, provided they adequately tag the same genomic segments. The focal SNP effect $\beta$ that we wish to test is implicit in $\boldsymbol{\alpha}$; it is introduced explicitly in the working model (Section S1.3).

The diagonal matrix $\mathbf{W} = \mathrm{diag}(W_{11}, \ldots, W_{MM})$ encodes the **effect-variance architecture** for the chosen set of model SNPs: all diagonal elements are close to 1 except a few that are significantly

*Department of Animal Science, North Carolina State University, Raleigh, NC 27607, USA.

larger than 1, corresponding to major-effect loci. The true $\mathbf{W}$ is unknown. When estimating variance components, we assume $\mathbf{W} = \mathbf{I}$; REML is robust to moderate violations of this assumption.

In practice, $M$ need not be very large. A moderate-density SNP panel ($M \sim$ 50,000) can effectively capture the genetic signal when $M \gg M_e$, because the $M_e$ independent genomic segments are fully tagged. Evidence from livestock genomic prediction supports this: prediction accuracy using ∼60,000 chip SNPs closely matches that from whole-genome sequence data (e.g., VanRaden *et al.* 2017), indicating that increasing marker density beyond a saturating panel adds negligible information.

We define $\tilde{\mathbf{W}} = \mathbf{W} - \mathbf{I}$, so that $\tilde{W}_{ss} = W_{ss} - 1$ measures the **excess effect** of SNP $s$ beyond the polygenic expectation. Under the polygenic null (all SNPs have equal variance), $\tilde{W}_{ss} = 0$. A major QTL has $\tilde{W}_{ss} > 0$.

Without loss of generality, we normalize the total phenotypic variance to unity: $M\sigma^2_\alpha + \sigma^2_e = 1$. The phenotypic variance explained (PVE) by all model SNPs is $h^2 = M\sigma^2_\alpha$, so $\sigma^2_e = 1 - h^2$. The PVE by SNP $s$ is $q = W_{ss}\sigma^2_\alpha = W_{ss}h^2/M$.

#### S1.2. SVD Decomposition

$\mathbf{Z}$ has thin SVD:

$$\mathbf{Z} = \mathbf{U}\boldsymbol{\Delta}\boldsymbol{\Phi}^T$$

where $\mathbf{U}$ is $N \times r$, $\boldsymbol{\Delta}$ is $r \times r$ diagonal with positive entries, $\boldsymbol{\Phi}$ is $M \times r$, and $r = \text{rank}(\mathbf{Z}) \leq \min(N, M)$. For standard SNP-chip panels after QC, exact linear dependencies are rare, so in practice $r$ is typically very close to $\min(N, M)$. This is consistent with the SNP saturation requirement ($M \gg M_e$ but with each SNP contributing non-redundant information at its genomic position).

#### S1.3. Working Model ($\mathbf{W} = \mathbf{I}$)

The working model tests the focal SNP $\mathbf{x}$ as a fixed effect against a polygenic background with equal effect variances ($\mathbf{W} = \mathbf{I}$):

$$\mathbf{y} = \mathbf{x}\beta + \mathbf{Z}\boldsymbol{\alpha} + \mathbf{e}$$

where $\mathbf{x}$ is the genotype vector at the focal SNP, $\beta$ is its fixed effect (the quantity being tested), $\boldsymbol{\alpha} \sim \mathcal{N}(\mathbf{0}, \mathbf{I}\sigma^2_\alpha)$, and $\mathbf{e} \sim \mathcal{N}(\mathbf{0}, \mathbf{I}\sigma^2_e)$. For full-GRM analysis, $\mathbf{Z}$ is the same as in the generative model (Section S1.1). Under leave-one-chromosome-out (LOCO) analysis, $\mathbf{Z} = \mathbf{Z}_{\text{LOCO}}$ excludes SNPs on the focal chromosome, and $\boldsymbol{\alpha} = \boldsymbol{\alpha}_{\text{LOCO}}$; the genetic effects of the focal chromosome are captured by $\mathbf{x}\beta$ and the residual. We use $\mathbf{Z}$ throughout for both cases, noting explicitly when $\mathbf{Z} = \mathbf{Z}_{\text{LOCO}}$.

The phenotypic variance-covariance matrix under the working model is:

$$\mathbf{V} = \frac{\mathbf{Z}\mathbf{Z}^T}{M} h^2 + \mathbf{I}(1 - h^2) = \mathbf{G}h^2 + \mathbf{I}(1 - h^2)$$

where $\mathbf{G} = \mathbf{Z}\mathbf{Z}^T/M$ is the GRM with eigenvalues $d_k$.

#### S1.4. LD Structure and Eigenvalues

For standardized genotypes, the sample LD correlation between SNPs $i$ and $j$ is:

$$r_{ij} = \frac{\mathbf{z}_i^T \mathbf{z}_j}{N}$$

The $M \times M$ LD matrix is $\mathbf{R} = \frac{1}{N}\mathbf{Z}^T\mathbf{Z}$, with eigenvalues $\ell_1 \geq \ell_2 \geq \cdots \geq \ell_M \geq 0$ satisfying $\sum_k \ell_k = \text{tr}(\mathbf{R}) = M$. Throughout, $r_{ij}$ denotes the sample LD correlation. The expected sample $r^2$ exceeds the population value by approximately $1/N$ (i.e., $E[r^2_{\text{sample}}] \approx r^2_{\text{population}} + 1/N$); for the sample sizes considered, this bias is negligible. The population expectation $E[r^2]$ under drift-recombination equilibrium (Section S1.5) is used only for quantitative predictions.

The GRM and LD matrix eigenvalues are related by $d_k = N\ell_k/M$.

#### S1.5. Key Population Genetic Parameters

For two loci separated by $d$ Morgans in a population with effective size $N_e$, the expected squared correlation at drift-recombination equilibrium is (Sved 1971):

$$E[r^2] \approx \frac{1}{1 + 4N_e d}$$

Therefore:

$$E[1 - r^2] \approx \frac{4N_e d}{1 + 4N_e d}$$

This is Equation 15. The characteristic segment length is $1/(4N_e)$ Morgans, yielding $M_e = 4N_eL$ independent segments across a genome of length $L$ Morgans (Stam 1980). Empirically, $M_e$ can be estimated as the number of LD matrix eigenvalues explaining ~98% of total eigenvalue mass, denoted $\text{EIG}_{98\%}$; eigenvalue analyses of livestock genotype data confirm $\text{EIG}_{98\%} \approx 4N_eL$ (Pocrnic *et al.* 2016).

#### S1.6. Notation Summary

| Symbol | Definition |
|---|---|
| $N$ | Sample size |
| $M$ | Number of SNPs in GRM |
| $r$ | Rank of $\mathbf{Z}$; $r = \min(N, M)$ in practice |
| $M_e$ | Effective number of independent segments $\approx 4N_eL$ |
| $N_e$ | Effective population size |
| $L$ | Genome length in Morgans |
| $h^2$ | PVE by all model SNPs $= M\sigma_\alpha^2$ |
| $\beta$ | Fixed effect of the focal SNP in the working model |
| $\mathbf{W}$ | Effect-variance architecture matrix; $W_{ss}$ is the weight of SNP $s$ |
| $\tilde{W}_{ss}$ | Excess effect weight $= W_{ss} - 1$ |
| $q$ | PVE of a focal SNP $= W_{ss}h^2/M$ |
| $c_l$ | Per-SNP GRAMMAR-Gamma coefficient $= \mathbf{z}_l^T\mathbf{V}^{-1}\mathbf{z}_l/N$ |
| $\bar{c}$ | Average GRAMMAR-Gamma coefficient over GRM SNPs |
| $c_{\text{LOCO}}$ | LOCO GRAMMAR-Gamma coefficient $= \mathbf{x}^T\hat{\mathbf{V}}_{\text{LOCO}}^{-1}\mathbf{x}/N$ |
| $S(N)$ | Sigmoid sum $= \sum_k N\ell_k/(N\ell_k + M\lambda)$ |
| $\lambda$ | Noise-to-signal ratio $= (1-h^2)/h^2$ |
| $d_k$ | GRM eigenvalues; $d_k = N\ell_k/M$ |
| $\ell_k$ | LD matrix eigenvalues; $\sum_k \ell_k = M$ |
| $r_{ls}^2$ | Squared sample LD correlation between SNPs $l$ and $s$ |
| $\mathbf{V}$ | Phenotypic covariance under the working model $= \mathbf{G}h^2 + \mathbf{I}(1-h^2)$ |
| $\rho$ | Fraction of eigenvalue mass in top $M_e$ eigenvalues |
| $h^2_{\text{LOCO}}$ | PVE by SNPs excluding the focal chromosome |
| $h^2_{\text{chr}}$ | PVE by SNPs on the focal chromosome; $h^2 = h^2_{\text{LOCO}} + h^2_{\text{chr}}$ |
| $M_{\text{LOCO}}$ | Number of SNPs excluding the focal chromosome |
| $M_{\text{chr}}$ | Number of SNPs on the focal chromosome |

---

### Part I: NCP of the Mixed-Model Association Test

This part derives the non-centrality parameter (NCP) of the score test under the mixed model, yielding the per-SNP formula (Equation 1) and its constant-coefficient ($\bar{c}$) approximation (Equation

5).

### S2. Test Statistic and Decomposition

**S2.1. GLS Estimate and Chi-Squared Statistic** The GLS estimate of $\beta$ and the score test statistic are:

$$\hat{\beta} = (\mathbf{x}^T\hat{\mathbf{V}}^{-1}\mathbf{x})^{-1}\,\mathbf{x}^T\hat{\mathbf{V}}^{-1}\mathbf{y}$$

$$\chi^2 = \frac{(\mathbf{x}^T\hat{\mathbf{V}}^{-1}\mathbf{y})^2}{\mathbf{x}^T\hat{\mathbf{V}}^{-1}\mathbf{x}}$$

where $\hat{\mathbf{V}} = \mathbf{G}\hat{h}^2 + \mathbf{I}(1-\hat{h}^2)$. Writing $\hat{\sigma}_e^2 = 1-\hat{h}^2$, factoring it out, and using the GRM eigendecomposition $\mathbf{G} = \mathbf{U}\,\mathrm{diag}(d_k)\,\mathbf{U}^T$:

$$\hat{\mathbf{V}} = \hat{\sigma}_e^2\left[\mathbf{U}\boldsymbol{\Lambda}\mathbf{U}^T + (\mathbf{I}-\mathbf{U}\mathbf{U}^T)\right]$$

where $\hat{\lambda} = (1-\hat{h}^2)/\hat{h}^2$ is the REML estimate of $\lambda$ and $\boldsymbol{\Lambda} = \mathrm{diag}(d_k/\hat{\lambda}+1)$ is the $r \times r$ diagonal matrix with $\Lambda_{kk} = d_k/\hat{\lambda}+1$. The matrix in brackets has eigenvalue $\Lambda_{kk}$ in $\mathrm{col}(\mathbf{U})$ and eigenvalue 1 in the orthogonal complement. We refer to $\mathrm{col}(\mathbf{U})$ as the **eigenspace** of the GRM throughout.

**S2.2. Decomposition of the Focal SNP** Decompose the focal SNP into within-eigenspace and orthogonal components:

$$\mathbf{x} = \mathbf{U}\mathbf{U}^T\mathbf{x} + (\mathbf{I}-\mathbf{U}\mathbf{U}^T)\mathbf{x} = \mathbf{U}\mathbf{a} + \mathbf{b}$$

where $\mathbf{a} = \mathbf{U}^T\mathbf{x}$ ($r \times 1$), $\mathbf{b} = (\mathbf{I}-\mathbf{U}\mathbf{U}^T)\mathbf{x}$ ($N \times 1$), $\mathbf{b}^T\mathbf{U} = \mathbf{0}$, and $\mathbf{x}^T\mathbf{x} = \mathbf{a}^T\mathbf{a} + \mathbf{b}^T\mathbf{b} = N$.

If $\mathbf{x}$ is column $l$ of $\mathbf{Z}$ (i.e., a GRM SNP), then $\mathbf{a} = \mathbf{U}^T\mathbf{z}_l$ and $\mathbf{b} = \mathbf{0}$ (since $\mathbf{z}_l \in \mathrm{col}(\mathbf{U})$).

**S2.3. Simplification of Quadratic Forms** From Section S2.1, $\hat{\mathbf{V}}/\hat{\sigma}_e^2 = \mathbf{U}\boldsymbol{\Lambda}\mathbf{U}^T + (\mathbf{I}-\mathbf{U}\mathbf{U}^T)$, with $\Lambda_{kk} = d_k/\hat{\lambda}+1$ in $\mathrm{col}(\mathbf{U})$ and eigenvalue 1 in the orthogonal complement. Its inverse is:

$$\hat{\mathbf{V}}^{-1} = \frac{1}{\hat{\sigma}_e^2}\left[\mathbf{U}\boldsymbol{\Lambda}^{-1}\mathbf{U}^T + (\mathbf{I}-\mathbf{U}\mathbf{U}^T)\right]$$

Substituting $\mathbf{x} = \mathbf{U}\mathbf{a} + \mathbf{b}$ and using $\mathbf{U}^T\mathbf{U} = \mathbf{I}_r$, $\mathbf{b}^T\mathbf{U} = \mathbf{0}$, and $(\mathbf{I}-\mathbf{U}\mathbf{U}^T)\mathbf{b} = \mathbf{b}$:

$$\hat{\sigma}_e^2\,\mathbf{x}^T\hat{\mathbf{V}}^{-1}\mathbf{y} = \mathbf{a}^T\boldsymbol{\Lambda}^{-1}\mathbf{U}^T\mathbf{y} + \mathbf{b}^T\mathbf{y}$$

$$\hat{\sigma}_e^2\,\mathbf{x}^T\hat{\mathbf{V}}^{-1}\mathbf{x} = \mathbf{a}^T\boldsymbol{\Lambda}^{-1}\mathbf{a} + \mathbf{b}^T\mathbf{b}$$

**S2.4. Weighted Decomposition of the Test Statistic** Define the two component statistics:

$$\chi^2_r = \frac{(\mathbf{b}^T\mathbf{y})^2}{\mathbf{b}^T\mathbf{b}\,\hat{\sigma}^2_e}, \qquad \chi^2_a = \frac{(\mathbf{a}^T\boldsymbol{\Lambda}^{-1}\mathbf{U}^T\mathbf{y})^2}{\mathbf{a}^T\boldsymbol{\Lambda}^{-1}\mathbf{a}\,\hat{\sigma}^2_e}$$

and the weights:

$$w_r = \frac{\mathbf{b}^T\mathbf{b}}{\mathbf{b}^T\mathbf{b} + \mathbf{a}^T\boldsymbol{\Lambda}^{-1}\mathbf{a}}, \qquad w_a = \frac{\mathbf{a}^T\boldsymbol{\Lambda}^{-1}\mathbf{a}}{\mathbf{b}^T\mathbf{b} + \mathbf{a}^T\boldsymbol{\Lambda}^{-1}\mathbf{a}}.$$

Substituting the results of Section S2.3 into the chi-squared statistic (Section S2.1):

$$\chi^2 = \frac{(\mathbf{a}^T\boldsymbol{\Lambda}^{-1}\mathbf{U}^T\mathbf{y} + \mathbf{b}^T\mathbf{y})^2}{(\mathbf{a}^T\boldsymbol{\Lambda}^{-1}\mathbf{a} + \mathbf{b}^T\mathbf{b})\,\hat{\sigma}^2_e} = \frac{\left(\sqrt{w_a}\,\frac{\mathbf{a}^T\boldsymbol{\Lambda}^{-1}\mathbf{U}^T\mathbf{y}}{\sqrt{\mathbf{a}^T\boldsymbol{\Lambda}^{-1}\mathbf{a}}} + \sqrt{w_r}\,\frac{\mathbf{b}^T\mathbf{y}}{\sqrt{\mathbf{b}^T\mathbf{b}}}\right)^2}{\hat{\sigma}^2_e}$$

Expanding the square, $E(\chi^2)$ contains $w_a\,E(\chi^2_a)$, $w_r\,E(\chi^2_r)$, and a cross-term proportional to $E(\mathbf{b}^T\mathbf{y} \cdot \mathbf{a}^T\boldsymbol{\Lambda}^{-1}\mathbf{U}^T\mathbf{y})$. The cross-term is zero. Under the generative model (Section S1.1), $E[\mathbf{y}] = \mathbf{0}$, so $E[\mathbf{y}\mathbf{y}^T] = \mathrm{Var}(\mathbf{y}) + E[\mathbf{y}]E[\mathbf{y}]^T = \mathrm{Var}(\mathbf{y})$. The cross-term therefore equals $\mathbf{b}^T\mathrm{Var}(\mathbf{y})\,\mathbf{U}\boldsymbol{\Lambda}^{-1}\mathbf{a}$.

Under the full-GRM generative model, $\mathrm{Var}(\mathbf{y}) = \mathbf{Z}\mathbf{W}\mathbf{Z}^T\sigma^2_\alpha + \mathbf{I}\sigma^2_e$. Under LOCO, $\mathbf{Z} = \mathbf{Z}_{\mathrm{LOCO}}$ (Section S1.3), and $\mathrm{Var}(\mathbf{y})$ additionally includes $\mathbf{Z}_{\mathrm{chr}}\mathbf{W}_{\mathrm{chr}}\mathbf{Z}^T_{\mathrm{chr}}h^2_{\mathrm{chr}}/M_{\mathrm{chr}}$ from the focal chromosome (Section S5), where $\mathbf{Z}_{\mathrm{chr}}$ is the focal chromosome genotype matrix, $\mathbf{W}_{\mathrm{chr}}$ its architecture matrix, $h^2_{\mathrm{chr}}$ the PVE, and $M_{\mathrm{chr}}$ the number of SNPs on that chromosome. Since $\mathbf{b} \perp \mathrm{col}(\mathbf{U}) = \mathrm{col}(\mathbf{Z})$, we have $\mathbf{b}^T\mathbf{Z} = \mathbf{0}$ and $\mathbf{b}^T\mathbf{U} = \mathbf{0}$, so the $\mathbf{Z}\mathbf{W}\mathbf{Z}^T$ term vanishes for any $\mathbf{W}$, and the $\mathbf{I}\sigma^2_e$ term gives $\sigma^2_e\,\mathbf{b}^T\mathbf{U}\boldsymbol{\Lambda}^{-1}\mathbf{a} = 0$. Under LOCO, $\mathbf{U} = \mathbf{U}_{\mathrm{LOCO}}$ (the eigenvectors of the LOCO GRM). In the ideal weak-cross-chromosome limit, $\mathbf{Z}^T_{\mathrm{chr}}\mathbf{U}_{\mathrm{LOCO}}$ is small, so the focal-chromosome contribution to this cross-term is also negligible. Thus the decomposition into $\chi^2_r$ and $\chi^2_a$ remains informative, with LOCO shifting weight toward the residual component.

Therefore:

$$E(\chi^2) = w_r\,E(\chi^2_r) + w_a\,E(\chi^2_a).$$

**Which component dominates depends on the SNPs used to construct the GRM.** If the GRM is constructed from a saturated set of whole-genome SNPs, then $\mathrm{col}(\mathbf{U})$ spans effectively the entire sample space, and the focal SNP $\mathbf{x}$ always lies in $\mathrm{col}(\mathbf{U})$: $\mathbf{b} = \mathbf{0}$, $w_r = 0$, and $w_a = 1$. In this case (full-GRM analysis), the test statistic is entirely determined by the within-eigenspace component $\chi^2_a$, which is subject to contamination-induced shrinkage.

Under LOCO analysis, the GRM excludes SNPs on the focal chromosome, so the focal SNP $\mathbf{x}$ typically projects much less strongly onto $\mathrm{col}(\mathbf{U}_{\mathrm{LOCO}})$ than under the full-GRM: $\|\mathbf{a}\|^2$ is reduced, $w_a$ is typically smaller, and $w_r$ correspondingly larger. The test statistic is therefore shifted toward the residual component $\chi^2_r$, which is free from contamination-induced shrinkage.

### S3. Expected Value of the Within-Eigenspace Component

**S3.1. Value of $E(\chi^2_a)$** For the full-GRM case where $w_a = 1$ (Section S2.4), $E(\chi^2) = E(\chi^2_a)$. Since $E[\mathbf{y}] = \mathbf{0}$ under the generative model:

$$E[(\mathbf{a}_l^T \boldsymbol{\Lambda}^{-1} \mathbf{U}^T \mathbf{y})^2] = \mathbf{a}_l^T \boldsymbol{\Lambda}^{-1} \mathbf{U}^T \mathrm{Var}(\mathbf{y})\, \mathbf{U} \boldsymbol{\Lambda}^{-1} \mathbf{a}_l$$

where $\mathbf{a}_l = \mathbf{U}^T \mathbf{x}$ is the focal SNP's projection onto the eigenspace. Substituting $\mathrm{Var}(\mathbf{y}) = \mathbf{Z}\mathbf{W}\mathbf{Z}^T \sigma^2_\alpha + \mathbf{I}\sigma^2_e$ and using $\mathbf{W} = \mathbf{I} + \tilde{\mathbf{W}}$:

$$\mathbf{U}^T \mathrm{Var}(\mathbf{y})\, \mathbf{U} = \mathbf{U}^T \mathbf{Z}\mathbf{W}\mathbf{Z}^T \mathbf{U} \sigma^2_\alpha + \mathbf{I}\sigma^2_e = (\boldsymbol{\Delta}^2 \sigma^2_\alpha + \mathbf{I}\sigma^2_e) + \sum_s \mathbf{a}_s \mathbf{a}_s^T \tilde{W}_{ss} \sigma^2_\alpha$$

where $\mathbf{a}_s = \mathbf{U}^T \mathbf{z}_s$ denotes the projection of GRM SNP $s$ onto the eigenspace, and we used $\mathbf{U}^T \mathbf{Z}\mathbf{Z}^T \mathbf{U} = \boldsymbol{\Delta}^2$ (from the SVD) and $\mathbf{U}^T \mathbf{Z}\tilde{\mathbf{W}}\mathbf{Z}^T \mathbf{U} = \sum_s \mathbf{a}_s \mathbf{a}_s^T \tilde{W}_{ss}$. For the first term, using $\Delta^2_k = M d_k$ and $M\sigma^2_\alpha/\sigma^2_e = 1/\lambda$: $\boldsymbol{\Delta}^2 \sigma^2_\alpha + \mathbf{I}\sigma^2_e = \sigma^2_e\, \mathrm{diag}(d_k/\lambda + 1)$. Since $\Lambda_{kk} = d_k/\hat{\lambda} + 1$ (Section S2.1), this can be written as $\sigma^2_e\Big(\boldsymbol{\Lambda} + \mathrm{diag}\Big(d_k(1/\lambda - 1/\hat{\lambda})\Big)\Big)$. Let $\rho_e = \sigma^2_e/\hat{\sigma}^2_e$. Dividing by the denominator $\mathbf{a}_l^T \boldsymbol{\Lambda}^{-1} \mathbf{a}_l\, \hat{\sigma}^2_e$:

$$E(\chi^2_a) = \rho_e + \frac{\mathbf{a}_l^T \boldsymbol{\Lambda}^{-1} \mathbf{D} \boldsymbol{\Lambda}^{-1} \mathbf{a}_l}{\mathbf{a}_l^T \boldsymbol{\Lambda}^{-1} \mathbf{a}_l} \left(\frac{1}{\lambda} - \frac{1}{\hat{\lambda}}\right) \rho_e + \frac{\sum_s (\mathbf{a}_l^T \boldsymbol{\Lambda}^{-1} \mathbf{a}_s)^2 \tilde{W}_{ss} \sigma^2_\alpha}{\mathbf{a}_l^T \boldsymbol{\Lambda}^{-1} \mathbf{a}_l\, \hat{\sigma}^2_e}$$

where $\mathbf{D} = \mathrm{diag}(d_1, \ldots, d_r)$ is the diagonal matrix of GRM eigenvalues.

**S3.2. Simplification Under Accurate Variance Component Estimation** When REML accurately estimates both variance components ($\hat{h}^2 \approx h^2$, equivalently $\hat{\lambda} \approx \lambda$), $\rho_e = 1$ and $1/\lambda - 1/\hat{\lambda} = 0$. The middle term in Section S3.1 vanishes, and the expected chi-squared reduces to:

$$E(\chi^2_a) = 1 + \frac{\sum_s (\mathbf{a}_l^T \boldsymbol{\Lambda}^{-1} \mathbf{a}_s)^2 \tilde{W}_{ss} \sigma^2_\alpha}{\mathbf{a}_l^T \boldsymbol{\Lambda}^{-1} \mathbf{a}_l\, \hat{\sigma}^2_e}$$

The NCP (the excess of $E(\chi^2)$ above 1) is therefore:

$$\mathrm{NCP}_l = \frac{\sum_s (\mathbf{a}_l^T \boldsymbol{\Lambda}^{-1} \mathbf{a}_s)^2 \tilde{W}_{ss} \sigma^2_\alpha}{\mathbf{a}_l^T \boldsymbol{\Lambda}^{-1} \mathbf{a}_l\, \hat{\sigma}^2_e}$$

To connect this to observable LD correlations, note that for GRM SNPs $\mathbf{z}_l \in \mathrm{col}(\mathbf{U})$, so $\mathbf{z}_l = \mathbf{U}\mathbf{U}^T \mathbf{z}_l = \mathbf{U}\mathbf{a}_l$. From Section S2.3, $\hat{\mathbf{V}}^{-1} = (1/\hat{\sigma}^2_e)[\mathbf{U}\boldsymbol{\Lambda}^{-1}\mathbf{U}^T + (\mathbf{I} - \mathbf{U}\mathbf{U}^T)]$. Therefore:

$$\hat{\sigma}^2_e\, \mathbf{z}_l^T \hat{\mathbf{V}}^{-1} \mathbf{z}_s = \mathbf{z}_l^T [\mathbf{U}\boldsymbol{\Lambda}^{-1}\mathbf{U}^T + (\mathbf{I} - \mathbf{U}\mathbf{U}^T)] \mathbf{z}_s = \mathbf{a}_l^T \boldsymbol{\Lambda}^{-1} \mathbf{a}_s + \mathbf{z}_l^T (\mathbf{I} - \mathbf{U}\mathbf{U}^T) \mathbf{z}_s = \mathbf{a}_l^T \boldsymbol{\Lambda}^{-1} \mathbf{a}_s$$

since $(\mathbf{I} - \mathbf{U}\mathbf{U}^T)\mathbf{z}_s = \mathbf{0}$ for GRM SNPs. The denominator gives $\mathbf{a}_l^T \boldsymbol{\Lambda}^{-1} \mathbf{a}_l = \hat{\sigma}^2_e\, c_l N$, where

$c_l = \mathbf{z}_l^T \hat{\mathbf{V}}^{-1} \mathbf{z}_l / N$ is the per-SNP GRAMMAR-Gamma coefficient (Section S4.2). Substituting:

$$\text{NCP}_l = \frac{\sigma_\alpha^2}{c_l N} \sum_s (\mathbf{z}_l^T \hat{\mathbf{V}}^{-1} \mathbf{z}_s)^2 \tilde{W}_{ss}$$

We extend the GRAMMAR-Gamma approximation to cross-terms: for SNPs $s$ in strong LD with the focal SNP $l$, the cross-term coefficient is approximately the same as the diagonal coefficient, giving $\mathbf{z}_l^T \hat{\mathbf{V}}^{-1} \mathbf{z}_s \approx c_l \, \mathbf{z}_l^T \mathbf{z}_s = c_l \, N \, r_{ls}$ (Section S4.3). These are precisely the SNPs that dominate the NCP sum through $r_{ls}^2$. Substituting:

$$\text{NCP}_l \approx \frac{c_l \, N \, h^2}{M} \sum_s r_{ls}^2 \tilde{W}_{ss}$$

This is the per-SNP NCP with SNP-specific $c_l$ (Equation 1). The GRAMMAR-Gamma $\bar{c}$ approximation further replaces the SNP-specific coefficient $c_l$ with the average $\bar{c}$ (a constant across all SNPs; Section S4.3), yielding the constant-coefficient form (Section S4.4).

### S4. Per-SNP NCP and the GRAMMAR-Gamma Approximation

**S4.1. Per-SNP NCP** The score chi-squared statistic for focal SNP $l$ (Section S2.1) has, under the generative model with accurately estimated variance components (Section S3.2), the non-centrality parameter:

$$\boxed{\text{NCP}_l \approx \frac{c_l \cdot N \cdot h^2}{M} \cdot \sum_s r_{ls}^2 \tilde{W}_{ss}}$$

where $c_l$ is the **per-SNP GRAMMAR-Gamma coefficient**:

$$c_l = \frac{\mathbf{z}_l^T \mathbf{V}^{-1} \mathbf{z}_l}{N}$$

This is Equation 1; the coefficient $c_l$ (Equation 2) is defined immediately above. The NCP retains the SNP-specific coefficient $c_l$, allowing it to vary across SNPs according to their projections onto the GRM eigenspace.

Here and below, $h^2$ denotes the working-model value used to construct $\mathbf{V}$; likewise, $h^2_{\text{LOCO}}$ in Section S5 denotes the working-model value used in the LOCO covariance. We omit hats in the coefficient formulas for notational simplicity.

**S4.2. Eigenvalue Expansion of $c_l$** Using the eigendecomposition $\mathbf{V}^{-1} = \mathbf{U} \, \text{diag}\left(\frac{1}{d_k h^2 + 1 - h^2}\right) \mathbf{U}^T + \frac{1}{1-h^2}(\mathbf{I} - \mathbf{U}\mathbf{U}^T)$ (Section S2.3), the per-SNP coefficient for a GRM SNP $\mathbf{z}_l$ is:

$$c_l = \frac{1}{N} \sum_k \frac{a_{lk}^2}{d_k h^2 + 1 - h^2}$$

where $a_{lk} = \mathbf{u}_k^T \mathbf{z}_l$ is the projection of the focal SNP onto the $k$-th GRM eigenvector. The $(\mathbf{I} - \mathbf{U}\mathbf{U}^T)$ term does not contribute because GRM SNPs lie entirely within $\mathrm{col}(\mathbf{U})$, so $\mathbf{b}_l = (\mathbf{I} - \mathbf{U}\mathbf{U}^T)\mathbf{z}_l = \mathbf{0}$.

This is Equation 3. The eigenvalue expansion reveals that $c_l$ depends on how the focal SNP's genotype vector distributes across the GRM eigenspace. Since the denominator $d_k h^2 + 1 - h^2$ is largest for top eigenvalues, projections onto top eigenvectors are more heavily shrunk. A SNP whose projection concentrates on large-eigenvalue modes (e.g., one in strong LD with many other GRM SNPs) will therefore have a **smaller** $c_l$: its effect is more thoroughly absorbed by the GRM random effects. Conversely, a SNP whose projections are concentrated on small eigenvalues will have a larger $c_l$, approaching $1/(1 - h^2)$.

**S4.3. The Average Coefficient $\bar{c}$ and the Global Approximation** Averaging $c_l$ over all $M$ GRM SNPs and using $\frac{1}{M}\sum_l a_{lk}^2 = d_k$ (since $\sum_l a_{lk}^2 = \sum_l (\mathbf{u}_k^T \mathbf{z}_l)^2 = \mathbf{u}_k^T \mathbf{Z}\mathbf{Z}^T \mathbf{u}_k = M d_k$):

$$\boxed{\bar{c} = \frac{1}{M}\sum_{l=1}^{M} c_l = \frac{1}{N}\sum_{k=1}^{r} \frac{d_k}{d_k h^2 + 1 - h^2}}$$

where $r = \mathrm{rank}(\mathbf{Z})$. For standard chip panels after QC, $r$ is typically very close to $\min(N, M)$. This is Equation 4. The expression depends on the **spectral structure of the GRM**, not on $M$ directly, ensuring invariance to SNP duplication in the GRM.

The GRAMMAR-Gamma method approximates $c_l \approx \bar{c}$ for all SNPs, replacing the per-SNP quadratic form with a global scalar. This approximation is accurate when individual SNPs have similar projections onto the GRM eigenspace, which holds for well-designed genotyping panels where SNPs are approximately evenly spaced and moderately correlated. For such panels, the coefficient of variation (CV) of $c_l$ across all SNPs is smaller than 0.3 empirically. In genotype panels containing many redundant markers, $c_l$ can deviate substantially from $\bar{c}$ (CV $> 0.5$).

**S4.4. NCP in Terms of LD Matrix Eigenvalues ($\ell_k$ form)** Substituting $d_k = N\ell_k/M$ into the $\bar{c}$ equation of Section S4.3:

$$\bar{c} = \frac{1}{N}\sum_{k=1}^{r} \frac{d_k}{d_k h^2 + 1 - h^2} = \frac{1}{N}\sum_{k=1}^{r} \frac{N\ell_k/M}{(N\ell_k/M)h^2 + 1 - h^2} = \frac{1}{Nh^2}\sum_{k=1}^{r} \frac{N\ell_k}{N\ell_k + M\lambda}$$

The NCP under the $\bar{c}$ approximation becomes:

$$\boxed{\mathrm{NCP} \approx \frac{S(N)}{M}\sum_s r_{ls}^2 \tilde{W}_{ss}}$$

where $S(N) = \sum_{k=1}^{r} \frac{N\ell_k}{N\ell_k + M\lambda} = \bar{c}\, N\, h^2$ is the **sigmoid sum** (Equation 6). This form makes the sample size dependence explicit: each eigenmode contributes a sigmoid in $\log N$ that rises from 0 to 1, with half-saturation at $N = M\lambda/\ell_k$. The equivalent $d_k$ form, $S(N) = \sum_k d_k/(d_k + \lambda)$, hides $N$

inside $d_k = N\ell_k/M$. The properties of $S(N)$ are established in Part II.

**NCP in the ideal tagging case.** For a single focal signal whose effect is fully captured by the generative model and local LD tagging (the focal SNP need not be the literal causal variant; the signal's effect can be carried by the focal SNP directly or distributed across nearby GRM SNPs in tight LD), $\sum_s r_{ls}^2 \tilde{W}_{ss} \approx qM/h^2 - 1$ (where $q = W_{ss}h^2/M$ is the focal SNP's PVE; Section S1.1). Substituting into the boxed NCP equation gives the compact form:

$$\boxed{\text{NCP} \approx S(N)\left(\frac{q}{h^2} - \frac{1}{M}\right)}$$

This is Equation 7. At chip scale ($M \sim 50{,}000$) the $1/M$ term is negligible compared to $q/h^2$ in practice, so NCP $\approx S(N) \cdot q/h^2$.

**S4.5. Invariance to SNP Duplication** If all SNPs in the GRM are duplicated, by the Kronecker product theorem the $2M \times 2M$ LD matrix has eigenvalues $2\ell_k$ (for $k = 1, \ldots, M$) and $M$ additional zeros. Under duplication: $M \to 2M$, $M\lambda \to 2M\lambda$, and $\sum_s r_{ls}^2 \tilde{W}_{ss}$ doubles (each term counted twice). The NCP:

$$\frac{1}{2M}\sum_k \frac{N \cdot 2\ell_k}{N \cdot 2\ell_k + 2M\lambda} \cdot 2\sum_s r_{ls}^2 \tilde{W}_{ss} = \frac{1}{M}\sum_k \frac{N\ell_k}{N\ell_k + M\lambda} \cdot \sum_s r_{ls}^2 \tilde{W}_{ss}$$

is exactly invariant, confirming that the NCP depends on the spectral structure of the GRM, not on the raw marker count.

### S5. Leave-One-Chromosome-Out (LOCO)

**S5.1. LOCO Working and Generative Models** Under LOCO, $\mathbf{Z} = \mathbf{Z}_{\text{LOCO}}$ excludes the focal chromosome (Section S1.3). The working model is:

$$\mathbf{y} = \mathbf{x}\beta + \mathbf{Z}_{\text{LOCO}}\boldsymbol{\alpha}_{\text{LOCO}} + \mathbf{e}$$

with $\hat{\mathbf{V}}_{\text{LOCO}} = \mathbf{G}_{\text{LOCO}}\hat{h}^2_{\text{LOCO}} + \mathbf{I}(1 - \hat{h}^2_{\text{LOCO}})$. Since the focal chromosome's genetic variance $h^2_{\text{chr}}$ is not modeled, it is absorbed into the residual estimate: $1 - \hat{h}^2_{\text{LOCO}} \approx 1 - h^2_{\text{LOCO}} = (1 - h^2) + h^2_{\text{chr}}$.

Under the generative model, the true phenotypic variance includes the focal chromosome:

$$\text{Var}(\mathbf{y}) = \mathbf{Z}_{\text{chr}}\mathbf{W}_{\text{chr}}\mathbf{Z}_{\text{chr}}^T \frac{h^2_{\text{chr}}}{M_{\text{chr}}} + \mathbf{V}_{\text{LOCO}}$$

where $\mathbf{Z}_{\text{chr}}$ is the $N \times M_{\text{chr}}$ genotype matrix of the focal chromosome, $\mathbf{W}_{\text{chr}}$ its architecture matrix, and $\mathbf{V}_{\text{LOCO}} = \mathbf{G}_{\text{LOCO}}h^2_{\text{LOCO}} + \mathbf{I}(1 - h^2)$ with $h^2 = h^2_{\text{chr}} + h^2_{\text{LOCO}}$.

**S5.2. LOCO Expected Chi-Squared** The test statistic is:

$$\chi^2 = \frac{(\mathbf{x}^T\hat{\mathbf{V}}_{\text{LOCO}}^{-1}\mathbf{y})^2}{\mathbf{x}^T\hat{\mathbf{V}}_{\text{LOCO}}^{-1}\mathbf{x}}$$

By the same eigendecomposition as in Section S2.3 (with $\mathbf{U} = \mathbf{U}_{\text{LOCO}}$), the inverse is:

$$\hat{\mathbf{V}}_{\text{LOCO}}^{-1} = \mathbf{U}_{\text{LOCO}}\,\text{diag}\left(\frac{1}{d_k\hat{h}^2_{\text{LOCO}} + 1 - \hat{h}^2_{\text{LOCO}}}\right)\mathbf{U}_{\text{LOCO}}^T + \frac{1}{1-\hat{h}^2_{\text{LOCO}}}(\mathbf{I} - \mathbf{U}_{\text{LOCO}}\mathbf{U}_{\text{LOCO}}^T)$$

**The LOCO coefficient.** Decomposing the focal SNP as in Section S2.2, $\mathbf{x} = \mathbf{U}_{\text{LOCO}}\mathbf{a} + \mathbf{b}$ with $\|\mathbf{a}\|^2 + \|\mathbf{b}\|^2 = N$:

$$c_{\text{LOCO}} = \frac{\mathbf{x}^T\hat{\mathbf{V}}_{\text{LOCO}}^{-1}\mathbf{x}}{N} = \frac{1}{N}\left[\sum_k \frac{a_k^2}{d_k h^2_{\text{LOCO}} + 1 - h^2_{\text{LOCO}}} + \frac{\|\mathbf{b}\|^2}{1 - h^2_{\text{LOCO}}}\right]$$

The LOCO coefficient $c_{\text{LOCO}}$ and its upper bound $1/(1-h^2_{\text{LOCO}})$ constitute Equation 12. The value of $c_{\text{LOCO}}$ depends on $\|\mathbf{a}\|^2$ (variance captured by the LOCO eigenspace) versus $\|\mathbf{b}\|^2 = N - \|\mathbf{a}\|^2$ (variance in the orthogonal complement). The orthogonal complement contributes $1/(1-h^2_{\text{LOCO}})$ per unit of $\|\mathbf{b}\|^2$, while the eigenspace contributes less per unit of $\|\mathbf{a}\|^2$ because of shrinkage ($1/(d_k h^2_{\text{LOCO}} + 1 - h^2_{\text{LOCO}}) < 1/(1-h^2_{\text{LOCO}})$ for $d_k > 0$). Therefore, transferring variance from $\|\mathbf{b}\|^2$ to $\|\mathbf{a}\|^2$ always **reduces** $c_{\text{LOCO}}$.

Since $\mathbf{x}$ is on the left-out chromosome, it has no physical linkage with LOCO SNPs. However, $\|\mathbf{a}\|^2 > 0$ due to two sources:

1. **Random projections.** For unrelated samples, the standardized focal-chromosome SNP satisfies $E[\mathbf{x}\mathbf{x}^T] \approx \mathbf{I}$ (each entry has zero mean and unit variance by construction). Since $\mathbf{u}_k$ is a unit eigenvector of $\mathbf{G}_{\text{LOCO}}$, built from a disjoint set of chromosomes and therefore approximately independent of $\mathbf{x}$, we have $E[a_k^2] = E[(\mathbf{u}_k^T\mathbf{x})^2] = \mathbf{u}_k^T E[\mathbf{x}\mathbf{x}^T]\mathbf{u}_k \approx \|\mathbf{u}_k\|^2 = 1$. This gives $\|\mathbf{a}\|^2_{\text{noise}} \approx r$, where $r = \text{rank}(\mathbf{G}_{\text{LOCO}})$. Crucially, the expected projection is 1, not 0: even without physical linkage, finite-sample correlations produce nonzero projections that scale with the rank of the LOCO eigenspace.

2. **Pedigree structure.** In livestock populations, individuals share pedigree relationships that create correlations between genotypes at unlinked loci. The top eigenvectors of $\mathbf{G}_{\text{LOCO}}$ (large $d_k$) capture these relationships, and the focal SNP shares the same pedigree, so $E[a_k^2]$ can be substantially larger than 1 on these modes. This further increases $\|\mathbf{a}\|^2$ beyond the random-projection baseline, reducing $c_{\text{LOCO}}$ because the additional variance lands on heavily-shrunk eigenvectors.

**Limiting cases.** The formula interpolates:

- $N \leq M_{\text{LOCO}}$**:** In this regime, $r \approx N$, so the LOCO eigenspace is effectively full rank, $\|\mathbf{b}\|^2 \approx 0$, and $\|\mathbf{a}\|^2 \approx N$. Then $c_{\text{LOCO}} = (1/N)\sum_k a_k^2/(d_k h^2_{\text{LOCO}} + 1 - h^2_{\text{LOCO}})$, with the constraint $\sum_k a_k^2 = N$ and $\bar{d} = \sum_k d_k/N = 1$. If the projections are approximately isotropic, so that $a_k^2 \approx 1$, then $c_{\text{LOCO}} = (1/N)\sum_k 1/(d_k h^2_{\text{LOCO}} + 1 - h^2_{\text{LOCO}}) \geq 1$ by Jensen's inequality (convexity of $1/(dh^2_{\text{LOCO}} + 1 - h^2_{\text{LOCO}})$, with $\bar{d} = 1$). Thus, without strong pedigree structure, $c_{\text{LOCO}}$ is expected to be close to 1, typically slightly above 1 when the $d_k$ are dispersed. If some projections are concentrated on leading large-$d_k$ modes, then $c_{\text{LOCO}}$ is reduced because those modes are shrunk most strongly by $\hat{\mathbf{V}}^{-1}_{\text{LOCO}}$.

- $N \gg M_{\text{LOCO}}$**, weak pedigree:** In this regime, $r \approx M_{\text{LOCO}} \ll N$, so the focal SNP projects only weakly onto the LOCO eigenspace. Under approximate isotropy, $E[a_k^2] \approx 1$, $\|\mathbf{a}\|^2 \approx r$, and $\|\mathbf{b}\|^2 \approx N - r$. Substituting into the exact expression gives

$$c_{\text{LOCO}} \approx \frac{1}{N}\sum_{k=1}^{r} \frac{1}{d_k h^2_{\text{LOCO}} + 1 - h^2_{\text{LOCO}}} + \frac{N-r}{N(1-h^2_{\text{LOCO}})}.$$

  As $r/N \to 0$, the second term dominates, so $c_{\text{LOCO}} \to 1/(1-h^2_{\text{LOCO}})$. Thus, when sample size greatly exceeds the dimensionality of the LOCO GRM, LOCO approaches its maximum theoretical advantage.

- $N \gg M_{\text{LOCO}}$**, with pedigree:** Pedigree structure inflates some projections $a_k^2$ on leading eigenvectors with large $d_k$. Because these modes are heavily shrunk, this lowers $c_{\text{LOCO}}$ relative to the weak-pedigree case. However, the reduction is limited if only a small number of leading modes carry strong pedigree structure, while the bulk of the eigenspace behaves approximately like random projection noise. In that case, $c_{\text{LOCO}}$ remains close to $1/(1-h^2_{\text{LOCO}})$, but is systematically smaller than that ideal value.

**Cross-term approximation.** The same analysis applies to cross-terms between focal-chromosome SNPs $\mathbf{x}$ and $\mathbf{z}_s$: $\mathbf{x}^T\hat{\mathbf{V}}^{-1}_{\text{LOCO}}\mathbf{z}_s \approx c_{\text{LOCO}}\, N\, r_{ls}$, with the same $c_{\text{LOCO}}$.

**Expected chi-squared.** Since $E[\mathbf{y}] = \mathbf{0}$ (Section S1.1), $E[(\mathbf{x}^T\hat{\mathbf{V}}^{-1}_{\text{LOCO}}\mathbf{y})^2] = \mathbf{x}^T\hat{\mathbf{V}}^{-1}_{\text{LOCO}}\,\text{Var}(\mathbf{y})\,\hat{\mathbf{V}}^{-1}_{\text{LOCO}}\mathbf{x}$. Substituting $\text{Var}(\mathbf{y}) = \mathbf{Z}_{\text{chr}}\mathbf{W}_{\text{chr}}\mathbf{Z}^T_{\text{chr}}h^2_{\text{chr}}/M_{\text{chr}} + \mathbf{V}_{\text{LOCO}}$ and $\mathbf{V}_{\text{LOCO}} = \hat{\mathbf{V}}_{\text{LOCO}} - \mathbf{I}h^2_{\text{chr}}$ (Section S5.1), and using the cross-term approximation:

$$E(\chi^2) \approx 1 + c_{\text{LOCO}}\, N\, \frac{h^2_{\text{chr}}}{M_{\text{chr}}}\sum_s r^2_{ls}W_{ss}$$

where the first term (from $\mathbf{V}_{\text{LOCO}}$) is approximately 1 since $h^2_{\text{chr}}$ is small.

**S5.3. LOCO NCP** The NCP (excess of $E(\chi^2)$ above 1) is:

$$\boxed{\text{NCP}_{\text{LOCO}} \approx c_{\text{LOCO}}\, N \cdot \frac{h^2_{\text{chr}}}{M_{\text{chr}}} \cdot \sum_s r^2_{ls}W_{ss}}$$

This is Equation 13. The coefficient $c_{\text{LOCO}}$ depends on $N/M_{\text{LOCO}}$ and the pedigree structure (Section S5.2). The upper bound $1/(1-h^2_{\text{LOCO}})$ is approached when $N \gg M_{\text{LOCO}}$ with weak pedigree; pedigree structure reduces $c_{\text{LOCO}}$ below this bound. The NCP grows approximately linearly in $N$, with the growth rate modulated by $c_{\text{LOCO}}$.

**The LOCO NCP uses $W_{ss}$, not $\tilde{W}_{ss}$.** Under LOCO, the focal chromosome's genetic variance $h^2_{\text{chr}}$ is entirely unmodeled, so the test statistic captures the **full** effect of each SNP (including its polygenic component) rather than only the excess above the polygenic expectation. In contrast, the full-GRM NCP (Equation 1) uses $\tilde{W}_{ss} = W_{ss} - 1$ because the polygenic contribution is absorbed by the fitted GRM. This $W_{ss}$-versus-$\tilde{W}_{ss}$ distinction is the algebraic root of the block-level signal phenomenon described in Section S14.

Importantly, the LOCO association statistic does not isolate individual SNP effects. The term $\sum_s r^2_{ls} W_{ss}$ aggregates effects across all SNPs on the focal chromosome that are in LD with the focal SNP. In livestock populations with long-range LD (small $N_e$), this sum extends over large genomic blocks. The LOCO test therefore detects **block-level** genetic signals, not the effect of a single causal variant. This has implications for fine-mapping resolution (Section S12) and the interpretation of GWAS signals (Section S13).

---

### Part II: Properties of the Sigmoid Sum and Practical Saturation

This part establishes the mathematical properties of $S(N) = \sum_{k=1}^{r} \frac{N\ell_k}{N\ell_k + M\lambda}$ (Equation 6).

#### S6. Diminishing Returns in $N$

**Statement.** $S(N)$ is strictly increasing and strictly concave in $N$: each additional sample contributes positively but less than the previous one.

**Proof.** Each term $g_k(N) = N\ell_k/(N\ell_k + M\lambda)$ has first and second derivatives with respect to $N$:

$$g'_k(N) = \frac{M\lambda\ell_k}{(N\ell_k + M\lambda)^2} > 0$$

$$g''_k(N) = \frac{-2M\lambda\ell_k^2}{(N\ell_k + M\lambda)^3} < 0$$

Summing over all $k = 1, \ldots, r$:

$$S'(N) = \sum_{k=1}^{r} \frac{M\lambda\ell_k}{(N\ell_k + M\lambda)^2} > 0, \qquad S''(N) = \sum_{k=1}^{r} \frac{-2M\lambda\ell_k^2}{(N\ell_k + M\lambda)^3} < 0$$

Therefore $S(N)$ is strictly increasing and strictly concave for all $N > 0$.

In modern GWAS, $N > M$ is common (e.g., $N \sim 10^5$ with $M \sim 50{,}000$ chip SNPs), so $r = M$ and the hard upper limit is $S(N) \xrightarrow{N\to\infty} M$. However, since near-zero eigenvalues contribute negligibly at any practically achievable $N$, the effective upper limit is $M_e \ll M$, as established next.

### S7. Practical Saturation at $M_e$

**S7.1. Concavity of the Sigmoid in $\ell$** The sigmoid $g(\ell) = N\ell/(N\ell + M\lambda)$, viewed as a function of $\ell$ at fixed $N$, has second derivative:

$$g''(\ell) = \frac{-2N^2 M\lambda}{(N\ell + M\lambda)^3} < 0$$

so $g(\ell)$ is strictly concave in $\ell$. This concavity enables Jensen's inequality to bound $S(N)$ from above in terms of group-level averages.

**S7.2. Two-Group Decomposition** To analyze saturation as sample size increases, we work in the regime $N > M$, so $r = M$ and all $M$ LD eigenvalues are represented. Partition these $M$ eigenvalues into two groups:

- **Group A**: top $M_e$ eigenvalues, with total mass $\sum_{k\in A} \ell_k = \rho M$ and mean $\bar{\ell}_A = \rho M/M_e$
- **Group B**: remaining $M_B = M - M_e$ eigenvalues, with total mass $\sum_{k\in B} \ell_k = (1-\rho)M$ and mean $\bar{\ell}_B = (1-\rho)M/M_B$

The sigmoid sum then decomposes exactly as:

$$S(N) = S_A(N) + S_B(N)$$

$$S_A(N) = \sum_{k\in A} \frac{N\ell_k}{N\ell_k + M\lambda}, \qquad S_B(N) = \sum_{k\in B} \frac{N\ell_k}{N\ell_k + M\lambda}$$

**S7.3. Jensen Upper Bounds** Applying Jensen's inequality with equal weights $1/M_e$ within Group A:

$$\frac{S_A(N)}{M_e} = \frac{1}{M_e} \sum_{k\in A} g(\ell_k) \le g\left(\frac{1}{M_e} \sum_{k\in A} \ell_k\right) = g(\bar{\ell}_A)$$

Substituting $\bar{\ell}_A = \rho M/M_e$:

$$g(\bar{\ell}_A) = \frac{N \cdot \rho M/M_e}{N \cdot \rho M/M_e + M\lambda} = \frac{N\rho/M_e}{N\rho/M_e + \lambda} = \frac{N\rho}{N\rho + M_e\lambda}$$

Therefore:

$$\boxed{S_A(N) \le U_A(N) \equiv \frac{N\rho\, M_e}{N\rho + M_e\lambda}}$$

Applying the same argument to Group B with equal weights $1/M_B$ and $\bar{\ell}_B = (1-\rho)M/M_B$:

$$\boxed{S_B(N) \le U_B(N) \equiv \frac{N(1-\rho)\,M_B}{N(1-\rho) + M_B\lambda}}$$

Combining: $S(N) \le U_A(N) + U_B(N)$ for all $N \ge M_e$.

**S7.4. Transition Scales** The ceiling of $U_A$ as $N \to \infty$ is $M_e$. The sample size at which $U_A$ reaches half its ceiling is:

$$\boxed{N_A^* = \frac{M_e\,\lambda}{\rho}}$$

Similarly, the ceiling of $U_B(N)$ is $M_B$, and its half-ceiling transition scale is:

$$\boxed{N_B^* = \frac{M_B\,\lambda}{1-\rho}}$$

The ratio of transition scales is:

$$\frac{N_B^*}{N_A^*} = \underbrace{\frac{M_B}{M_e}}_{\text{size factor}} \times \underbrace{\frac{\rho}{1-\rho}}_{\text{mass factor}}$$

Both factors are generally large. The size factor $M_B/M_e = M/M_e - 1$ equals 4–12 for typical livestock chip panels ($M \approx 50{,}000$ and $M_e \approx 4{,}000$–$10{,}000$; Section S15). The mass factor $\rho/(1-\rho)$ equals 49–99 when $\rho = 0.98$–$0.99$. Together, $N_B^*/N_A^*$ ranges from hundreds to about a thousand, placing $N_B^*$ beyond any practically achievable sample size.

**S7.5. Practical Regime: $N_A^* \ll N \ll N_B^*$** **Group A.** As $N \gg N_A^*$, each of the $M_e$ Group A sigmoids approaches 1:

$$S_A(N) \xrightarrow{N\to\infty} M_e$$

In the practical regime $N \gg N_A^*$: $S_A(N) \approx M_e$.

**Group B.** When $N \ll N_B^*$, the average Group B eigenvalue satisfies $N\bar{\ell}_B \ll M\lambda$. In this regime, each Group B sigmoid is well below saturation and approximately linear in $\ell$: $g(\ell) \approx N\ell/M\lambda$. Since $g(\ell)$ is nearly linear, Jensen's inequality is nearly tight, giving $S_B(N) \approx U_B(N)$. Therefore:

$$S_B(N) \approx U_B(N) = \frac{N(1-\rho)M_B}{N(1-\rho) + M_B\lambda} \approx \frac{N(1-\rho)M_B}{M_B\lambda} = \frac{N(1-\rho)}{\lambda}$$

**Bounding $S_B/S_A$.** First, $S_B(N) \approx U_B(N)$, so:

$$\frac{S_B(N)}{S_A(N)} \approx \frac{U_B(N)}{M_e} = \frac{N(1-\rho)}{M_e\lambda}$$

Since $N_A^* = M_e\lambda/\rho$ (Section S7.4), this equals $\frac{1-\rho}{\rho} \cdot \frac{N}{N_A^*}$. Even at $N = 10\,N_A^*$ with $\rho = 0.98$: $S_B/S_A \approx (0.02/0.98) \times 10 \approx 0.2$. At more typical $N/N_A^*$ ratios, $S_B/S_A$ is small.

Therefore throughout the practical regime $N_A^* \ll N \ll N_B^*$:

$$\boxed{S(N) = S_A(N) + S_B(N) \approx M_e}$$

**Small-$N$ regime: $N \ll N_A^*$.** When $N \ll N_A^*$, even the Group A sigmoids are far from saturation. For any eigenvalue with $N\ell_k \ll M\lambda$, the sigmoid is approximately linear: $g(\ell_k) = N\ell_k/(N\ell_k+M\lambda) \approx N\ell_k/M\lambda$. Since this holds for all eigenvalues when $N \ll N_A^*$ (because $N\bar{\ell}_A = N\rho M/M_e \ll M\lambda$ when $N \ll M_e\lambda/\rho = N_A^*$), both groups are in the linear regime:

$$S(N) \approx \frac{N}{M\lambda}\sum_{k=1}^{M} \ell_k = \frac{NM}{M\lambda} = \frac{Nh^2}{1-h^2}$$

In this regime, $S(N)$ is proportional to $N$ and $S_B(N)/S_A(N) \approx (1-\rho)/\rho$, which is small. The corresponding NCP under the $\bar{c}$ approximation is $\text{NCP} = S(N)/M \cdot \sum_s r_{ls}^2 \tilde{W}_{ss} \approx Nh^2/(M(1-h^2)) \cdot \sum_s r_{ls}^2 \tilde{W}_{ss}$, which is Equation 8.

#### S8. The Two Ceilings

The two-group decomposition directly yields two distinct ceilings for $S(N)$:

**Hard ceiling.** For $N > M$, $S(N) \to M$ as $N \to \infty$. Reaching this ceiling requires all eigenvalues, including the near-zero Group B eigenvalues, to saturate, which requires $N \gg N_B^*$.

**Practical ceiling.** For $N$ in the regime $N_A^* \ll N \ll N_B^*$: $S(N) \approx M_e$.

| Ceiling | Value | Transition scale | Typical $N^*$ for livestock ($\rho = 0.98$–$0.99$) |
|---|---|---|---|
| Practical | $M_e$ | $N_A^* = M_e\lambda/\rho$ | $\sim 10^3$–$10^5$ |
| Hard | $M$ | $N_B^* = M_B\lambda/(1-\rho)$ | $\sim 10^6$–$10^7$ |

The practical ceiling $M_e = 4N_eL$ is the operationally relevant upper limit on $S(N)$, and consequently on GWAS power, for any dataset of currently achievable size.

**NCP at each ceiling.** In the ideal tagging case, the NCP under the $\bar{c}$ approximation reduces to $\text{NCP} \approx S(N)(q/h^2 - 1/M)$ (Equation 7; Section S4.4), where $q$ is the focal SNP's PVE.

At the **hard ceiling** ($S(N) \to M$ as $N \gg N_B^*$): $\mathrm{NCP}_{\mathrm{hard}} \approx M(q/h^2 - 1/M) = qM/h^2 - 1$.

At the **practical ceiling** ($S(N) \approx M_e$ for $N_A^* \ll N \ll N_B^*$):

$$\mathrm{NCP}_{\mathrm{practical}} \approx M_e \left( \frac{q}{h^2} - \frac{1}{M} \right) \approx \frac{q \cdot M_e}{h^2}$$

equivalent to Equation 9 evaluated in the ideal tagging case. For common livestock SNP chips, the practical ceiling ($M_e$) is far below the hard ceiling ($M$). At genome-wide significance ($p < 5 \times 10^{-8}$, corresponding to $\chi^2 > 29.7$ for 1 df), NCP $\approx 30$ is required for 50% power. Setting $\mathrm{NCP}_{\mathrm{practical}} \approx 30$, the minimum detectable PVE is:

$$\boxed{q_{\min} \approx \frac{30\, h^2}{M_e}}$$

This is Equation 10.

**Genome-wide significance threshold.** The $p < 5 \times 10^{-8}$ threshold used in the derivation above follows human GWAS convention; $M_e$ in this framework should not be used directly as a multiple-testing divisor. $M_e$ and the effective number of independent tests $M_{\mathrm{eff,test}}$ reflect different summaries of the LD-matrix eigenvalue spectrum: $M_e$ counts only the leading eigenmodes that carry ~98% or ~99% of spectral mass, whereas $M_{\mathrm{eff,test}}$ depends on the count of eigenmodes across the full spectrum, including small-mass tail modes that contribute little to power. Power follows the mass-weighted sum (Equation 6) with a practical ceiling of $M_e$, whereas the family-wise error rate (FWER) across all genome-wide association tests for a trait follows the count of eigenmodes across the spectrum and is unaffected by mass weighting. This distinction has empirical consequences for threshold calibration: FWER calibration in Duroc pig sequence GWAS (Wang *et al.* 2025b) shows that $p < 5 \times 10^{-7}$ achieves FWER $\approx 0.05$ (implying $M_{\mathrm{eff,test}} \sim 10^5$), whereas a naive $0.05/M_e$ correction would be substantially looser and inflate FWER well above 0.05. Therefore the conventional $5 \times 10^{-8}$ threshold remains appropriately conservative for FWER control of livestock sequence GWAS. In humans, the larger $N_e$ produces a flatter LD eigenvalue spectrum, so $M_e$ and $M_{\mathrm{eff,test}}$ can be more similar in magnitude.

**Comparison across GWAS approaches.** In human GWAS, $M_e = 4N_eL \approx 1.4 \times 10^6$ (with $N_e \approx 10{,}000$ and $L \approx 35$ Morgans). With $h^2 = 0.3$, the Group A transition scale is $N_A^* = M_e\lambda/\rho \approx 3.3 \times 10^6$. Most studies have $N < N_A^*$, placing them in the linear regime of $S(N)$ (Section S7.5) where all three approaches (linear regression, LOCO, and full-GRM) scale with $N$. In livestock, $M_e \approx$ 4,000–10,000 (Section S15) and $N_A^* \approx 10^3$–$10^5$, so $N > N_A^*$ can be easily reached and the differences between methods become large.

### S9. Exchange of $N$ and $h^2$ in $S(N)$

The sigmoid sum $S(N) = \sum_k N\ell_k/(N\ell_k + M\lambda)$ depends on $(N, h^2)$ only through one combination (recall $\lambda = (1 - h^2)/h^2$). This section derives the equivalence rigorously and applies it to GWAS using de-regressed breeding values (or de-regressed proofs; DRPs) as pseudo-phenotypes.

**S9.1. Term-by-term invariance** Consider two scenarios $(N_1, h_1^2)$ and $(N_2, h_2^2)$ that share the same population LD-matrix eigenvalues $\{\ell_k\}$. For each $k$, the per-eigenvalue term

$$g_k(N, h^2) := \frac{N\ell_k}{N\ell_k + M(1 - h^2)/h^2}$$

depends on $(N, h^2)$ only through the ratio $\xi_k := N\ell_k/M\lambda = (N\ell_k/M) \cdot h^2/(1 - h^2)$. The two scenarios produce equal $g_k$ iff $\xi_k^{(1)} = \xi_k^{(2)}$, which holds iff

$$N_1 \cdot \frac{h_1^2}{1 - h_1^2} \;=\; N_2 \cdot \frac{h_2^2}{1 - h_2^2}.$$

Because this equality is independent of $\ell_k$, it forces equality of every term simultaneously, and hence $S(N_1; h_1^2) = S(N_2; h_2^2)$ exactly. The conserved quantity is $Nh^2/(1 - h^2)$.

Solving for $N_1$ gives the $N$**-equivalence formula**:

$$N_1 = N_2 \cdot \frac{h_2^2\,(1 - h_1^2)}{h_1^2\,(1 - h_2^2)}.$$

**Population-eigenvalue framing.** The derivation treats $\{\ell_k\}$ as fixed population-level eigenvalues of the LD matrix $\mathbf{R}$. Sample LD matrices computed from $N$ individuals converge to this spectrum at rate $O(1/N)$ in the bulk and faster than that for the top eigenvalues, so the equivalence holds with empirical sample LD matrices whenever both $N_1$ and $N_2$ are large enough for the sample spectrum to approximate the population spectrum (typically $N \gtrsim M_e$). The equivalence is not an asymptotic statement about saturation of $S(N)$; it is an exact algebraic property of the sigmoid sum at any $N$.

**S9.2. Application to DRP-based GWAS** De-regressed breeding values are commonly used as pseudo-phenotypes in livestock GWAS. We require two properties of the DRP transformation: (i) the DRP's effective heritability as a pseudo-phenotype equals its reliability, $h^2_{\text{eff}} = r^2$; and (ii) each causal variant's relative contribution to total genetic variance, $\beta_s^2/\sigma_g^2$, is preserved.

For (i): with $\sigma_g^2$ denoting the additive genetic variance and the DRP's prediction-error variance equal to $\sigma_g^2(1 - r^2)/r^2$ under the standard DRP construction (Garrick *et al.* 2009), the DRP has total variance $\sigma_g^2 + \sigma_g^2(1 - r^2)/r^2 = \sigma_g^2/r^2$, so $h^2_{\text{eff}} = \sigma_g^2/(\sigma_g^2/r^2) = r^2$.

Property (ii) is an assumption about the DRP construction: de-regression rescales noise without redistributing the genetic signal across variants.

Under (i) and (ii), the per-SNP architectural weight $W_{ss} = qM/h^2$ is invariant under DRP (the analysis-scale PVE $q$ and the analysis-scale heritability $h^2$ both transform by the same factor, leaving $q/h^2 = \beta_s^2/\sigma_g^2$ unchanged), and so is $\tilde{W}_{ss} = W_{ss} - 1$ and the aggregate $\sum_s r_{ls}^2 \tilde{W}_{ss}/M$.

Under the $\bar{c}$ approximation (Section S4.3), the NCP takes the $S(N)$ form

$$\mathrm{NCP}_l \approx \frac{S(N)}{M} \sum_s r_{ls}^2 \tilde{W}_{ss}$$

(Equation 5). With the aggregate weight invariant under DRP, the NCP equivalence between an original-phenotype scenario $(N_{\mathrm{equiv}}, h^2_{\mathrm{orig}})$ and a DRP scenario $(N_{\mathrm{DRP}}, r^2)$ reduces to the $S(N)$ equivalence above. Substituting into the $N$-equivalence formula above,

$$N_{\mathrm{equiv}} = N_{\mathrm{DRP}} \cdot \frac{r^2\,(1 - h^2_{\mathrm{orig}})}{h^2_{\mathrm{orig}}\,(1 - r^2)}.$$

This is Equation 11.

The cohort-average $r^2$ is used as a single effective heritability; per-individual reliability heterogeneity is not modeled by this back-of-envelope formula.

**Numerical example (based on the dairy GWAS by Wang *et al.* 2025).** For dairy milk-yield GWAS using $N_{\mathrm{DRP}} \approx$ 50,000 Holstein bulls with average reliability $r^2 \approx 0.8$ and a single-record heritability $h^2_{\mathrm{orig}} \approx 0.2$,

$$N_{\mathrm{equiv}} = 50{,}000 \cdot \frac{0.8 \cdot 0.8}{0.2 \cdot 0.2} = 50{,}000 \cdot 16 = 800{,}000,$$

i.e., per-SNP detection power is equivalent to a single-record-phenotype GWAS of approximately 800,000 genotyped cows.

---

### Part III: Genomic Prediction

This part derives GBLUP reliability from the same eigenvalue framework used for association.

#### S10. Connection to Genomic Prediction

The effective genomic dimensionality that caps association power (Sections S4–S8) also governs genomic prediction. We derive the in-sample reliability of SNP-BLUP (equivalently, GBLUP) from the same eigenvalue spectrum, bound the leave-one-out (out-of-sample) reliability, and bound the optimism between in-sample and out-of-sample reliability, all through the single quantity $S(N)/N$. Throughout, $\lambda = (1 - h^2)/h^2$ and $S(N) = \sum_k N\ell_k/(N\ell_k + M\lambda)$ is the sigmoid sum (Equation 6), with practical ceiling $M_e = 4N_eL$ (Section S7); the $M$ markers densely tag the genome, so $M > M_e$.

**S10.1. Model and SNP-BLUP** We use the working model of Section S1.3: $\mathbf{y} = \mathbf{Z}\boldsymbol{\alpha} + \mathbf{e}$, $\boldsymbol{\alpha} \sim \mathcal{N}(\mathbf{0}, \sigma_\alpha^2\mathbf{I})$, $\mathbf{e} \sim \mathcal{N}(\mathbf{0}, \sigma_e^2\mathbf{I})$, with $h^2 = M\sigma_\alpha^2$ and $\sigma_e^2 = 1 - h^2$ (Section S1.1). SNP-BLUP predicts the marker effects as

$$\hat{\boldsymbol{\alpha}} = (\mathbf{Z}^\top\mathbf{Z} + M\lambda\mathbf{I})^{-1}\mathbf{Z}^\top\mathbf{y} = \mathbf{Z}^\top(\mathbf{Z}\mathbf{Z}^\top + M\lambda\mathbf{I})^{-1}\mathbf{y},$$

the two forms equal by the push-through identity, with ridge $M\lambda = \sigma_e^2/\sigma_\alpha^2$ shrinking each effect in proportion to the noise-to-signal ratio. SNP-BLUP is algebraically equivalent to GBLUP, since $\mathbf{Z}\hat{\boldsymbol{\alpha}} = \mathbf{G}(\mathbf{G} + \lambda\mathbf{I})^{-1}\mathbf{y}$ gives identical predicted genetic values, so we do not distinguish them. The predicted genetic value of an individual with genotype vector $\mathbf{t}$ ($M \times 1$) is $\hat{g} = \mathbf{t}^\top\hat{\boldsymbol{\alpha}}$.

The reliability results below assume the working model (Section S1.3), under which the population's genetic variance is captured by the genome-wide GRM, $\mathrm{Var}(\mathbf{g}) = h^2\mathbf{G}$ (an effectively infinitesimal genetic architecture). This holds for the highly polygenic traits typical of livestock genomic evaluation and most complex human traits. When heritability is instead concentrated in a few large-effect variants, the model is misspecified: GBLUP on a genome-wide GRM predicts poorly relative to variable-selection methods such as BayesB (Daetwyler *et al.* 2010), and the formulas of this section do not apply.

**S10.2. Variance of the estimated SNP effects** Under the working model (Section S1.3), $\mathrm{Var}(\mathbf{y}) = \mathbf{Z}\mathbf{Z}^\top\sigma_\alpha^2 + \mathbf{I}\sigma_e^2 = \sigma_\alpha^2(\mathbf{Z}\mathbf{Z}^\top + M\lambda\mathbf{I})$, so the sandwich form of the variance collapses:

$$\mathrm{Var}(\hat{\boldsymbol{\alpha}}) = \sigma_\alpha^2\,\mathbf{Z}^\top(\mathbf{Z}\mathbf{Z}^\top + M\lambda\mathbf{I})^{-1}\mathbf{Z} =: \sigma_\alpha^2\,\mathbf{S}_\Phi,$$

defining the shrinkage matrix

$$\mathbf{S}_\Phi = \mathbf{Z}^\top(\mathbf{Z}\mathbf{Z}^\top + M\lambda\mathbf{I})^{-1}\mathbf{Z} = (\mathbf{Z}^\top\mathbf{Z} + M\lambda\mathbf{I})^{-1}\mathbf{Z}^\top\mathbf{Z} \quad (M \times M),$$

again by the push-through identity. Its nonzero eigenvalues lie in $(0, 1)$, with the remaining eigenvalues equal to zero when $\mathbf{Z}$ has rank below $M$; consequently, the estimated effects carry less variance than their prior $\sigma_\alpha^2\mathbf{I}$, by a factor set by the signal-to-ridge ratio.

Inserting the thin SVD $\mathbf{Z} = \mathbf{U}\boldsymbol{\Delta}\boldsymbol{\Phi}^\top$ (Section S1.2) and simplifying as in Sections S2.1 and S2.3:

$$\mathbf{S}_\Phi = \boldsymbol{\Phi}\,\mathrm{diag}(g_k)\,\boldsymbol{\Phi}^\top, \qquad g_k = \frac{\Delta_k^2}{\Delta_k^2 + M\lambda} = \frac{N\ell_k}{N\ell_k + M\lambda},$$

with $\ell_k = \Delta_k^2/N$ the LD-matrix eigenvalues (Section S1.4). The $g_k$'s are exactly the per-mode sigmoid terms of $S(N)$ (Equation 6): the precision of each estimated effect is inherited from the same spectrum that builds the association NCP.

Using $\mathbf{\Phi}^\top\mathbf{\Phi} = \mathbf{I}$,

$$\mathrm{tr}(\mathbf{S}_\Phi) = \mathrm{tr}(\mathbf{\Phi}\,\mathrm{diag}(g_k)\,\mathbf{\Phi}^\top) = \mathrm{tr}(\mathbf{\Phi}^\top\mathbf{\Phi}\,\mathrm{diag}(g_k)) = \sum_k g_k = S(N).$$

Thus $S(N)$ is the effective number of parameters SNP-BLUP resolves, its effective degrees of freedom in the ridge-regression sense (Hastie *et al.* 2009): each mode contributes a fractional parameter $g_k \in (0,1)$, fully estimated when $N\ell_k \gg M\lambda$ and shrunk away when $N\ell_k \ll M\lambda$. By Section S7, $S(N) \le M_e$ in the practical regime, so SNP-BLUP resolves at most about $M_e$ effective parameters regardless of the marker count $M$.

**S10.3. Reliability for a target genotype vector** For a target with genotype vector $\mathbf{t}$ ($M \times 1$), the true genetic value is $g = \mathbf{t}^\top\boldsymbol{\alpha}$ (a genetic value, distinct from the per-mode term $g_k$) and the prediction is $\hat{g} = \mathbf{t}^\top\hat{\boldsymbol{\alpha}}$. Reliability is the squared correlation between the two; for BLUP $\mathrm{Cov}(g,\hat{g}) = \mathrm{Var}(\hat{g})$, so

$$R^2(\mathbf{t}) = \frac{\mathrm{Var}(\hat{g})}{\mathrm{Var}(g)} = \frac{\mathbf{t}^\top\mathbf{S}_\Phi\mathbf{t}}{\mathbf{t}^\top\mathbf{t}}.$$

Split $\mathbf{t}$ into its part inside the SNP-eigenspace $\mathrm{col}(\mathbf{\Phi})$ and the orthogonal remainder,

$$\mathbf{t} = \mathbf{\Phi}\mathbf{p} + \mathbf{q}, \qquad \mathbf{p} = \mathbf{\Phi}^\top\mathbf{t}, \qquad \mathbf{q} = (\mathbf{I} - \mathbf{\Phi}\mathbf{\Phi}^\top)\mathbf{t},$$

the SNP-space counterpart of the focal-SNP split $\mathbf{x} = \mathbf{U}\mathbf{a} + \mathbf{b}$ (Section S2.2). Because $\mathbf{S}_\Phi$ acts only within $\mathrm{col}(\mathbf{\Phi})$, $\mathbf{\Phi}^\top\mathbf{t} = \mathbf{p}$ and

$$R^2(\mathbf{t}) = \frac{\sum_k g_k\, p_k^2}{\mathbf{p}^\top\mathbf{p} + \mathbf{q}^\top\mathbf{q}}, \qquad \mathbf{t}^\top\mathbf{t} = \mathbf{p}^\top\mathbf{p} + \mathbf{q}^\top\mathbf{q}.$$

The orthogonal part $\mathbf{q}$ adds to the true value but not the prediction: it is the component of the target outside the directions the training data span, and is intrinsically unpredictable. When $\mathbf{q} = \mathbf{0}$, which holds for every training individual and for any target when $N > M$ (Section S10.5), the target is fully spanned and $\mathbf{t}^\top\mathbf{t} = \mathbf{p}^\top\mathbf{p}$. Per-SNP standardization (Section S1.1) gives $\mathbf{t}^\top\mathbf{t} = M$ in expectation.

**S10.4. In-sample reliability** A training individual $i$ has genotype vector $\mathbf{t}_i = \mathbf{Z}^\top\mathbf{e}_i = \mathbf{\Phi}\mathbf{\Delta}\mathbf{U}^\top\mathbf{e}_i$ ($M \times 1$; Section S1.2), so $\mathbf{p}_i = \mathbf{\Delta}\mathbf{U}^\top\mathbf{e}_i$ and $\mathbf{q}_i = \mathbf{0}$, a training individual lying wholly in the row space of $\mathbf{Z}$. Then $p_{ik}^2 = \Delta_k^2 U_{ik}^2$, and averaging predicted and true variances over the $N$ training individuals (the pooled reliability, using $\sum_i U_{ik}^2 = 1$, $\Delta_k^2 = N\ell_k$, and mean $\mathbf{t}_i^\top\mathbf{t}_i = M$), with $g_k$ the per-mode term in $S(N)$ (Section S10.2):

$$\bar{R}^2_{\mathrm{in}} = \frac{1}{M}\sum_k g_k\,\ell_k = \frac{1}{M}\sum_k \frac{N\ell_k^2}{N\ell_k + M\lambda} = \frac{1}{N}\sum_k \frac{d_k^2}{d_k + \lambda} = 1 - \frac{\lambda S(N)}{N},$$

the in-sample average reliability, namely the fraction of genetic variance recovered by GBLUP within the training set (the $d_k = N\ell_k/M$ are the GRM eigenvalues, with $\sum_k d_k = N$). The last form uses $\frac{d_k^2}{d_k+\lambda} = d_k - \lambda\frac{d_k}{d_k+\lambda}$ and $\sum_k \frac{d_k}{d_k+\lambda} = \sum_k g_k = S(N)$, with $\sum_k d_k = N$, tying the in-sample reliability directly to the detection sum $S(N)$.

**Lower bound and approximation.** Write $\bar{R}^2_{\text{in}} = \frac{1}{M}\sum_k f(\ell_k)$ with $f(\ell) = N\ell^2/(N\ell + M\lambda)$, convex since $f''(\ell) = 2N(M\lambda)^2/(N\ell + M\lambda)^3 > 0$. The sample LD matrix has rank $\min(N, M)$, so for $N < M$ there are $N$ nonzero eigenmodes, with $\sum_k \ell_k = M$ (recall $M > M_e$). When $N < M_e$, Jensen's inequality across all $N$ nonzero eigenmodes gives only the trivial bound $\bar{R}^2_{\text{in}} \geq \frac{N}{M} f\left(\frac{1}{N}\sum_k \ell_k\right) = \frac{1}{1+\lambda} = h^2$ (the in-sample reliability is at least $h^2$, with equality when the $N$ GRM eigenvalues are all 1, that is $\mathbf{G} = \mathbf{I}$). When $N \geq M_e$, a sharper bound follows: partition the eigenmodes into Group A (the leading $M_e$, with mass $\sum_{k\in A} \ell_k = \rho M$) and Group B (the remainder), as in Section S7.2. Because $f \geq 0$, dropping Group B and applying Jensen's inequality to the convex $f$ over Group A gives the lower bound:

$$\bar{R}^2_{\text{in}} \geq \frac{1}{M}\sum_{k\in A} f(\ell_k) \geq \frac{M_e}{M} f\left(\frac{\rho M}{M_e}\right) = \frac{N\rho^2}{N\rho + M_e\lambda}.$$

For $\rho \to 1$ this is $Nh^2/[Nh^2 + M_e(1 - h^2)]$, Equation 18. Both inequalities are nearly tight in the regime $N_A^* \ll N \ll N_B^*$ (Section S7.5), so Equation 18 closely approximates $\bar{R}^2_{\text{in}}$ there: when $N \ll N_B^*$, dropping Group B loses little, and even less than the analogous step for $S(N)$, since each dropped term $f(\ell_k) = \ell_k g_k$ carries the small eigenvalue $\ell_k$ both as a weight and within $g_k$; when $N_A^* \ll N$, that is $N \gg M_e\lambda$, the average Group A eigenvalue satisfies $N\bar{\ell}_A = N\rho M/M_e \gg M\lambda$, so $f(\ell) \approx \ell - M\lambda/N$ is nearly linear over Group A and the Jensen gap is small. For $N < N_A^*$ the bound is loose.

**S10.5. Out-of-sample reliability** For an out-of-sample target the reliability averages $R^2(\mathbf{t})$ over the target's genotype distribution. Using $E[\mathbf{t}^\top \mathbf{S}_\Phi \mathbf{t}] = \text{tr}(\mathbf{S}_\Phi \boldsymbol{\Sigma})$ (Section S10.3) and $E[\mathbf{t}^\top \mathbf{t}] = \text{tr}(\boldsymbol{\Sigma}) = M$, with $\boldsymbol{\Sigma} = E[\mathbf{t}\mathbf{t}^\top]$ the population LD matrix,

$$\bar{R}^2_{\text{out}} = \frac{1}{M}\text{tr}(\mathbf{S}_\Phi \boldsymbol{\Sigma}).$$

We assume the $M$ markers adequately tag the genome (Section S10.1), so that spanning the marker space captures the genetic value; the regimes below concern whether the training data span it. Out-of-sample reliability can fall below in-sample for two distinct reasons: the target may lie partly outside the trained subspace (a nonzero $\mathbf{q}$, Section S10.3), and, even when fully spanned, the target's own record is absent from the fit. The first is governed by the sample size relative to the marker count $M$ and the effective dimension $M_e$:

- $\mathbf{q} = \mathbf{0}$ exactly. In-sample targets always satisfy this, and when $N > M$ with distinct individuals,

$\mathbf{\Phi}$ is $M \times M$ orthonormal, so $\text{col}(\mathbf{\Phi}) = \mathbb{R}^M$ and every target is fully spanned.

- $\mathbf{q} \approx \mathbf{0}$. When $N < M$ but $N \gtrsim M_e$, the training spans the $M_e$ leading directions that carry essentially all genomic variance (Section S7), so a target's unspanned component lies only in negligible-mass tail modes. A reference chosen to represent the population well falls here; this is the condition met by APY core animals, whose core size is set to $n_c \approx M_e$ (Pocrnic *et al.* 2016).
- $\mathbf{q} \neq \mathbf{0}$ almost surely. When $N < M_e$, the training cannot span the effective dimensions, so a new target retains a substantial unpredictable component, and out-of-sample reliability falls below in-sample. In the extreme $\mathbf{t} \perp \text{col}(\mathbf{\Phi})$, so $\mathbf{p} = \mathbf{0}$, one has $R^2(\mathbf{t}) = 0$.

Given $\mathbf{q} \approx \mathbf{0}$, the remaining difference between out-of-sample and in-sample reliability is the gain from a training individual's own record being in the fit, quantified exactly in Section S10.6.

**S10.6. Leave-one-out reliability and in-sample optimism** The leave-one-out (LOO) reliability (the reliability of predicting each training individual from the rest of the training data) is the out-of-sample reliability of Section S10.5 realized within the sample. We compute it, recover the in-sample reliability of Section S10.4 by an independent route, and bound both the leave-one-out reliability and the optimism of in-sample over it.

We work in the equivalent GRM space, with the GRM $\mathbf{G} = \mathbf{Z}\mathbf{Z}^\top/M$ ($N \times N$) and $\mathbf{A} = \mathbf{G} + \lambda\mathbf{I}$. The GBLUP predictor is $\hat{\mathbf{g}} = \mathbf{H}\mathbf{y}$ with the hat matrix $\mathbf{H} = \mathbf{G}\mathbf{A}^{-1}$. From $\mathbf{H} = \mathbf{I} - \lambda\mathbf{A}^{-1}$,

$$(\mathbf{A}^{-1})_{ii} = \frac{1 - H_{ii}}{\lambda},$$

where $H_{ii} = [\mathbf{G}(\mathbf{G} + \lambda\mathbf{I})^{-1}]_{ii}$ is the leverage of individual $i$, its own record's influence on its own prediction. The eigenvalues of $\mathbf{H}$ are $d_k/(d_k + \lambda) = g_k$ (the per-mode term in $S(N)$), identical to those of $\mathbf{S}_\Phi$ (Section S10.2), so $\sum_i H_{ii} = \text{tr}(\mathbf{H}) = \sum_k g_k = S(N)$. Because $\mathbf{G} \succeq \mathbf{0}$ and $\lambda > 0$, these eigenvalues lie in $[0, 1)$, so $\mathbf{0} \preceq \mathbf{H} \prec \mathbf{I}$ and $0 \le H_{ii} < 1$ for every individual $i$.

**In-sample reliability.** The in-sample predicted values $\hat{\mathbf{g}} = \mathbf{H}\mathbf{y}$ have $\text{Var}(\hat{\mathbf{g}}) = h^2\mathbf{G}\mathbf{A}^{-1}\mathbf{G}$ (using $\text{Var}(\mathbf{y}) = h^2\mathbf{A}$), and $\mathbf{G}\mathbf{A}^{-1}\mathbf{G} = \mathbf{G}(\mathbf{I} - \lambda\mathbf{A}^{-1}) = \mathbf{G} - \lambda\mathbf{H}$. With $\text{Var}(g_i) = h^2 G_{ii}$ for the genetic value $g_i$ (distinct from the per-mode term $g_k$), the per-individual in-sample reliability (as in Section S10.3) is

$$R^2_{\text{in},i} = \frac{(\mathbf{G}\mathbf{A}^{-1}\mathbf{G})_{ii}}{G_{ii}} = 1 - \frac{\lambda H_{ii}}{G_{ii}}.$$

Averaging predicted and true variances over individuals (the pooled reliability), with $\overline{G_{ii}} = \frac{1}{N}\sum_i G_{ii} = 1$ by standardization (the inbreeding coefficient does not enter) and $\overline{H_{ii}} = \frac{1}{N}\sum_i H_{ii} = S(N)/N$,

$$\bar{R}^2_{\text{in}} = 1 - \lambda\frac{\overline{H_{ii}}}{\overline{G_{ii}}} = 1 - \frac{\lambda S(N)}{N},$$

recovering the in-sample reliability of Section S10.4 by an independent route.

**Leave-one-out reliability.** Predicting $i$ from the others gives $\hat{g}_i^{(-i)} = \mathbf{G}_{i,-i}\mathbf{A}_{-i}^{-1}\mathbf{y}_{-i}$ with $\mathbf{A}_{-i} = \mathbf{G}_{-i,-i} + \lambda\mathbf{I}$, of variance $h^2\mathbf{G}_{i,-i}\mathbf{A}_{-i}^{-1}\mathbf{G}_{-i,i}$. By the Schur-complement identity, $(\mathbf{A}^{-1})_{ii} = [G_{ii} + \lambda - \mathbf{G}_{i,-i}\mathbf{A}_{-i}^{-1}\mathbf{G}_{-i,i}]^{-1}$, so $\mathbf{G}_{i,-i}\mathbf{A}_{-i}^{-1}\mathbf{G}_{-i,i} = G_{ii} + \lambda - 1/(\mathbf{A}^{-1})_{ii}$ and

$$R^2_{\mathrm{LOO},i} = \frac{\mathbf{G}_{i,-i}\mathbf{A}_{-i}^{-1}\mathbf{G}_{-i,i}}{G_{ii}} = 1 + \frac{\lambda}{G_{ii}} - \frac{\lambda}{G_{ii}(1 - H_{ii})}.$$

Because $\mathbf{A}_{-i} = \mathbf{G}_{-i,-i}+\lambda\mathbf{I}$ is positive-definite, the quadratic form $\mathbf{G}_{i,-i}\mathbf{A}_{-i}^{-1}\mathbf{G}_{-i,i} \geq 0$, so $R^2_{\mathrm{LOO},i} \geq 0$, with equality only for an isolated individual ($\mathbf{G}_{i,-i} = \mathbf{0}$). Hence the average leave-one-out reliability $\bar{R}^2_{\mathrm{LOO}} \geq 0$, reaching 0 only when every individual is isolated ($\mathbf{G} \approx \mathbf{I}$), exactly as expected. The leverage shortcut behind efficient leave-one-out cross-validation for GBLUP (Cheng et al. 2017) is the same one we use here to derive the expected reliability.

**The optimism.** Subtracting,

$$R^2_{\mathrm{in},i} - R^2_{\mathrm{LOO},i} = \frac{\lambda H_{ii}^2}{G_{ii}(1 - H_{ii})}.$$

Averaging over individuals,

$$\Delta_{\mathrm{opt}} = \bar{R}^2_{\mathrm{in}} - \bar{R}^2_{\mathrm{LOO}} = \frac{1}{N}\sum_{i=1}^{N} \frac{\lambda H_{ii}^2}{G_{ii}(1 - H_{ii})}.$$

For the upper bound, the Schur-complement identity gives $(\mathbf{A}^{-1})_{ii} = [G_{ii} + \lambda - \mathbf{G}_{i,-i}\mathbf{A}_{-i}^{-1}\mathbf{G}_{-i,i}]^{-1} \geq 1/(G_{ii}+\lambda)$, since $\mathbf{G}_{i,-i}\mathbf{A}_{-i}^{-1}\mathbf{G}_{-i,i} \geq 0$, so the leverage obeys $H_{ii} = 1-\lambda(\mathbf{A}^{-1})_{ii} \leq G_{ii}/(G_{ii}+\lambda)$, with equality only for an isolated individual ($\mathbf{G}_{i,-i}\mathbf{A}_{-i}^{-1}\mathbf{G}_{-i,i} = 0$). Equivalently $\lambda H_{ii}/(1 - H_{ii}) \leq G_{ii}$, so each optimism term satisfies $\frac{\lambda H_{ii}^2}{G_{ii}(1-H_{ii})} = \frac{H_{ii}}{G_{ii}} \cdot \frac{\lambda H_{ii}}{1-H_{ii}} \leq H_{ii}$, and summing $\sum_i H_{ii} = S(N)$ gives $\Delta_{\mathrm{opt}} \leq S(N)/N$. For the lower bound, write $\phi(h) = h^2/(1 - h)$ (convex on $[0, 1)$ since $\phi''(h) = 2(1 - h)^{-3} > 0$), so that $\Delta_{\mathrm{opt}} = \frac{\lambda}{N}\sum_i \phi(H_{ii})/G_{ii}$. When the diagonal $G_{ii}$ is concentrated about its mean, the per-individual ratios pool, $\frac{1}{N}\sum_i \phi(H_{ii})/G_{ii} \approx \overline{\phi(H_{ii})}/\overline{G_{ii}}$ (that is, the mean of the ratios approximately equal to the ratio of the means); therefore, with $\overline{G_{ii}} = 1$ by standardization, $\Delta_{\mathrm{opt}} \approx \frac{\lambda}{N}\sum_i \phi(H_{ii})$. Jensen's inequality then gives $\frac{1}{N}\sum_i \phi(H_{ii}) \geq \phi(S(N)/N)$, with equality at uniform leverage. Combining,

$$\lambda\,\frac{(S(N)/N)^2}{1 - S(N)/N} \;\leq\; \Delta_{\mathrm{opt}} \;\leq\; \frac{S(N)}{N}.$$

**Bounds of leave-one-out reliability.** Since $\bar{R}^2_{\mathrm{LOO}} = \bar{R}^2_{\mathrm{in}} - \Delta_{\mathrm{opt}}$ with $\bar{R}^2_{\mathrm{in}} = 1 - \lambda S(N)/N$, the optimism bracket gives

$$1 - (\lambda + 1)\frac{S(N)}{N} \;\leq\; \bar{R}^2_{\mathrm{LOO}} \;\leq\; 1 - \frac{\lambda S(N)/N}{1 - S(N)/N}.$$

The lower bound is non-negative because $S(N)/N \leq 1/(1+\lambda) = h^2$: for $N \leq M$ the sample GRM has $N$ eigenvalues averaging 1, so Jensen's inequality on the concave $d/(d+\lambda)$ gives $S(N)/N \leq 1/(1+\lambda) = h^2$, and since $S(N)/N = \sum_k \ell_k/(N\ell_k + M\lambda)$ decreases in $N$, the bound persists for $N > M$. The lower bound thus sharpens the exact $\bar{R}^2_{\text{LOO}} \geq 0$. The in-sample and leave-one-out reliabilities approach 1 at different scales: $\bar{R}^2_{\text{in}} = 1 - \lambda S(N)/N$ is close to 1 once $N \gg M_e\lambda$, whereas the leave-one-out lower bound $1 - (\lambda+1)S(N)/N = 1 - S(N)/(Nh^2)$ requires the larger $N \gg M_e/h^2$. Genomic evaluation programs in livestock can practically reach this larger $N$, particularly for moderate- to high-heritability traits, so both reliabilities approach 1. The upper bound is attained at uniform leverage and mirrors $\bar{R}^2_{\text{in}} = 1 - \lambda S(N)/N$ with an extra factor $1/(1 - S(N)/N)$, so out-of-sample reliability always sits below in-sample.

These results rest on the exact GBLUP model: the per-individual identities are algebraic, and the optimism upper bound $\Delta_{\text{opt}} \leq S(N)/N$ holds for any relatedness structure. A main assumption underlying the closed-form bounds is that the diagonal $G_{ii}$ is concentrated about its mean, so that the per-individual reliabilities pool. In practice, the GRM's diagonal elements can average above 1, because of inbreeding and the particular genotype scaling (e.g., VanRaden 2008); rescaling the whole GRM by its mean diagonal then matches the unit-mean diagonal used in our derivation. No independence assumption or large-sample approximation enters.

**S10.7. Interpretation** The single quantity $S(N)/N$ governs both reliabilities (in-sample $\bar{R}^2_{\text{in}} = 1 - \lambda S(N)/N$ and out-of-sample $\bar{R}^2_{\text{LOO}}$ bounded above), and its size is set by where $N$ falls relative to $M_e$, sharply separating livestock from humans.

**Livestock.** Because $M_e$ is small in livestock ($\sim 10^3$–$10^4$), well-established genomic evaluation programs reach $N \gg M_e/h^2$, where $S(N) \to M_e$ (Section S7) and $S(N)/N \approx M_e/N \ll h^2$: both $\bar{R}^2_{\text{in}}$ and $\bar{R}^2_{\text{LOO}}$ are close to 1 and the in-sample optimism is only $\approx \lambda(M_e/N)^2$ and at most $M_e/N$. Prediction is reliable and transfers out of sample.

**Humans.** Because $M_e$ is large in humans ($\sim 10^6$), even biobank-scale $N$ is generally below $M_e$. The $N$ GRM eigenvalues average 1, so $S(N)/N \leq h^2$ (Section S10.6), of order $h^2$ rather than the tiny $M_e/N$ of livestock. To estimate it, model the GRM as a Wishart matrix with $M_e$ independent dimensions, whose eigenvalues follow a Marchenko–Pastur distribution with aspect ratio $\gamma = N/M_e$. Then $S(N)/N \approx E_{\text{MP}}[d/(d+\lambda)]$, slightly below $h^2$ because $d/(d+\lambda)$ is concave. This is a heuristic illustration for the high-dimensional human regime. Real spectra carry additional structure (population-structure eigenvalues above the bulk), but the order of magnitude is robust. For $N = 5 \times 10^5$ and $M_e = 1.4 \times 10^6$ ($\gamma = 0.36$), the Marchenko–Pastur estimate gives $S(N)/N \approx 0.10$, 0.28, and 0.46 at $h^2 = 0.1$, 0.3, and 0.5, respectively; therefore, in-sample $\bar{R}^2_{\text{in}} \approx 0.13$, 0.35, and 0.54 but out-of-sample $\bar{R}^2_{\text{LOO}} \approx 0.03$, 0.10, and 0.16 (uniform-leverage upper bounds), respectively. In-sample reliability is moderate and out-of-sample small. In the limit $N \ll M_e$ ($\gamma \ll 1$), the Marchenko–Pastur eigenvalues concentrate near 1 with variance $\gamma$, and a second-order expansion of $d/(d+\lambda)$ about 1 gives $S(N)/N \approx h^2 - (1-h^2)h^4\gamma$. The uniform-leverage upper bound on $\bar{R}^2_{\text{LOO}}$ (Section S10.6) vanishes at $S(N)/N = h^2$, so to leading order $\bar{R}^2_{\text{LOO}} \lesssim h^2\gamma = Nh^2/M_e$, exactly the small-$N$ limit of the Daetwyler reliability $Nh^2/(Nh^2+M_e)$ (Daetwyler *et al.* 2008). Out-of-sample reliability is therefore limited by the information ratio $N/M_e$ in this regime. These are GBLUP reliabilities with genome-wide SNPs, performing well for an effectively infinitesimal genetic architecture (Section S10.1). For sparse architectures, variable-selection methods can exceed them.

**Contrast.** The same effective dimensionality governs both regimes: small $M_e$ in livestock makes $S(N)/N$ tiny with practically achievable $N$, so prediction can be easy and robust; large $M_e$ in humans keeps $S(N)/N$ of order $h^2$ even at biobank scale, so prediction is intrinsically demanding. Prediction is comparatively easy precisely where SNP-level mapping is hard.

---

### Part IV: Fine-Mapping Resolution Limit

This part extends the NCP framework to fine-mapping, showing that the resolution-governing factor $(1-r^2_{ij})$ links fine-mapping to GWAS power and genomic prediction through the common quantity $N_e$ (Equations 14–16).

#### S11. Fine-Mapping as Model Comparison

**S11.1. The Fine-Mapping Problem** Given a genomic region containing $K$ candidate SNPs in LD, fine-mapping asks: **which subset of SNPs have non-zero effects?** In the simplest case, we compare single-SNP causal models. For candidate SNP $l$, the single-SNP model is:

$$\mathcal{M}_l : \quad \mathbf{y} = \mathbf{z}_l \beta_l + \boldsymbol{\epsilon}$$

where $\boldsymbol{\epsilon}$ absorbs all other genetic and environmental effects. The maximum likelihood estimate is $\hat{\beta}_l = \mathbf{z}_l^T \mathbf{y}/N$ (since $\mathbf{z}_l^T \mathbf{z}_l = N$ for standardized genotypes), and the profile log-likelihood (up to a constant shared across models) is:

$$\ell_l = \frac{(\mathbf{z}_l^T \mathbf{y})^2}{2N\sigma^2_\epsilon}$$

**S11.2. Expected Log-Likelihood Difference Under Simple Regression** Suppose the true causal SNP is $i$ with effect $\beta_i$, so $E[\mathbf{y}] = \mathbf{z}_i \beta_i$ and $\text{Var}(\mathbf{y}) = \sigma^2_\epsilon \mathbf{I}$. For any candidate SNP $l$:

$$E[\mathbf{z}_l^T \mathbf{y}] = \beta_i \, \mathbf{z}_l^T \mathbf{z}_i = N r_{li} \beta_i$$

$$\text{Var}(\mathbf{z}_l^T \mathbf{y}) = \sigma^2_\epsilon \, \mathbf{z}_l^T \mathbf{z}_l = N\sigma^2_\epsilon$$

Therefore:

$$E[(\mathbf{z}_l^T \mathbf{y})^2] = N^2 r_{li}^2 \beta_i^2 + N\sigma_\epsilon^2$$

The expected log-likelihood difference between the true model ($l = i$) and an alternative model ($l = j$) is:

$$E[\ell_i - \ell_j] = \frac{1}{2N\sigma_\epsilon^2}\left(N^2\beta_i^2(1 - r_{ij}^2)\right) = \frac{N\beta_i^2}{2\sigma_\epsilon^2}(1 - r_{ij}^2)$$

since $r_{ii} = 1$. For standardized phenotypes ($\sigma_\epsilon^2 \approx 1$):

$$\boxed{E[\Delta\ell_{i\to j}] = \frac{N\beta_i^2}{2}(1 - r_{ij}^2)}$$

This is Equation 14. The expected evidence favoring the true causal SNP depends on sample size $N$, effect size $\beta_i^2$ (per-SNP PVE), and the LD imperfection $1 - r_{ij}^2$.

**S12. Fine-Mapping Under Mixed Models**

The simple regression result of Section S11 assumes $\text{Var}(\mathbf{y}) = \sigma_\epsilon^2\mathbf{I}$. In practice, fine-mapping is conducted after fitting a mixed model. We now derive the expected log-likelihood difference under both LOCO and full-GRM mixed models.

**S12.1. General GLS Framework for Model Comparison** Under a mixed model with phenotypic covariance $\mathbf{V}$, the GLS log-likelihood for candidate SNP $l$ (up to constants shared across models) is:

$$\ell_l^{\text{GLS}} = \frac{(\mathbf{z}_l^T\mathbf{V}^{-1}\mathbf{y})^2}{2\,\mathbf{z}_l^T\mathbf{V}^{-1}\mathbf{z}_l}$$

This is proportional to the score chi-squared statistic. The expectation decomposes as:

$$E[\ell_l^{\text{GLS}}] = \frac{\text{Var}(\mathbf{z}_l^T\mathbf{V}^{-1}\mathbf{y}) + (E[\mathbf{z}_l^T\mathbf{V}^{-1}\mathbf{y}])^2}{2\,\mathbf{z}_l^T\mathbf{V}^{-1}\mathbf{z}_l}$$

Under the working model, if $\text{Var}(\mathbf{y}) = \mathbf{V}$ exactly, then

$$\text{Var}(\mathbf{z}_l^T\mathbf{V}^{-1}\mathbf{y}) = \mathbf{z}_l^T\mathbf{V}^{-1}\mathbf{V}\mathbf{V}^{-1}\mathbf{z}_l = \mathbf{z}_l^T\mathbf{V}^{-1}\mathbf{z}_l,$$

so the variance contribution to $E[\ell_l^{\text{GLS}}]$ is exactly $\frac{1}{2}$ for every candidate model. Under the true generative model, $\text{Var}(\mathbf{y}) = \mathbf{Z}\mathbf{W}\mathbf{Z}^T\sigma_\alpha^2 + \mathbf{I}\sigma_e^2$ need not equal $\mathbf{V}$ when $\mathbf{W} \neq \mathbf{I}$. For the present approximation, we assume the working-model covariance is adequate so that this cancellation remains approximately valid. To cancel the variance terms in $E[\ell_i - \ell_j]$, we additionally need $\mathbf{z}_i^T\mathbf{V}^{-1}\mathbf{z}_i \approx \mathbf{z}_j^T\mathbf{V}^{-1}\mathbf{z}_j$. This is justified by the GRAMMAR-Gamma approximation: for GRM SNPs, $\mathbf{z}_l^T\mathbf{V}^{-1}\mathbf{z}_l \approx \bar{c}N$ (Section S4.3), and for within-block SNPs $i$ and $j$ (similar MAF and LD environment), the approximation is tighter still. The empirical coefficient of variation (CV) of $c_l$ across real livestock chip panels is well below 0.3, supporting this approximation (main text Section 4.1).

Under this approximation, the variance terms cancel and the expected signal is $E[\mathbf{z}_l^T\mathbf{V}^{-1}\mathbf{y}] = \beta_i\,\mathbf{z}_l^T\mathbf{V}^{-1}\mathbf{z}_i$. The expected log-likelihood difference is therefore:

$$\boxed{E[\Delta\ell_{i\to j}^{\text{GLS}}] = \frac{\beta_i^2}{2}\left[\mathbf{z}_i^T\mathbf{V}^{-1}\mathbf{z}_i - \frac{(\mathbf{z}_j^T\mathbf{V}^{-1}\mathbf{z}_i)^2}{\mathbf{z}_j^T\mathbf{V}^{-1}\mathbf{z}_j}\right]}$$

This is the general result. The specific form of $\mathbf{V}$ determines the fine-mapping resolution under each analysis strategy.

**S12.2. LOCO Mixed Model** Under LOCO, by the same analysis as in Section S5.2, $\hat{\mathbf{V}}_{\text{LOCO}}^{-1}$ acts on focal-chromosome genotype vectors with effective coefficient $c_{\text{LOCO}}$ (Section S5.2):

$$\mathbf{z}_l^T\hat{\mathbf{V}}_{\text{LOCO}}^{-1}\mathbf{z}_l \approx c_{\text{LOCO}}\,N, \qquad \mathbf{z}_j^T\hat{\mathbf{V}}_{\text{LOCO}}^{-1}\mathbf{z}_i \approx c_{\text{LOCO}}\,N\,r_{ji}$$

Substituting into the general formula (Section S12.1):

$$E[\Delta\ell_{i\to j}^{\text{LOCO}}] = \frac{\beta_i^2}{2}\left[c_{\text{LOCO}}N - \frac{(c_{\text{LOCO}}Nr_{ij})^2}{c_{\text{LOCO}}N}\right] = \frac{\beta_i^2}{2}\cdot c_{\text{LOCO}}\,N\,(1-r_{ij}^2)$$

Therefore:

$$\boxed{E[\Delta\ell_{i\to j}^{\text{LOCO}}] \approx \frac{c_{\text{LOCO}}\,N\,\beta_i^2}{2}(1-r_{ij}^2)}$$

When $N \gg M_{\text{LOCO}}$ with weak pedigree, $c_{\text{LOCO}} \approx 1/(1-h_{\text{LOCO}}^2)$ and LOCO provides a factor of $1/(1-h_{\text{LOCO}}^2)$ improvement over simple regression. With strong pedigree or $N$ comparable to $M_{\text{LOCO}}$, $c_{\text{LOCO}}$ is smaller than this ideal value and the improvement is reduced (Section S5.2).

**S12.3. Full-GRM Mixed Model** Under the full-GRM, $\mathbf{V} = \mathbf{G}h^2 + \mathbf{I}(1-h^2)$ where $\mathbf{G}$ includes all SNPs, including the focal region. The focal SNPs now lie **within** the column space of the GRM.

For fine-mapping, the relevant SNP pairs $i$ and $j$ are in strong LD ($r_{ij}^2$ close to 1), so $\mathbf{z}_i$ and $\mathbf{z}_j$ are highly similar. The GRAMMAR-Gamma cross-term approximation $\mathbf{z}_j^T\mathbf{V}^{-1}\mathbf{z}_i \approx \bar{c}\,\mathbf{z}_j^T\mathbf{z}_i = \bar{c}\,N\,r_{ij}$ therefore performs similarly well as the diagonal approximation $\mathbf{z}_l^T\mathbf{V}^{-1}\mathbf{z}_l \approx \bar{c}\,N$ (Section S3.2).

Substituting both the diagonal ($\bar{c}N$) and cross-term ($\bar{c}Nr_{ij}$) into the general formula:

$$E[\Delta\ell_{i\to j}^{\text{full}}] \approx \frac{\beta_i^2}{2}\left[\bar{c}N - \frac{(\bar{c}Nr_{ij})^2}{\bar{c}N}\right] = \frac{\beta_i^2}{2}\,\bar{c}N\,(1 - r_{ij}^2)$$

Therefore:

$$\boxed{E[\Delta\ell_{i\to j}^{\text{full}}] \approx \frac{\bar{c}\,N\,\beta_i^2}{2}(1 - r_{ij}^2)}$$

Since $f(d) = d/(dh^2 + 1 - h^2)$ is concave in $d$ (Section S7.1), Jensen's inequality gives $\bar{c} \leq \frac{M}{N}\,f(N/M) = \frac{1}{Nh^2/M + 1 - h^2}$ for $N > M$. The denominator exceeds 1, so $\bar{c} < 1$. In this regime, the full-GRM evidence is therefore less than simple regression and typically less than LOCO.

Moreover, $\bar{c}N = S(N)/h^2$ (Equation 6; Section S4.4) practically saturates at $M_e/h^2$ for $N \gg N_A^*$ (Section S7.5), so the full-GRM fine-mapping evidence has a finite ceiling:

$$E[\Delta\ell_{i\to j}^{\text{full}}] \xrightarrow{N\to\infty} \frac{M_e\,\beta_i^2}{2h^2}(1 - r_{ij}^2)$$

In contrast, the LOCO and simple regression evidence grow without bound in $N$.

**S12.4. Comparison** The three fine-mapping frameworks yield the following expected evidence:

| Framework | $E[\Delta\ell_{i\to j}]$ | Effective coefficient |
|---|---|---|
| Simple regression | $\frac{N\beta_i^2}{2}(1 - r_{ij}^2)$ | 1 |
| LOCO mixed model | $\frac{c_{\text{LOCO}}\,N\,\beta_i^2}{2}(1 - r_{ij}^2)$ | $c_{\text{LOCO}} \leq 1/(1 - h_{\text{LOCO}}^2)$; Section S5.2 |
| Full-GRM mixed model | $\frac{\bar{c}\,N\,\beta_i^2}{2}(1 - r_{ij}^2)$ | $\bar{c} < 1$ (for $N > M$) |

All three frameworks (simple regression, LOCO, and full-GRM mixed model) share the same $(1 - r_{ij}^2)$ factor governing within-block resolution, shaped by the population's recombination history $(1 - r_{ij}^2 \approx 4N_ed/(1 + 4N_ed))$. The frameworks differ only in the effective coefficient. The full-GRM coefficient $\bar{c}N$ saturates as $N \to \infty$ (Section S8), while simple regression ($N$) and LOCO ($c_{\text{LOCO}}N$) grow without bound. Whether LOCO improves over simple regression depends on the population structure and the ratio $N/M_{\text{LOCO}}$ (Section S5.2).

### S13. Fine-Mapping Resolution Regimes

We use $\beta_i^2$ (PVE) as the primary parameterization and 10 kb ($d \approx 10^{-4}$ Morgans) as the representative distance for fine-mapping resolution. At this scale, a candidate region contains tens of variants, consistent with the goal of localizing causal variants. For livestock species: $4N_ed \approx 0.04$ ($N_e \approx 100$, cattle) to 0.02 ($N_e \approx 50$, pigs/chickens), giving $1 - r_{ij}^2 \approx 0.038$–$0.020$.

The examples below use the simple regression formula $E[\Delta\ell] = N\beta_i^2(1 - r_{ij}^2)/2$ (Section S11.2). The LOCO mixed model shares the same form with coefficient $c_{\text{LOCO}}$ (Section S12.2), which changes the magnitude but not the qualitative conclusions.

**S13.1. Large-Effect QTL Under Simple Regression** Consider a SNP explaining $q = 1\%$ of phenotypic variance ($\beta_i^2 = 0.01$) at 10 kb ($d \approx 10^{-4}$ Morgans). The simple-regression $E[\Delta\ell]$ (Section S11.2) is tabulated below.

| Species | $N_e$ | $4N_e d$ | $1 - r_{ij}^2$ | $E[\Delta\ell]$ per unit $N$ |
|---|---|---|---|---|
| Cattle | 100 | 0.04 | 0.038 | $1.9 \times 10^{-4}\,N$ |
| Pig/chicken | 50 | 0.02 | 0.020 | $1.0 \times 10^{-4}\,N$ |
| Human | 10,000 | 4.0 | 0.80 | $4.0 \times 10^{-3}\,N$ |

For cattle at $N = 100{,}000$: $E[\Delta\ell] = 19$ (decisive). For pig/chicken at $N = 100{,}000$: $E[\Delta\ell] = 10$ (strong). For humans at $N = 100{,}000$: $E[\Delta\ell] = 400$ (trivially resolved). Large-effect QTL ($q \sim 1\%$) are fine-mappable to 10 kb resolution at achievable sample sizes across all species, but the required $N$ differs by an order of magnitude between livestock and humans.

**S13.2. Polygenic SNP Under Simple Regression** Under the diffuse polygenic model, a causal variant out of 50,000 has effect size $\beta_i^2 = h^2/50{,}000$. For $h^2 = 0.3$: $\beta_i^2 = 6 \times 10^{-6}$. The simple-regression $E[\Delta\ell]$ (Section S11.2) is tabulated below.

| Species | $N_e$ | $1 - r_{ij}^2$ (10 kb) | $E[\Delta\ell]$ per unit $N$ |
|---|---|---|---|
| Cattle | 100 | 0.038 | $1.1 \times 10^{-7}\,N$ |
| Pig/chicken | 50 | 0.020 | $6.0 \times 10^{-8}\,N$ |
| Human | 10,000 | 0.80 | $2.4 \times 10^{-6}\,N$ |

For cattle at $N = 1{,}000{,}000$: $E[\Delta\ell] = 0.11$ (negligible). For humans at $N = 1{,}000{,}000$: $E[\Delta\ell] = 2.4$ (marginal). These results suggest that polygenic effects are difficult to resolve to 10 kb resolution with genotype-phenotype associations alone at achievable sample sizes.

**S13.3. The Simple-Regression Resolution Threshold** For fine-mapping to distinguish the causal SNP $i$ from an alternative $j$, we require $E[\Delta\ell_{i\to j}] > \theta$, where $\theta$ is a decision threshold (e.g., $\theta = 3$ for a Bayes factor of $\approx 20$). This gives:

$$\boxed{\beta_i^2 > \frac{2\theta}{N} \cdot \frac{1 + 4N_e d}{4N_e d}}$$

or equivalently, the minimum sample size for resolution:

$$\boxed{N > \frac{2\theta}{\beta_i^2} \cdot \frac{1 + 4N_e d}{4N_e d}}$$

**Generality of the two-SNP result.** Although the derivation considers a single pair $(i, j)$, the result is fully general:

1. **One causal SNP among many candidates.** The hardest alternative to exclude is the one with the highest $r_{ij}^2$. The resolution is governed by the closest neighbor, exactly the pairwise formula.

2. **Multiple causal variants in a region.** Fine-mapping becomes strictly harder, never easier. The pairwise formula provides an **upper bound** on resolution.

3. **Tiny-effect SNPs in a block are indistinguishable from each other.** If all SNPs in a block have similar small effects ($\beta_i^2 \approx h^2/M$), $E[\Delta\ell_{i\to j}]$ between any pair is negligible.

**S13.4. The Practical Full-GRM Effect-Size Floor** The simple-regression formula (Section S13.3) describes an idealized scenario in which polygenic confounding is set aside. Under full-GRM mixed-model fine-mapping, polygenic effects are explicitly modeled, and the expected log-likelihood difference saturates as $N$ grows (Section S12.3): $E[\Delta\ell_{i\to j}^{\text{full}}] \to M_e\beta_i^2(1 - r_{ij}^2)/(2h^2)$ in the practical regime ($N_A^* \ll N \ll N_B^*$). Setting this $> \theta$ gives a fine-mapping threshold that depends only on $N_e$, $L$, $h^2$, and the inter-marker distance $d$:

$$\boxed{\beta_i^2 > \frac{2\theta h^2}{M_e\,(1 - r_{ij}^2)}}$$

Substituting $M_e = 4N_eL$ and $1 - r^2 = 4N_ed/(1 + 4N_ed)$:

$$\boxed{\beta_i^2 > \frac{\theta h^2\,(1 + 4N_e d)}{8N_e^2 L\, d}}$$

Unlike the simple-regression threshold of Section S13.3, this threshold does not decrease with $N$: increasing the sample size approaches the asymptote but cannot practically lower this floor.

**When the floor is binding.** The full-GRM coefficient $\bar{c}\,N = S(N)/h^2$ (Equation 6; Section S4.4) practically saturates at $M_e/h^2$ for $N \gg N_A^*$ (Section S7.5). For sample sizes well below the transition scale ($N \ll N_A^*$), the full-GRM fine-mapping evidence is rising with $N$. For sample sizes well above the transition scale ($N \gg N_A^*$), the evidence has plateaued and the floor above is the governing constraint. For livestock species at currently feasible sample sizes ($N \sim 10^5$), $N_A^*$ is in the range of a few $\times 10^3$ to $10^5$, so $N$ can exceed the saturation point and the full-GRM floor can be binding. Quantitative thresholds for cattle, pig, chicken, and human are in Section S15.

### S14. What LOCO Detects: Block-Level vs. SNP-Level Signals

**S14.1. LOCO Detects Block-Level Signals** Under LOCO, the focal chromosome's genetic variance $h^2_{\text{chr}}$ is unmodeled and absorbed into the residual. The LOCO NCP for focal SNP $l$ is (from Section S5.3):

$$\text{NCP}_{\text{LOCO}} = c_{\text{LOCO}}\, N \cdot \frac{h^2_{\text{chr}}}{M_{\text{chr}}} \cdot \sum_s r^2_{ls} W_{ss}$$

with $c_{\text{LOCO}}$ as defined in Section S5.2. Under a weak independence assumption (that $r^2_{ls}$ and $W_{ss}$ are approximately uncorrelated across SNPs on the focal chromosome), this reduces to:

$$\text{NCP}_{\text{LOCO}} \approx c_{\text{LOCO}}\, N \cdot \frac{h^2_{\text{chr}}}{M_{\text{chr}}} \cdot R_l$$

where $R_l = \sum_s r^2_{ls}$ is the LD score of the focal SNP. The key step is:

$$\frac{1}{M_{\text{chr}}} \sum_s r^2_{ls} W_{ss} \approx \left(\frac{\sum_s r^2_{ls}}{M_{\text{chr}}}\right) \left(\frac{\sum_s W_{ss}}{M_{\text{chr}}}\right) = \frac{R_l}{M_{\text{chr}}} \cdot 1$$

since $\sum_s W_{ss}/M_{\text{chr}} = 1$ by normalization. This assumption holds whenever large-effect QTL are not systematically concentrated in high-LD or low-LD regions. For a SNP in a region of strong LD (large $R_l$), the LOCO NCP can be substantial even without any detectable QTL.

**S14.2. The Full-GRM Correctly Filters Block-Level Signals** Under the full-GRM analysis, the NCP at practical saturation ($N^*_A \ll N \ll N^*_B$) is:

$$\text{NCP}^{N\to\infty}_{\text{full}} = \frac{M_e}{M} \sum_s r^2_{ls} \tilde{W}_{ss}$$

For diffuse polygenic effects ($W_{ss} \approx 1$), $\tilde{W}_{ss} \approx 0$ and the NCP is negligible.

**S14.3. Three Classes of GWAS Signals** This analysis reveals three classes of associations:

**Class 1: Large-effect QTL** ($\tilde{W}_{ss} \gg 0$).

- Detected by both full-GRM and LOCO.
- Fine-mappable: $E[\Delta\ell] \propto N W_{ss} h^2 (1 - r^2_{ij})/M$ is large because $W_{ss} \gg 1$.
- Biologically actionable.

**Class 2: Polygenic block signals** ($W_{ss} \approx 1$ for all $s$, but $R_l$ is large).

- Detected by LOCO at large $N$ (NCP $\propto N R_l$, grows without bound).
- **Not detected** by full-GRM (NCP $\approx 0$).
- **Not fine-mappable**: $E[\Delta\ell] \propto N h^2 (1 - r^2_{ij})/M$ is negligible within blocks because $W_{ss} \approx 1$.
- Statistically real but uninformative.

**Class 3: Moderate-effect QTL** ($\tilde{W}_{ss}$ small but positive).

- Detected by LOCO; in a shorthand best-case criterion, detected by full-GRM only if $\tilde{W}_{ss} \cdot M_e/M > 30$ (the NCP threshold for 50% power at $p < 5 \times 10^{-8}$; Section S8).
- Marginally fine-mappable: requires very large $N$ to resolve within blocks.

#### S15. Quantitative Predictions for Fine-Mapping

We use the following species-specific parameters throughout this section and in the main text:

| Species | $N_e$ | $L$ (Morgans) | $M_e = 4N_eL$ | Reference for $N_e$ | Reference for $L$ |
|---|---|---|---|---|---|
| Cattle | 100 | 25 | 10,000 | Lozada-Soto *et al.* 2022 | Wang *et al.* 2016 |
| Pig | 50 | 20 | 4,000 | Zanella *et al.* 2016 | Tortereau *et al.* 2012 |
| Chicken | 50 | 30 | 6,000 | Qanbari *et al.* 2010 | Groenen *et al.* 2009 |
| Human | 10,000 | 35 | 1,400,000 | Charlesworth 2009 | Kong *et al.* 2002 |

The $N_e$ values represent recent effective population sizes in commercial breeding populations. The genome lengths $L$ are sex-averaged autosomal genetic map lengths: ∼2,500 cM for cattle (Wang *et al.* 2016), ∼2,000 cM for pig (Tortereau *et al.* 2012), ∼3,000 cM for chicken (Groenen *et al.* 2009), and ∼3,500 cM for human (Kong *et al.* 2002). Chicken and pig share a similar $N_e$ but differ in genome length, yielding different $M_e$.

The minimum PVE for fine-mapping resolution depends on which framework is used. The simple-regression threshold (Section S13.3) is $\beta_i^2 > 2\theta/[N(1 - r_{ij}^2)]$, which decreases as $1/N$; the full-GRM threshold (Section S13.4) is $\beta_i^2 > 2\theta h^2/[M_e(1 - r_{ij}^2)]$, independent of $N$. Substituting $1 - r_{ij}^2 \approx 4N_ed/(1 + 4N_ed)$ gives species-specific thresholds. The table below uses $\theta = 3$ (Bayes factor $\approx 20$), $N = 100{,}000$ for the simple-regression column, and $h^2 = 0.3$ for the full-GRM column.

| Species | $N_e$ | $M_e$ | Distance | $d$ (Morgans) | $1 - r^2$ | Min PVE: simple regression ($N$=$10^5$) | Min PVE: full-GRM method ($h^2$=0.3) |
|---|---|---|---|---|---|---|---|
| Cattle | 100 | 10,000 | 10 kb | $10^{-4}$ | 0.038 | 0.16% | 0.47% |
| Cattle | 100 | 10,000 | 100 kb | $10^{-3}$ | 0.29 | 0.021% | 0.062% |
| Pig | 50 | 4,000 | 10 kb | $10^{-4}$ | 0.020 | 0.30% | 2.25% |
| Pig | 50 | 4,000 | 100 kb | $10^{-3}$ | 0.17 | 0.036% | 0.27% |
| Chicken | 50 | 6,000 | 10 kb | $10^{-4}$ | 0.020 | 0.30% | 1.5% |
| Chicken | 50 | 6,000 | 100 kb | $10^{-3}$ | 0.17 | 0.036% | 0.18% |
| Human | 10,000 | 1,400,000 | 10 kb | $10^{-4}$ | 0.80 | 0.0075% | $1.6 \times 10^{-4}$% |
| Human | 10,000 | 1,400,000 | 100 kb | $10^{-3}$ | 0.98 | 0.0061% | $1.3 \times 10^{-4}$% |

The simple-regression column depends on $N_e$ (through LD decay) but not on $L$, so pig and chicken share identical values in that column; the full-GRM column depends on $M_e = 4N_eL$ and so distinguishes pig from chicken through their different $L$.

For comparison, the full-GRM mixed-model detection threshold at the practical NCP ceiling is $q_{\min} \approx 30h^2/M_e$ (Section S8):

| Species | $N_e$ | $M_e = 4N_eL$ | $q_{\min}$ ($h^2 = 0.3$) |
|---|---|---|---|
| Cattle | 100 | 10,000 | 0.09% |
| Pig | 50 | 4,000 | 0.23% |
| Chicken | 50 | 6,000 | 0.15% |
| Human | 10,000 | 1,400,000 | $6.4 \times 10^{-4}$% |

In livestock species, variants whose effects are too small to be detected at the practical full-GRM detection floor ($q < q_{\min}$) are generally not fine-mappable to a reasonably high resolution (e.g., tens of kbp) under either fine-mapping framework. For cattle, the detection threshold is $q_{\min} \approx 0.09\%$, while the minimum PVE for 10 kb fine-mapping resolution is 0.16% under simple regression at $N = 100{,}000$ (already exceeding the detection threshold) and 0.47% under the full-GRM method. For pig (detection $q_{\min} \approx 0.23\%$; simple-regression 10 kb threshold 0.30%; full-GRM floor 2.25%) and chicken (detection $q_{\min} \approx 0.15\%$; simple-regression 10 kb threshold 0.30%; full-GRM floor 1.5%), the same pattern holds: in all three livestock species the full-GRM floor is the most restrictive of the three. In humans ($N_e \approx 10{,}000$), both fine-mapping thresholds and the detection threshold are orders of magnitude smaller than those of livestock, and the full-GRM floor is the loosest of the three because $M_e/h^2$ is much larger than achievable $N$.

This establishes a unified effect-size floor for genotype–phenotype association: in livestock, the same population parameter $N_e$ simultaneously constrains detection power through the practical full-GRM ceiling ($M_e = 4N_eL$) and fine-mapping resolution through $4N_e$-scaled LD decay.

---

### Summary of Key Results

#### Part I: NCP of the mixed-model association test

| Quantity | Expression | Reference |
|---|---|---|
| Per-SNP NCP | $\frac{c_l \cdot N \cdot h^2}{M} \cdot \sum_s r_{ls}^2 \tilde{W}_{ss}$ | Equation 1; S4.1 |
| Per-SNP coefficient | $c_l = \mathbf{z}_l^T \mathbf{V}^{-1} \mathbf{z}_l \,/\, N$ | Equation 2; S4.1 |
| Eigenvalue expansion of $c_l$ | $c_l = \frac{1}{N} \sum_k \frac{a_{lk}^2}{d_k h^2 + 1 - h^2}$ | Equation 3; S4.2 |
| Average coefficient ($d_k$ form) | $\bar{c} = \frac{1}{N} \sum_k \frac{d_k}{d_k h^2 + 1 - h^2}$ | Equation 4; S4.3 |
| NCP ($\ell_k$ form, $\bar{c}$ approx.) | $\frac{S(N)}{M} \sum_s r_{ls}^2 \tilde{W}_{ss}$ | Equation 5; S4.4 |
| Sigmoid sum | $S(N) = \sum_k \frac{N\ell_k}{N\ell_k + M\lambda} = \bar{c}\, N\, h^2$ | Equation 6; S4.4 |
| NCP (ideal tagging case) | $S(N)\,(q/h^2 - 1/M)$ | Equation 7; S4.4 |
| LOCO coefficient | $c_{\text{LOCO}} \leq 1/(1 - h_{\text{LOCO}}^2)$; depends on $N/M_{\text{LOCO}}$ and pedigree | Equation 12; S5.2 |
| LOCO NCP | $c_{\text{LOCO}}\, N \cdot h_{\text{chr}}^2 / M_{\text{chr}} \cdot \sum_s r_{ls}^2 W_{ss}$ | Equation 13; S5.3 |

#### Part II: Properties of the sigmoid sum and practical saturation

| Quantity | Expression | Reference |
|---|---|---|
| Per-mode transition scale | $N_k^* = M\lambda/\ell_k$ | S4.4 |
| Group A transition scale | $N_A^* = M_e\lambda/\rho$ | S7.4 |
| Group B transition scale | $N_B^* = M_B\lambda/(1-\rho)$ | S7.4 |
| Small-$N$ NCP ($N \ll N_A^*$) | $\frac{Nh^2}{M(1-h^2)} \sum_s r_{ls}^2 \tilde{W}_{ss}$ | Equation 8; S7.5 |
| Practical-ceiling NCP ($N_A^* \ll N \ll N_B^*$) | $\frac{M_e}{M} \sum_s r_{ls}^2 \tilde{W}_{ss}$ | Equation 9; S8 |
| Min detectable PVE | $q_{\min} \approx 30h^2/M_e$ | Equation 10; S8 |
| DRP $N$-equivalence | $N_{\text{equiv}} = N_{\text{DRP}} \cdot r^2(1 - h_{\text{orig}}^2)/[h_{\text{orig}}^2(1 - r^2)]$ | Equation 11; S9.2 |

#### Part III: Genomic prediction

| Quantity | Expression | Reference |
|---|---|---|
| In-sample GBLUP reliability | $\bar{R}^2_{\text{in}} = 1 - \lambda S(N)/N$ | Equation 17; S10.4 |
| In-sample GBLUP reliability ($N^*_A \ll N \ll N^*_B$) | $\bar{R}^2_{\text{in}} \approx \frac{Nh^2}{Nh^2+M_e(1-h^2)}$ | Equation 18; S10.4 |
| Out-of-sample (LOO) reliability | $1-(\lambda+1)\frac{S(N)}{N} \leq \bar{R}^2_{\text{LOO}} \leq 1-\frac{\lambda S(N)/N}{1-S(N)/N}$ | Equation 19; S10.6 |
| In-sample optimism | $\lambda\frac{(S(N)/N)^2}{1-S(N)/N} \leq \Delta_{\text{opt}} \leq \frac{S(N)}{N}$ | S10.6 |
| Daetwyler limit ($N \ll M_e$) | $r^2_{\text{Daetwyler}} = \frac{Nh^2}{Nh^2+M_e} \approx \frac{Nh^2}{M_e}$ | S10.7 |

**Part IV: Fine-mapping resolution limit**

| Quantity | Expression | Reference |
|---|---|---|
| Fine-mapping evidence (simple regression) | $\frac{N\beta_i^2}{2}(1-r^2_{ij})$ | Equation 14; S11.2 |
| LD decay | $1-r^2_{ij} \approx \frac{4N_e d}{1+4N_e d}$ | Equation 15; S1.5 |
| LOCO fine-mapping evidence | $\frac{c_{\text{LOCO}}\,N\,\beta_i^2}{2}(1-r^2_{ij})$ | S12.2 |
| Full-GRM fine-mapping evidence | $\frac{\bar{c}\,N\,\beta_i^2}{2}(1-r^2_{ij})$ | S12.3 |
| Min PVE for resolution (simple regression) | $\beta_i^2 > \frac{2\theta}{N} \cdot \frac{1+4N_e d}{4N_e d}$ | S13.3 |
| Min PVE for resolution (full-GRM) | $\beta_i^2 > \frac{2\theta h^2}{M_e(1-r^2_{ij})}$ | Equation 16; S13.4 |

### References


Charlesworth B. 2009. Fundamental concepts in genetics: effective population size and patterns of molecular evolution and variation. Nature Reviews Genetics. 10:195–205. doi:10.1038/nrg2526.

Cheng H, Garrick DJ, Fernando RL. 2017. Efficient strategies for leave-one-out cross validation for genomic best linear unbiased prediction. Journal of Animal Science and Biotechnology. 8:38. doi:10.1186/s40104-017-0164-6.

Daetwyler HD, Pong-Wong R, Villanueva B, Woolliams JA. 2010. The impact of genetic architecture on genome-wide evaluation methods. Genetics. 185(3):1021–1031. doi:10.1534/genetics.110.116855.

Daetwyler HD, Villanueva B, Woolliams JA. 2008. Accuracy of predicting the genetic risk of disease using a genome-wide approach. PLoS ONE. 3:e3395. doi:10.1371/journal.pone.0003395.

Garrick DJ, Taylor JF, Fernando RL. 2009. Deregressing estimated breeding values and weighting information for genomic regression analyses. Genetics Selection Evolution. 41:55. doi:10.1186/1297-9686-41-55.

Groenen MAM, Wahlberg P, Foglio M, Cheng HH, Megens H-J, Crooijmans RPMA, Besnier F, Lathrop M, Muir WM, Wong GK-S, Gut I, Andersson L. 2009. A high-density SNP-based linkage map of the chicken genome reveals sequence features correlated with recombination rate. Genome Research. 19:510–519. doi:10.1101/gr.086538.108.

Hastie T, Tibshirani R, Friedman J. 2009. The elements of statistical learning: data mining, inference, and prediction. 2nd ed. New York (NY): Springer.

Kong A, Gudbjartsson DF, Sainz J, Jonsdottir GM, Gudjonsson SA, Richardsson B, Sigurdardottir S, Barnard J, Hallbeck B, Masson G, Shlien A, Palsson ST, Frigge ML, Thorgeirsson TE, Gulcher JR, Stefansson K. 2002. A high-resolution recombination map of the human genome. Nature Genetics. 31:241–247. doi:10.1038/ng917.

Lozada-Soto EA, Tiezzi F, Jiang J, Cole JB, VanRaden PM, Maltecca C. 2022. Genomic characterization of autozygosity and recent inbreeding trends in all major breeds of US dairy cattle. Journal of Dairy Science. 105:8956–8971. doi:10.3168/jds.2022-22116.

Pocrnic I, Lourenco DAL, Masuda Y, Legarra A, Misztal I. 2016. The dimensionality of genomic information and its effect on genomic prediction. Genetics. 203:573–581. doi:10.1534/genetics.116.187013.

Qanbari S, Hansen M, Weigend S, Preisinger R, Simianer H. 2010. Linkage disequilibrium reveals different demographic history in egg laying chickens. BMC Genetics. 11:103. doi:10.1186/1471-2156-11-103.

Stam P. 1980. The distribution of the fraction of the genome identical by descent in finite random mating populations. Genetics Research. 35:131–155.

Sved JA. 1971. Linkage disequilibrium and homozygosity of chromosome segments in finite populations. Theoretical Population Biology. 2:125–141. doi:10.1016/0040-5809(71)90011-6.

Tortereau F, Servin B, Frantz L, Megens H-J, Milan D, Rohrer G, Wiedmann R, Beever J, Archibald AL, Schook LB, Groenen MAM. 2012. A high density recombination map of the pig reveals a correlation between sex-specific recombination and GC content. BMC Genomics. 13:586. doi:10.1186/1471-2164-13-586.

VanRaden PM. 2008. Efficient methods to compute genomic predictions. Journal of Dairy Science. 91:4414–4423. doi:10.3168/jds.2007-0980.

VanRaden PM, Tooker ME, O'Connell JR, Cole JB, Bickhart DM. 2017. Selecting sequence variants to improve genomic predictions for dairy cattle. Genetics Selection Evolution. 49:32. doi:10.1186/s12711-017-0307-4.

Wang J, Gao Y, Toghiani S, Cole JB, Maltecca C, Ma L, Jiang J. 2025. Genome-wide association study and fine-mapping using imputed sequences to prioritize candidate genes for 30 complex traits in 50,309 Holstein bulls. Journal of Dairy Science. 108(12):12506–12518. doi:10.3168/jds.2025-27058.

Wang J, Tiezzi F, Huang Y, See G, Schwab C, Wei J, Maltecca C, Jiang J. 2025. Fine-mapping methods for complex traits: essential adaptations for samples of related individuals. Briefings in Bioinformatics. 26(6):bbaf614. doi:10.1093/bib/bbaf614.

Wang Z, Shen B, Jiang J, Li J, Ma L. 2016. Effect of sex, age and genetics on crossover interference in cattle. Scientific Reports. 6:37698. doi:10.1038/srep37698.

Zanella R, Peixoto JO, Cardoso FF, Cardoso LL, Biegelmeyer P, Cantão ME, Otaviano A, Freitas MS, Caetano AR, Ledur MC. 2016. Genetic diversity analysis of two commercial breeds of pigs using genomic and pedigree data. Genetics Selection Evolution. 48:24. doi:10.1186/s12711-016-0203-3.
